# Antibiotic-induced protein mislocalization drives hypervesiculation in *Salmonella*

**DOI:** 10.64898/2026.11.03.764146

**Authors:** Yongjun Son, Jihye Yang, Wonjae Kim, Jinwon Park, Hyosung Choi, Hyeongseop Jeong, Woojun Park

## Abstract

The incorporation of cytosolic proteins into bacterial outer membrane vesicles (OMVs) has remained mechanistically unresolved, as these proteins are normally excluded from vesiculation sites by the inner membrane (IM) barrier. Here, we show that antibiotic-induced envelope perturbation compromises this spatial barrier, enabling an aberrant interaction between proteins that are ordinarily segregated across cellular compartments and thereby promoting MV formation in *Salmonella.* Exposure to non-lethal cephalexin (LEX) induced robust hypervesiculation, accompanied by inhibition of the essential division penicillin-binding protein (PBP) 3 and accumulation of the PBP5. Under envelope-perturbing conditions, PBP5 accumulated at lateral sites associated with MV emergence and promoted periplasmic crowding independently of its catalytic activity. A cytosolic acyl carrier protein (ACP) emerged as a PBP5-interacting partner under LEX exposure, coinciding with cryogenic electron microscopy–detected IM perturbation and aberrant ACP mislocalization. Formation of the PBP5–ACP complex promoted PBP5 accumulation, resulting in periplasmic crowding, progressive weakening of outer membrane–peptidoglycan tethering, and ultimately MV formation. Together, these findings uncover a mechanism whereby stress-induced disruption of subcellular compartmentalization enables a non-canonical interaction between normally segregated proteins, driving hypervesiculation in *Salmonella*.

## Introduction

Gram-negative bacteria constitutively release membrane vesicles (MVs), a process that is often enhanced under stress conditions such as antibiotic exposure, heat shock, and oxidative stress (1,2). Among the different types of MVs, outer membrane vesicles (OMVs) have been the most extensively studied in terms of their biogenesis (3). Representative models include the weakening of outer membrane (OM)–peptidoglycan (PG) tethering through reduced OmpA- or Lpp-mediated connections, and periplasmic crowding arising from the accumulation of unfolded proteins or other periplasmic material, both of which can promote local envelope remodeling and vesiculation (4–6). In addition, lipid-driven OM curvature can stimulate OMV production through phospholipid accumulation in the outer leaflet caused by defects in the Mla (VacJ/Yrb) transport pathway or through the insertion of amphipathic molecules such as the *Pseudomona*s quinolone signal into the OM (7,8). In addition to conventional OMVs, Gram-negative bacteria can also naturally produce outer-inner membrane vesicles (O-IMVs) containing both outer and inner membranes (IM) and cytoplasmic components (9,10). One proposed mechanism involves transient autolysin-mediated PG degradation, which may facilitate the protrusion of both membranes and subsequent vesicle release (1,11). MVs function as versatile delivery vehicles that transport a broad range of biomolecules. These nanosized vesicles contain OM-derived components (e.g., OmpA and Lpp), periplasmic proteins (e.g., Tol-Pal and LolA), and other cargo, including DNA, quorum-sensing molecules, and virulence factors, thereby contributing to diverse microbe–host interactions such as adhesion, biofilm formation, immune evasion, and toxin delivery (1,2,6). Intriguingly, cytoplasmic proteins, including glyceraldehyde 3-phosphate dehydrogenase (GAPDH), elongation factor Tu (EF-Tu), GroEL, and ribosomal proteins, are frequently detected in non-lytic MVs (12–14). Because these proteins are normally segregated from vesiculation sites by the inner membrane (IM), their recurrent detection remains difficult to reconcile with current models of OMV biogenesis. This phenomenon cannot be readily explained as a simple consequence of cell lysis and instead suggests that cytoplasmic cargo incorporation may reflect an unrecognized aspect of vesicle formation. However, the mechanism by which cytoplasmic proteins gain access to OMVs, and whether this process contributes directly to OMV biogenesis, remains unknown.

Enteric *Salmonella enterica* serovar Typhimurium is a major cause of foodborne disease worldwide, accounting for millions of infections and substantial mortality each year (15). As a facultative intracellular pathogen capable of surviving and replicating within macrophages inside the *Salmonella*-containing vacuole (SCV), *Salmonella* poses a significant public health burden and has become an important model for investigating bacterial stress adaptation (16). Because β-lactam antibiotics are widely used to control *Salmonella* infections by targeting PG synthesis, the cellular consequences of β-lactam exposure have been extensively studied (17–19). Previous studies have reported several envelope-associated responses to β-lactam stress, including filamentation, altered PG remodeling, and increased OMV production (17–19). During a screen of β-lactam antibiotics, we observed robust non-lytic hypervesiculation in *Salmonella* exposed to sub-lethal concentrations of cephalexin (LEX). Investigation of this phenotype revealed a previously unrecognized interaction between the penicillin-binding protein (PBP) 5 and the cytosolic acyl carrier protein (ACP) under LEX stress. Mechanistically, localized perturbation of inner membrane (IM) integrity was associated with aberrant ACP localization and formation of a non-canonical PBP5–ACP complex, leading to periplasmic crowding and enhanced vesiculation. These findings identify a mechanism by which antibiotic-induced disruption of subcellular compartmentalization enables interactions between normally segregated proteins and provide insight into the long-standing observation of cytoplasmic protein incorporation into non-lytic MVs.

## Results

### Non-lytic hypervesiculation in *S.* Typhimurium under LEX stress

We first examined cell viability and MV production in *S.* Typhimurium exposed to subinhibitory concentrations of β-lactam antibiotics **(Fig. 1a–c)**. Based on our minimum inhibitory concentration (MIC) test, short-term killing curve assays showed more than ∼95% of cells remained viable during the initial 60-min exposure at 1/4×MIC of each β-lactam antibiotic **(Table S1, Fig. S1)**. Based on these MIC and viability results, 1/4×MIC was selected to compare growth and MV production across the β-lactam antibiotics tested. Although growth profiles were broadly comparable among the β-lactam-treated conditions, MV production varied markedly depending on the antibiotic used **(Fig. 1a–c)**. LEX-treated cells exhibited the highest MV production, showing a 5.0-fold increase relative to the untreated control **(Fig. 1c)**. To examine whether the elevated MV production under LEX treatment was associated with increased cell death, the proportion of propidium iodide (PI)-stained cells was quantified by flow cytometry after 6 h of antibiotic exposure. LEX-treated cells showed a relatively low proportion of PI-positive cells (6.1%), whereas ceftriaxone (CRO)-treated cells exhibited the highest PI-positive proportion (12.0%) despite inducing only a modest increase in MV production (1.4-fold relative to the untreated control) **(Fig. S2)**. These results suggest that the markedly increased non-lytic vesiculation under LEX stress was not directly associated with an increase in dead cells. Growth and MV production of *S.* Typhimurium were further examined across a range of LEX concentrations (0–256 µg/mL; 1/64 to 1× MIC) **(Fig. 1d, 1e)**. Among the concentrations tested, 1/4×MIC induced the highest MV production. A conditioned-medium re-inoculation assay further demonstrated that MV production continued to increase for up to 2 h in the absence of detectable cell death, to exclude the possibility that LEX-induced MVs originated from a minor dying subpopulation **(Fig. S3a-d)**.

**Figure 1.**
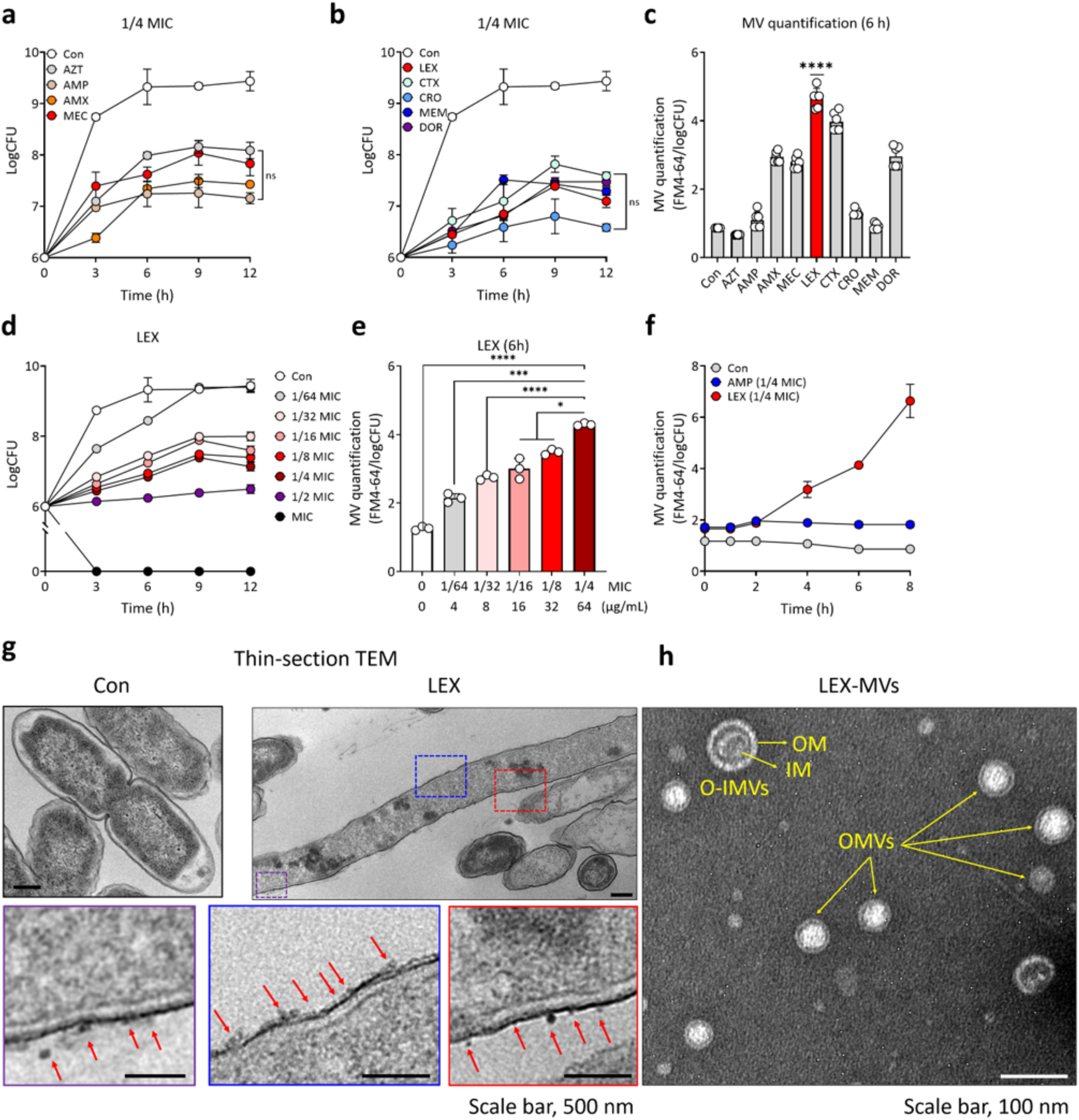
Measurements of the growth curves and MV formation in *S.* Typhimurium under β-lactam antibiotic stress. **a, b** Growth curves of *S.* Typhimurium under subinhibitory concentrations (1/4×MIC) of nine β-lactam antibiotics. Experiments were performed at 37°C under static conditions. Data represent three independent biological replicates (N=3). **c** Quantification of MV production at 6 h under subinhibitory concentrations (1/4×MIC) of each β-lactam antibiotic using the lipophilic dye FM4-64 (N=6). **d** Growth curves of *S.* Typhimurium under LEX stress (0-256 µg/mL). **e** MV quantification of *S.* Typhimurium at 6 h under LEX stress (0–64 µg/mL) using FM4-64. **f** Time-dependent MV level in *S.* Typhimurium treated with AMP (1/4×MIC, 16 µg/mL) and LEX (1/4×MIC, 64 µg/mL), respectively (N = 3). **g** Thin-section TEM images of cells cultured in LB or LB with LEX (1/4×MIC) for 6 h. **h** Negative-stain TEM images of MVs derived from LEX-treated cells. Statistical comparisons were performed using one-way ANOVA with Tukey’s multiple comparison test (*P < 0.05, ***P < 0.001, and ****P < 0.0001; ns, not significant).

To distinguish whether MV production under LEX stress occurs progressively due to accumulation of periplasmic components or arises as a sudden event, time-dependent MV formation was monitored in wild-type (WT) strain cells until the stationary phase (∼8 h) **(Fig. 1f)**. In LEX-treated cells, MV formation was initiated at 4 h and continued to increase until the stationary phase. In contrast, MV levels remained unchanged throughout the growth in both ampicillin (AMP)-treated and untreated cells **(Fig. 1f)**. These findings suggest that MV production in LEX-treated cells might be associated with progressive periplasmic crowding over time. Our field-emission scanning electron microscopy (FE-SEM) and negative-stain transmission electron microscopy (TEM) analyses revealed that LEX-treated cells were markedly elongated compared with non-treated cells **(Fig. S4a-d)**. Notably, numerous MVs were distributed across the entire cell surface, rather than being restricted to specific regions such as the septum or poles **(red arrows, Fig. S4b, S4d)**. Thin-section TEM further captured extensive lateral membrane blebbing, with large numbers of MVs appearing to bud from the lateral cell envelope in LEX-treated cells **(red arrows, Fig. 1g)**. To distinguish the types of MVs produced from LEX-treated cells, the MVs were isolated from culture supernatants and purified by OptiPrep density gradient centrifugation **(Fig. S5a, S5b)**. Distinct whitish bands were observed in the 20–25% gradient fractions, particularly under LEX stress **(Fig. S5b)**. Western blot analyses of the collected fractions (20–35%) revealed that all four proteins representing different subcellular compartments, OmpA (OM), DsbA (periplasm), TatC (IM), and GAPDH (cytoplasm), were predominantly detected in the vesicle-enriched fractions, particularly those corresponding to the bands at 20–25% **(Fig. S5c)**. Our Proteomic analysis of the recovered vesicles further identified proteins associated with different subcellular compartments, including OM proteins (e.g., OmpA, Lpp, and LpoA), periplasmic proteins (e.g., SurA, DegP, and DsbA), IM proteins (e.g., PBP1a, TatA, and PBP5), and cytoplasmic proteins (e.g., EF-Tu, GroEL, and ACP) **(Table S2)**. Native TEM analysis of the pooled vesicle-containing fractions revealed two distinct morphological types of MVs, predominantly conventional OMVs (99.4%; 1044/1050), with a small proportion of double-membrane structures consistent with O-IMVs (0.6%; 6/1050) **(yellow arrows; Fig. 1h)**. Notably, the coexistence of predominantly conventional OMVs with a small population of O-IMVs is consistent with previous observations that both vesicle types can be produced in the absence of overt cell lysis (6,9,10,20). Moreover, MVs isolated from LEX-treated cells contained higher amounts of DNA and protein and exhibited a more uniform size distribution than those from untreated cells **(Fig. S5e–h).** Together, our findings indicate that two morphologically distinct types of MVs, predominantly conventional OMVs and a small proportion of O-IMVs, are produced under LEX stress.

### PBP3 inhibition and PBP5/6 upregulation under LEX stress

We performed a Bocillin fluorescent-labeling (Bocillin-FL) assay using a penicillin analog that selectively binds catalytically active PBPs to evaluate the profile of DD-transpeptidase activity under LEX stress **(Table S3, Fig. 2a)** (21,22). Our Bocillin-FL signals revealed that LEX-treated cells exhibited a marked reduction in PBP3 (∼0.3-fold) relative to non-treated cells, whereas the levels of PBP5 and PBP6 were strongly elevated (∼2.1-11.5-fold) **(Fig. 2a, 2b)**. If hypervesiculation results from a LEX-specific PBP pattern, the resulting Bocillin-FL profile would differ from that induced by other β-lactam antibiotics (AMP, CTX, and MEM) **(Fig. S6a–c)**. As expected, the DD-transpeptidase activity profiles of AMP- and MEM-treated cells showed distinct patterns from that of the LEX-treated cells **(Fig. S6a-b)**. In contrast, the DD-transpeptidase activity pattern of cells exposed to CTX (the second strongest MV-inducing β-lactam) was similar to that of the LEX-treated cells **(Fig. S6c)**. To further validate our Bocillin-FL assay, major PBP-levels (PBP2, PBP3, PBP5, PBP6a/b) were quantified by Western blot analyses using chromosomally FLAG-tagged PBP strains under LEX stress **(Fig. 2c)**. Consistent with our Bocillin-FL assay results, both PBP5 and PBP6a/6b were upregulated (1.2∼11.5-fold) under LEX stress relative to control **(Fig. 2c, 2d)**. PBP3 was barely detected at 2 h but sharply increased at 4 and 6 h following LEX treatment (1.7- and 2.0-fold, respectively) relative to the control condition **(Fig. 2c, 2d)**. Since Western blot assay can detect FLAG-tagged proteins regardless of their catalytic state, the increased PBP3 signal may reflect accumulation of both active and inactive forms (23). We hypothesized that impaired cell division under LEX stress may trigger a compensatory upregulation of *ftsI*, leading to the accumulation of dysfunctional PBP3. To test this hypothesis, the expression of genes encoding PBPs and muropeptide profiling were analyzed under LEX stress **(Fig. S7, S8)**. The expression of the *ftsI* gene remained unchanged at 2 h but was significantly upregulated (∼2.0-fold) at 4 h after exposure to LEX stress **(Fig. S7a)**. Our muropeptide profiling further supported that inactivation of PBP3 under LEX stress resulted in a marked reduction of 4,3-crosslinked muropeptides **(Fig. S8a-d)**. Consistent with our Bocillin-FL and Western blot results, the expression of genes encoding PBP5 and PBP6a/b was rapidly upregulated within 1 h of LEX exposure **(Fig. S7b)**. In parallel, muropeptide analyses revealed an increased proportion of monomeric tetra–muropeptides **(Table S4, Fig. S8a-d)**. Therefore, these results indicate that LEX-induced impairment of PBP3 activity results in the accumulation of dysfunctional PBP3, accompanied by the upregulation of PBP5 and PBP6.

**Figure 2.**
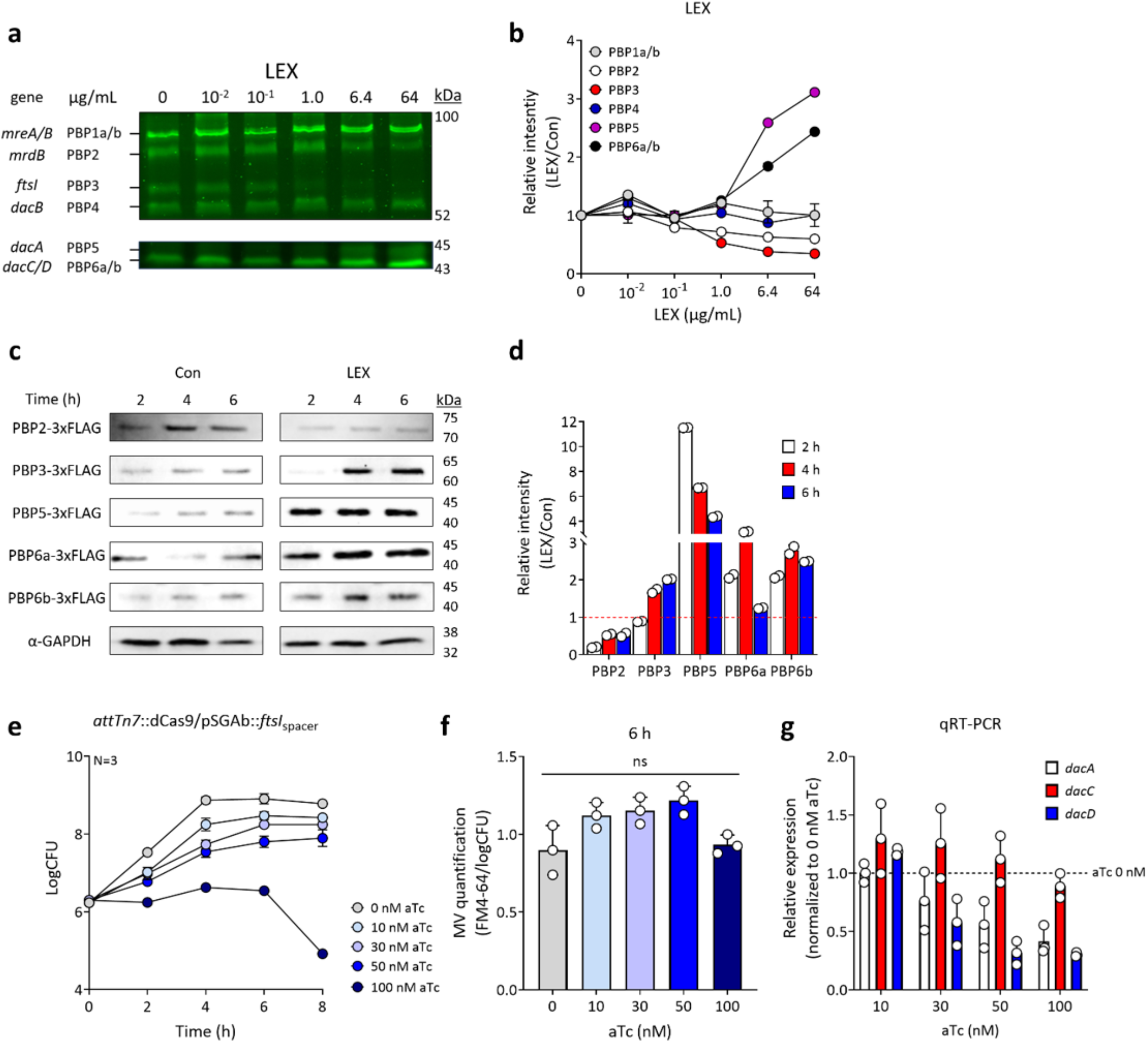
PBP profiling in *S.* Typhimurium under LEX stress. **a, b** WT cells were grown with concentration-dependent LEX (0-64 µg/mL) at 37 °C. Bocillin-FL–labeled PBPs were resolved by SDS–PAGE. The upper panels correspond to samples loaded with 300 µg/mL membrane protein, and the lower panels correspond to samples loaded with 600 µg/mL membrane protein. Band intensities of Bocillin-FL–labeled PBPs were quantified using ImageJ. Data represent three independent biological replicates (N=3). **c, d** PBP profiling of chromosomally 3×FLAG-tagged WT strains assessed by Western blotting. Cells were grown at 37 °C and treated with LEX (1/4×MIC). GAPDH was used as a loading control. Representative Western blotting images are shown. Band intensities were quantified using ImageJ, and protein levels are presented relative to the control. Crude extracts were normalized to 100 µg/mL total protein. Data represent two independent biological experiments performed in duplicate (N=2). **e, f** Quantification of growth curves and MV levels in *ftsI*^down^ strain (*attTn7*::dCas9/pSGAb::*ftsI*_spacer_) at different anhydrotetracycline (aTc) concentrations (0-100 nM) at 37 °C. Data represent three independent biological replicates (N=3). **g** Relative expression levels of genes encoding the PBP5, PBP6a, and PBP6b in *ftsI*^down^ strain. Data represent three independent biological replicates (N=3).

As LEX-mediated inhibition of PBP3 induces hypervesiculation in *S.* Typhimurium, we hypothesized that artificial repression of *ftsI* may similarly recapitulate these LEX-induced phenotypes in the absence of LEX stress **(Fig. 2e)**. We used CRISPR interference (CRISPRi)-based knockdown of *ftsI* gene by inducing the chromosomally integrated dCas9 using anhydrotetracycline (aTc) **(Fig. 2e–g)** (24). Increasing aTc concentrations progressively reduced growth, with a marked decrease observed at 50 nM aTc **(Fig. 2e)**. However, the amount of MVs did not increase after 6 h of aTc treatment in all tested strains **(Fig. 2f)**. Microscopic analyses revealed dose-dependent cell elongation, but the cells remained shorter than LEX-treated cells **(Fig. S9a, S9b)**. Additionally, the relative expression of the genes encoding the three monofunctional DD-carboxypeptidases was downregulated (∼0.5-fold) in the *ftsI* knockdown strain compared to the non-treated control **(Fig. 2g)**. Our results indicate that simply reducing the PBP3 levels could not mimic the LEX-driven accumulation of dysfunctional PBP3. Thus, our data suggest that the LEX-specific phenotypes, extreme filamentation and hypervesiculation, cannot be explained solely by reduced PBP3 abundance, but rather by the buildup of impaired PBP3.

### Essential role of PBP5/6 upregulation in hypervesiculation under LEX stress

To determine whether these DD-carboxypeptidases directly promote MV formation, individual knockout mutants of *dacA*, *dacC*, and *dacD* were constructed. All three knockout mutants exhibited the same MIC for LEX as the WT strain **(Fig. S10a)**. The growth curves of the mutant strains were comparable to that of the WT strain in LB and LB with LEX **(S10b, S10c)**. As expected, MV production in each mutant under LEX stress was markedly reduced (∼0.5-fold) at 6 h relative to the WT, whereas no significant difference in MV production was observed in LB **(Fig. 3a, S10d)**. Because deletion of the *dacA* gene resulted in the most severe MV-production defect under LEX stress, six *dacA* variants [the *dacA* gene knockout/complemented/overexpressing strains, the truncation (N/C-terminal region, respectively) mutants, and the activate site (K242A) mutant] were evaluated to identify the structural and functional elements of PBP5 required for MV biogenesis **(Fig. 3b, S10e-g)** (25). The growth of the WT and all six *dacA* variants showed no differences in LB or LB with LEX **(Fig. S10e, S10g)**. Additionally, MV levels showed no differences among the strains under control conditions **(Fig. S10f)**. Under LEX stress, among the WT, *dacA*-complemented, and *dacA*_OE_ strains produced comparable amounts of MV at 6 h **(Fig. 3b)**. In contrast, MVs were not detected in the *AdacA* strain under LEX stress throughout the growth period. Interestingly, the truncation mutants (*AdacA*_NT_ and *AdacA*_CT_) exhibited intermediate MV yields between the hypervesiculating WT strain and the MV-deficient *AdacA* mutant **(Fig. 3b)**. Given that the truncation mutants lack catalytic or anchoring regions, their reduced MV production likely reflects structural defects that impair PBP5 localization or accumulation within the periplasm, rather than a loss of D,D-carboxypeptidase activity. Supporting this interpretation, the catalytic-site mutant strain *dacA*_K242A_, which preserves the full-length structure and localization of PBP5, restored the MV production level similar to WT strain.

**Figure 3.**
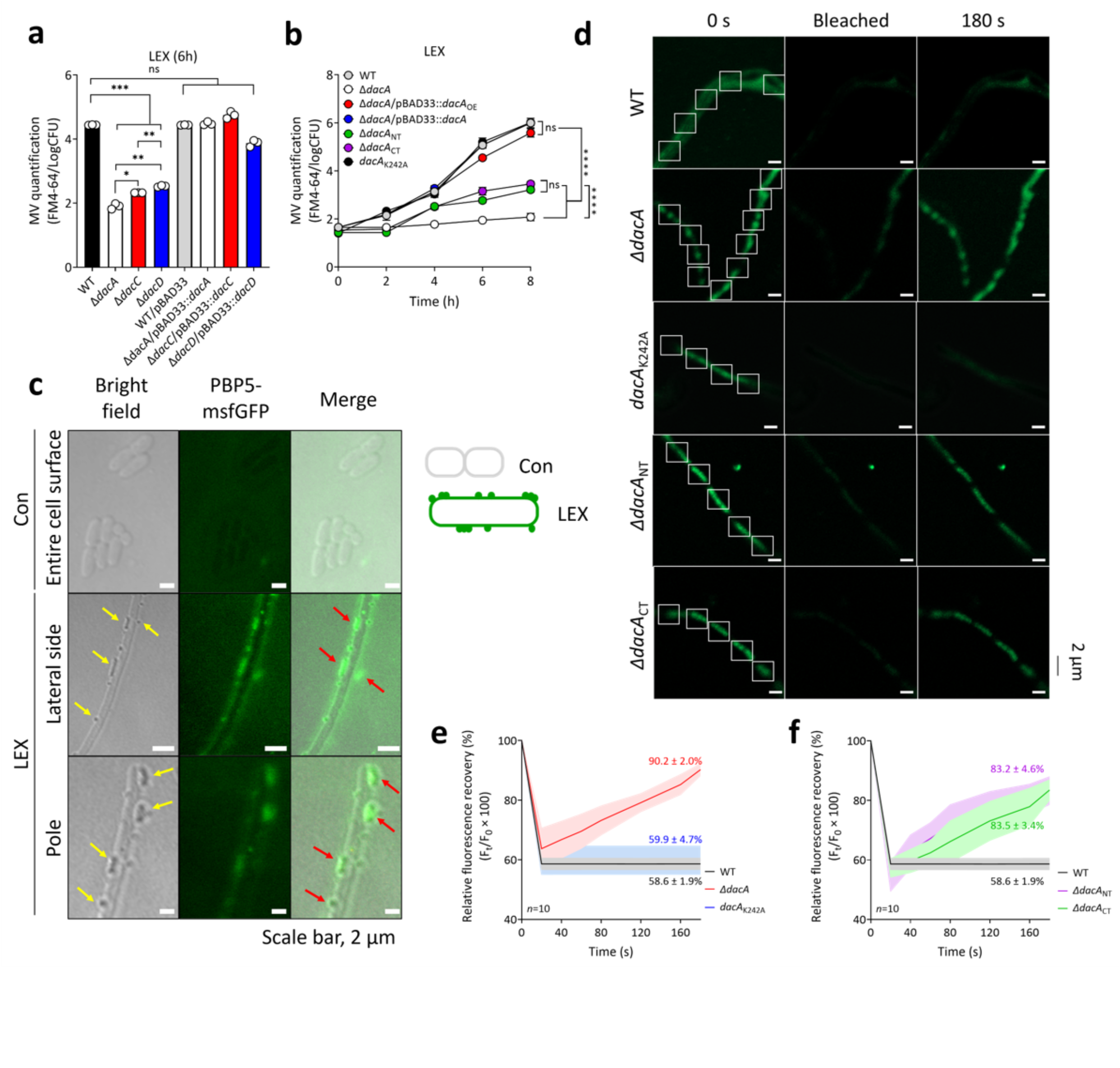
Measurement of MV level and periplasm crowding in WT and *dacA* variants. **a, b** MV production in WT, *AdacA*, *AdacC*, and *AdacD* single-knockout strains, their complemented strains, and *dacA* variant strains under LEX stress (N=3). **c** Localization of PBP5–msfGFP in WT at 6 h after LEX treatment, visualized by CLSM. Yellow arrows, cell-associated MVs; Red arrows, sites of colocalization between MVs and PBP5. **d** Fluorescence recovery after photobleaching (FRAP) analyses were performed in WT and *dacA* variant strains harboring pSEVA234::SP*_dsbA_*–msfGFP (N=10). Cells were grown at 37°C in the presence of LEX for 6 h, then shifted to 26°C and induced with 0.02 mM IPTG for 30 min prior to FRAP analyses. **e, f** FRAP recovery kinetics over a 180-s time course. Statistical comparisons were performed using one-way ANOVA with Tukey’s multiple comparison test (***P < 0.001 and ****P < 0.0001; ns, not significant).

We next constructed a chromosomally encoded PBP5 fused with monomeric superfolder GFP (msfGFP) and examined whether the sites of PBP5 accumulation coincide with MV formation at 6 h **(Fig. 3c)** (31,32). Interestingly, under LEX stress, PBP5–msfGFP clearly localized to the periplasm and formed discrete lateral accumulations **(yellow arrows, Fig. 3c)**. Notably, localized PBP5–msfGFP fluorescence signals were observed at sites associated with MV formation under LEX stress **(red arrows, Fig. 3c)**. Additionally, periplasmic crowding was assessed under LEX conditions in all tested strains using a fluorescence recovery after photobleaching (FRAP) assay with an IPTG-inducible periplasmic GFP construct (SP*_dsbA_*–sfGFP), which enabled monitoring of crowding-dependent diffusivity and fluorescence recovery rates **(Fig. 3d–f, S11a, S11b)** (33). Immediately after photobleaching, the fluorescence intensity of sfGFP decreased to ∼ 40% of the pre-bleach level in all tested strains **(Fig. 3d-f)**. In WT strain, the bleached region showed no fluorescence recovery during the 180-s period, indicating restricted periplasmic mobility under LEX stress **(Fig. 3d, 3e)**. In contrast, the *AdacA* mutant exhibited a gradual fluorescence recovery, reaching 90.2% of the initial intensity at 180 s, consistent with a markedly reduced periplasmic crowding state **(Fig. 3d, 3e)**. Interestingly, the truncation mutants displayed intermediate recovery (83.2 % and 83.5 %, respectively), whereas the catalytic-site mutant *dacA*_K242A_ showed no recovery **(Fig. 3d, 3f)**. To directly test whether the altered diffusion dynamics reflected changes in periplasmic protein content, the total periplasmic protein level was quantified **(Fig. S12)**. Under non-stress conditions, no significant differences were observed among strains, indicating comparable basal periplasmic composition **(Fig. S12a)**. In contrast, under LEX stress, WT cells exhibited a marked increase in periplasmic protein abundance, whereas the *AdacA* mutant showed a significant reduction (∼0.5 fold) compared to WT and *dacA*_K242A_ strains **(Fig. S12b)**. Notably, the truncation mutants displayed intermediate levels, while the *dacA*_K242A_ strain retained WT-like accumulation. This pattern closely mirrored the FRAP recovery profiles, supporting the conclusion that PBP5-dependent periplasmic protein accumulation under LEX stress underlies the observed crowding state. Together, our results indicate that periplasmic crowding mediated by the accumulation of PBP5 drives hypervesiculation under LEX stress.

### Non-canonical interaction between PBP5 and ACP under LEX stress

Although periplasmic chaperones and proteases maintain periplasmic quality control by removing mis-/unfolded proteins and debris, excessive accumulation of PBP5 under LEX stress leads to periplasmic crowding. We hypothesized that PBP5 interacts with unidentified periplasmic proteins, which may stabilize PBP5 and promote its accumulation in the periplasm. We aimed to identify PBP5-associated proteins under LEX stress at an early time point (4 h), when periplasmic crowding is established but before overt hypervesiculation, to uncover factors contributing to periplasmic crowding. Accordingly, PBP5-associated complexes were isolated by in vivo crosslinking at 4 h after LEX exposure in a chromosomally FLAG-tagged PBP5 strain **(Fig. 4a)** (29). Our immunoblot analyses revealed a prominent PBP5 band and several distinct bands in both PBP5-associated complexes **(Fig. 4a)**. The PBP5-containing complexes were further analyzed by mass spectrometry, revealing eight proteins annotated as cytosolic or extracellular, including PBP5 itself **(Table 1**, **Fig. 4b)**. Highly abundant cytosolic proteins such as GAPDH, GroEL, and EF-Tu were interpreted cautiously as potential nonspecific or transiently crosslinked proteins rather than definitive PBP5-interacting partners under both conditions (6,14,30). Surprisingly, one of the essential cytosolic proteins, ACP, was only detected exclusively in PBP5-containing complexes isolated under LEX stress. When massive MV occurs at 6 hours after LEX exposure, our membrane proteomic analyses showed that the high amounts of ACP were detected in OM fractions, suggesting that a large amount of ACP becomes mislocalized to membrane-associated regions, potentially as a consequence of stress-induced IM leakage **(Table S5-6, Fig. S13a)**.

**Figure 4.**
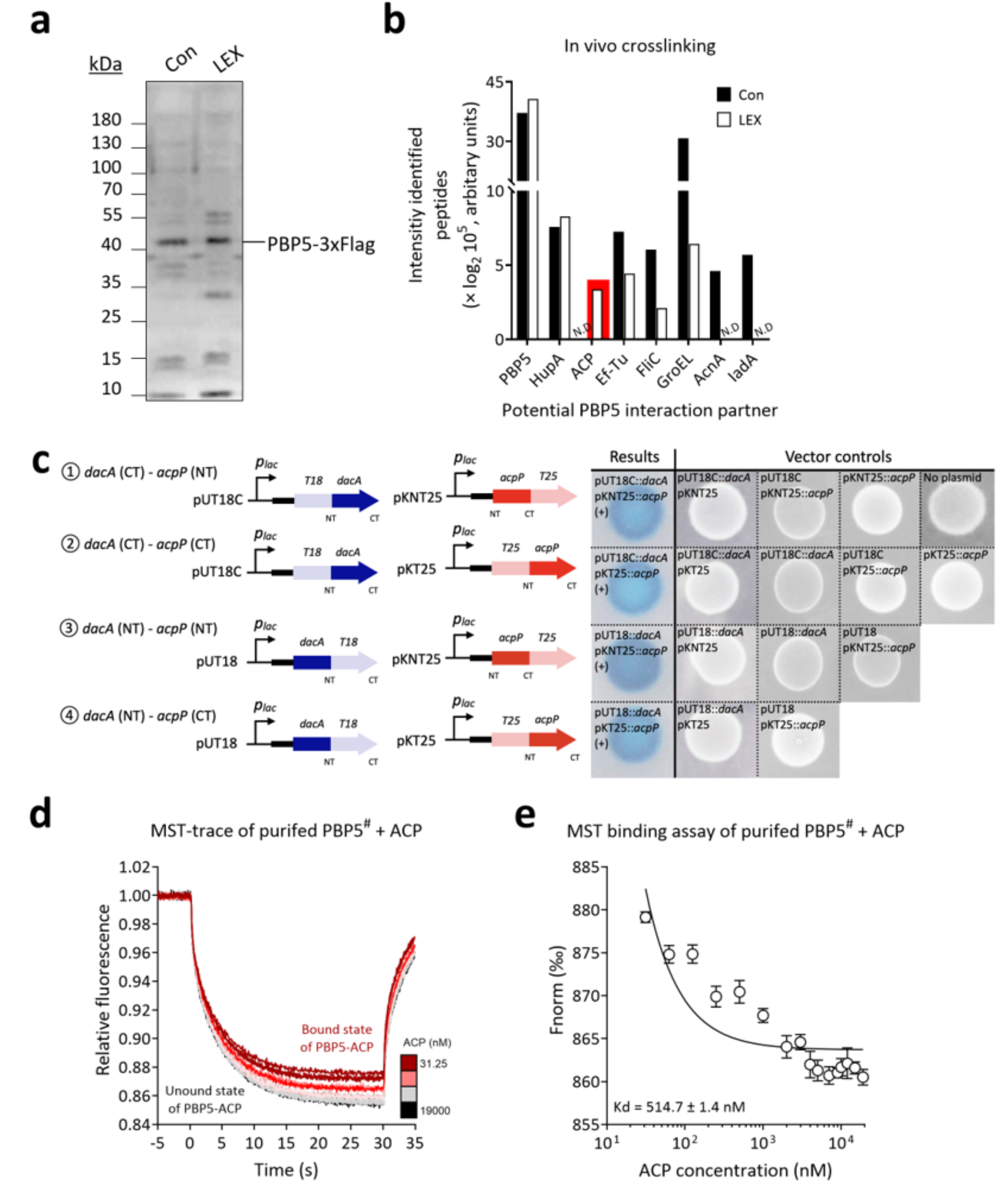
Non-canonical interaction between PBP5 and ACP under LEX stress. **a** In vivo crosslinking analyses of PBP5-associated complexes in a chromosomally FLAG-tagged PBP5 strain. Protein complexes were crosslinked using disuccinimidyl suberate (DSS) and subjected to anti-FLAG immunoprecipitation, followed by immunoblotting. Samples were prepared under control and LEX stress conditions. **b** Identification of PBP5-associated proteins under control and LEX conditions by LC/MS analysis of immunoprecipitated complexes. **c** In vivo validation of PBP5–ACP interaction using the bacterial adenylate cyclase two-hybrid (BACTH) system *in E. coli* BTH101. Co-transformants harboring pUT18C-*dacA*_ΔSP_ or pUT18-*dacA*_ΔSP_ in combination with pKT25-*acpP* or pKNT25-*acpP* were grown on LB agar supplemented with ampicillin (50 µg/mL), kanamycin (25 µg/mL), IPTG (40 µg/mL), and X-Gal (0.5 mM) and incubated at 37 °C for 24 h in the dark prior to imaging. **d, e** Microscale thermophoresis (MST) analyses of purified PBP5^#^ and ACP using a Monolith NT.115 instrument. ACP was subjected to thrombin cleavage prior to analysis. Fluorescently labeled PBP5 (400 nM) was used as the fixed binding partner, and increasing concentrations of ACP (31.25 -19,000 nM) were titrated to determine binding affinity. *SP, signal peptide.

**Table 1.** Proteins identified in PBP5-FLAG tagged strain by DSS-based in vivo crosslinking with anti-FLAG under control and LEX stress.

| **Gene** | **Protein** | **Fold change** | **Normalized data (Log<sub>2</sub>)** |  | **MW (kDa)** | **Coverage (%)** | **Accession number** | **Subcellular localization** |
|---|---|---|---|---|---|---|---|---|
|  |  | **LEX/Con** | **Con** | **LEX** |  |  |  |  |
| *dacA* | DD-carboxypeptidase PBP5 | 1.7 | 21.8 | 22.6 | 44.5 | 2 | A0A0F6AYD4 | Periplasm |
| *hupA* | DNA-binding protein HU-alph HupA | 1.7 | 19.5 | 20.3 | 9.5 | 14 | A0A0F6BA04 | Cytosol |
| *acpP* | Acyl carrier protein ACP | - | N.D | 19.0 | 8.6 | 12 | A0A0F6B032 | Cytosol |
| *tuf_2* | Elongation factor Tu | 0.9 | 19.5 | 19.4 | 43.3 | 3 | A0A0F6B9X6 | Cytosol |
| *fliC* | Flagellin FliC | 0.5 | 19.2 | 18.3 | 51.6 | 2 | A0A0F6B2U2 | Extracellular |
| *groEL* | Chaperonin GroEL | 0.3 | 21.6 | 19.9 | 57.3 | 7 | A0A0F6BAJ3 | Cytosol |
| *acnA* | Aconitate hydratase AcnA | - | 18.8 | N.D | 97.4 | 1 | A0A0F6B200 | Cytosol |
| *iadA* | Isoaspartyl dipeptidase IadA | - | 19.1 | N.D | 40.3 | 3 | A0A0F6BB47 | Cytosol |

\*DSS, Disuccinimidyl suberate; N.D, Not detected.

As PBP5 and ACP localize different subcellular compartments under basal conditions, this in vivo crosslinking result was further evaluated via bacterial two-hybrid (BACTH) and microscale thermophoresis (MST) assays **(Fig. 4c-e)** (31–34). In the BACTH assay performed in *E. coli* BTH101, T18-fused PBP5* lacking its signal peptide (hereafter referred to as PBP5*; *dacA* gene) was co-expressed with T25-fused ACP (*acpP*) or GAPDH (*gapA*) **(Fig. 4c, S14)**. GAPDH, an abundant cytosolic protein also reported in bacterial MVs, was included as a comparison partner to assess whether the observed interaction with PBP5* was specific to ACP rather than a general feature of abundant cytosolic proteins (6,14,30). Our BACTH analyses showed blue coloration for all combinations of N- and C-terminal fusions between PBP5* and ACP **(Fig. 4c)**. In contrast, no blue coloration was observed for any of the N- or C-terminal fusion combinations between PBP5* and GAPDH **(Fig. S14a)**. Consistent with these observations, β-galactosidase activity was detected only in the PBP5*– ACP combinations, whereas no detectable activity was observed in any of the PBP5*– GAPDH combinations **(Fig. S14b)**. Our MST analyses using purified PBP5^#^ lacking both the signal peptide and IM anchoring region (hereafter referred to as PBP5^#^) and ACP revealed an ACP concentration-dependent negative shift in thermophoretic behavior, with increasing ACP concentrations producing a distinct bound-state signal that was clearly separated from the unbound baseline **(Fig. 4d, S15)** (34). A binding curve generated from these fluorescence traces yielded a dissociation constant (K_d_) of 514.7 ± 1.4 nM **(Fig. 4d, 4e)**. To further explore the potential interaction interface between PBP5* and ACP, in silico molecular docking simulations were performed using Alphafold3 and HADDOCK (ver. 2.5) **(Table S7, Fig. S16)** (35,36). The predicted interface positioned ACP toward a positively charged surface region within the N-terminal domain of PBP5*, supporting a potential contribution of electrostatic interactions to complex formation **(Fig. S16a-c)**. The dominant docking cluster exhibited a high HADDOCK score (−100.4 ± 2.8), indicative of a stable and reproducible interaction mode **(Table S7)**. Inspection of the predicted docking complex further revealed that ACP is positioned within an active-site-proximal cavity of PBP5*, where it forms hydrogen-bond interactions with residues Thr_273_ and Lys_275_. The presence of positively charged residues in this region further suggests that electrostatic complementarity may contribute to stabilizing the complex **(Fig. S16c)**. Docking analyses of truncated PBP5* variants lacking the N-terminal (PBP5*_ΔNT_) or C-terminal domain (PBP5*_ΔCT_) resulted in less favorable HADDOCK scores (−66.3 ± 9.8 and −88.8 ± 4.7, respectively) than that of full-length PBP5*, while still indicating potential electrostatic contributions to ACP binding **(Table S7, Fig. S16d-e)**. Taken together, our complementary experimental and computational analyses strongly support a specific non-canonical interaction between PBP5 and ACP under LEX stress.

### Mislocalized ACP-mediated PBP5-ACP interaction in MV biogenesis

We next sought to understand how the cytosolic protein ACP becomes mislocalized under LEX stress. Given that ACP was detected in complex with PBP5 and in both IM and OM fractions under LEX stress, perturbation of IM integrity may underline its aberrant localization. To investigate this possibility, we observed the ultrastructure of the cell envelope using cryogenic transmission electron microscopy (cryo-TEM) in both WT and *AdacA* strains under control and LEX stress conditions **(Fig. 5a)**. Under control conditions, both strains exhibited intact and continuous inner and OM with well-defined bilayer structures **(Fig. 5a)**. Remarkably, LEX-exposed WT cells exhibited irregular and undulated IM structures. In contrast, LEX-exposed *AdacA* cells maintained largely intact IM structures **(Fig. 5a)**. Consistent with these observations, periplasmic width was dramatically increased in LEX-exposed WT cells, whereas LEX-exposed *AdacA* cells showed periplasmic widths comparable to those of control cells **(Fig. 5b)**. Cryo-electron tomography followed by three-dimensional rendering was used to visualize the overall cell-envelope architecture of WT cells under control and LEX conditions **(Fig. 5c, 5d, Videos S1–3)**. Interestingly, while the OM and IM remained continuous and well organized under control conditions **(Fig. 5c)**, irregular OM protrusions and apparent membrane detachment were observed under LEX stress, giving rise to vesicle-like structures **(purple; Fig. 5d)**. MV formation was observed not only at septal regions but also along the lateral cell envelope. Consistent with our previous native-TEM observation, we also identified two distinct vesicle types **(Fig. S17a, S17b)**. Concurrently, the IM displayed irregular and locally disrupted structures, including regions that appeared as hole-like discontinuities **(blue arrowhead; Fig. 5d)**. At the division site, the OM and IM were not clearly separated and appeared to remain connected. These observations are consistent with PBP5-dependent local disruption of IM integrity under LEX stress.

**Figure 5.**
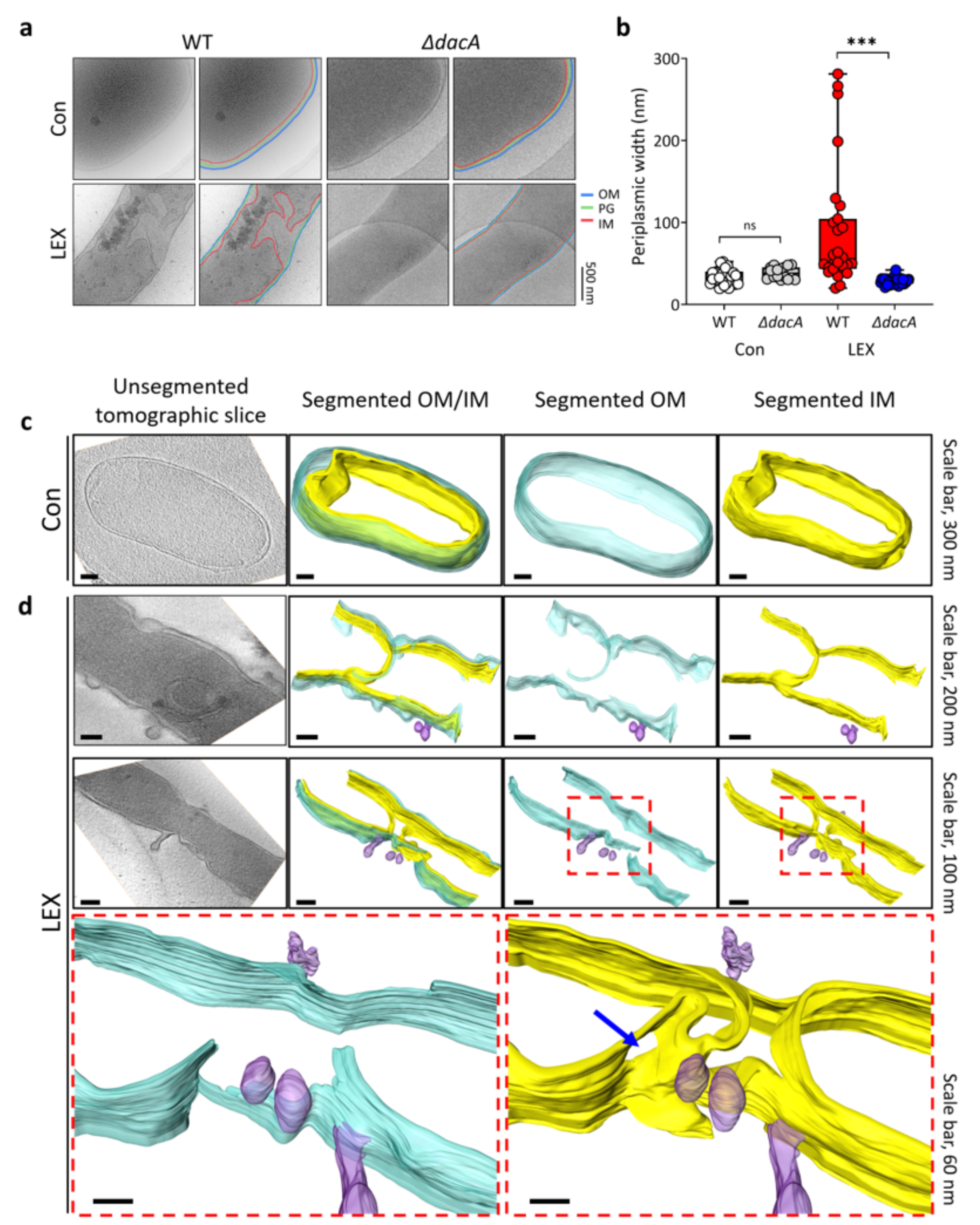
PBP5-dependent ACP mislocalization under LEX stress. **a** Cryogenic-TEM analyses of WT and *AdacA* cells under control and LEX stress conditions. OM, PG, and IM are shown in blue, green, and red, respectively. **b** Quantification of periplasmic width from the cryo-TEM images (n=28, 16, 27, and 29 cells for WT control, *AdacA* control, WT LEX, and *AdacA* LEX, respectively). **c, d** Cryo-tomograms and 3D segmentations of WT cells under control and LEX stress conditions. The OM and IM are shown in cyan and yellow, respectively. Blue arrowheads indicate IM holes, and red dashed boxes indicate magnified regions. Statistical comparisons were performed using one-way ANOVA with Tukey’s multiple comparison test ***P < 0.001; ns, not significant).

Based on the observed disruption of IM integrity, ACP localization in the membrane fraction was examined using a WT strain carrying chromosomally 3×FLAG-tagged ACP under LEX stress **(Fig. S18a–c)**. Our Western blot analyses showed that ACP levels in the membrane fraction were higher under LEX stress than under control conditions, particularly at 6 h (∼2-fold), whereas ACP levels in the cytosolic fraction progressively decreased over time **(Fig. S18a–c)**. To further validate ACP mislocalization and its periplasmic translocation in vivo, a HiBiT–LgBiT system was established by fusing ACP at its C-terminus to a HiBiT tag and pairing it with periplasmically localized LgBiT **(Fig. S19a–e)** (37). Luminescence is generated only when ACP-HiBiT enters periplasm and complements periplasmic LgBiT to restore luciferase activity **(Fig. S19e)**. Because MV production and PBP5 accumulation are more pronounced at higher LEX concentrations, we next examined whether ACP leakage into the periplasm shows a similar pattern across the same LEX concentration range (1/32– 1/4×MIC) **(Fig. S19f)**. The luminescence signal became progressively stronger with higher LEX concentrations, indicating enhanced periplasmic detection of ACP-HiBiT under LEX stress. The luminescence signal became progressively stronger with higher LEX concentrations **(Fig. S19f)**. At 6 h, cells treated with 1/4×MIC LEX showed markedly higher luminescence than the untreated control (∼80-fold). Periplasmic ACP levels were then compared among WT and *dacA* variant strains using the ACP-HiBiT/periplasmic LgBiT system at 6 h under LEX stress **(Fig. S19g)**. The WT and *dacA*_K242A_ strains showed substantially higher luminescence than the *dacA* deletion/truncation variants (∼8-fold). Taken together, our cryo-tomographic analyses and HiBiT–LgBiT assay results support a role for PBP5 in promoting the translocation of cytosolic ACP into the periplasm under LEX stress.

### Periplasmically mislocalized ACP–mediated accumulation of PBP5

To assess the contribution of periplasmically mislocalized ACP to MV formation via its non-canonical interaction with PBP5, changes in MV levels under LEX stress were evaluated in a CRISPRi-based *acpP* knockdown strain **(Fig. 6a, 6b, S20a, S20b)**. To exclude growth-related effects, *acpP* knockdown was finely tuned by adjusting the aTc concentration (50 nM) to a level that did not alter bacterial growth **(Fig. 6a)**. Consistently, no detectable differences in cellular morphology were observed between the *acpP* knockdown strain and the control strain under either control or LEX stress conditions **(Fig. S20c)**. Surprisingly, *acpP* knockdown under LEX stress resulted in a ∼0.5-fold decrease in MV production **(Fig. 6b)**. CRISPRi-mediated *acpP* knockdown did not result in detectable changes in PBP5 band intensity by Western blotting over the course of cultivation **(Fig. S21a-c)**. To further investigate the contribution of the PBP5–ACP interaction, we generated a PBP5 interface mutant (*dacA*_T273A/K275A_), which were predicted to form hydrogen bonds with ACP in our in silico docking analysis **(Fig. S16, S22)**. These substitutions were designed to weaken the predicted PBP5*–ACP interaction. Although the PBP5 interface mutant retained a positive interaction signal in the BACTH assay, its β-galactosidase activity was lower than that of WT PBP5* paired with ACP **(Fig. S22a, S22b)**. To assess the biological significance of the PBP5–ACP interaction in MV formation, we next examined bacterial growth and MV production using the PBP5 interface mutant **(Fig. 6c, 6d, S22c, S22d)**. The *dacA*_T273A/K275A_ interface mutant exhibited growth profiles comparable to those of the WT *dacA*-complemented strain under both control and LEX stress conditions **(Fig. 6c, S22c)**. However, MV production was significantly reduced in the *dacA*_T273A/K275A_ interface mutant compared with the WT *dacA*-complemented strain under LEX stress **(Fig. 6d, S22d**). Together, the reduced MV production following *acpP* knockdown and in the PBP5–ACP interface mutant suggests that the PBP5–ACP interaction strongly influences the level of MV production under LEX stress.

**Figure 6.**
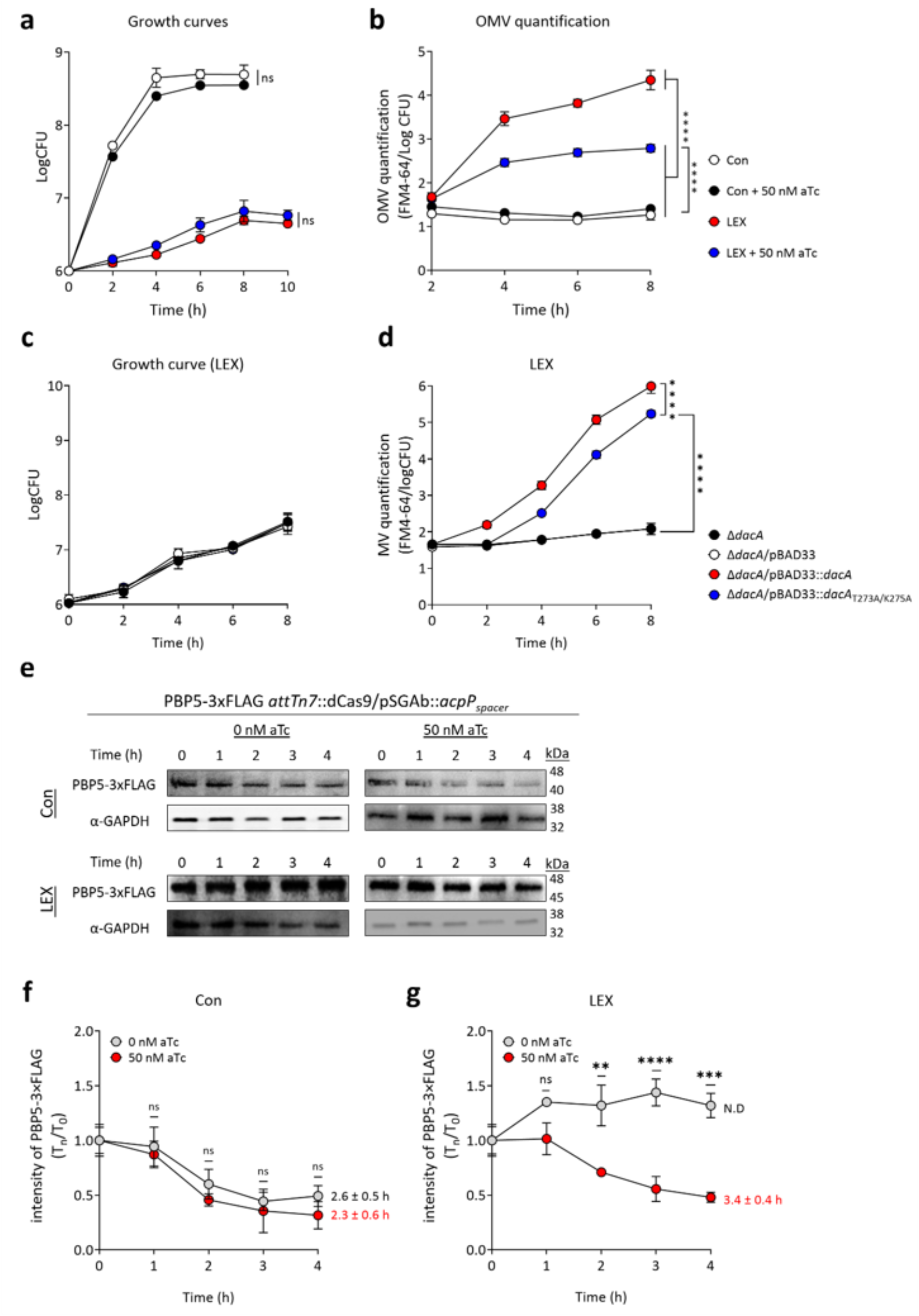
Quantification of MV levels and PBP5 stability in *acpP* knockdown strain. **a, b** Measurement of growth curves and MV level in the *acpP*^down^ strain (*attTn7*::dCas9/pSGAb::*acpP*_spacer_) under control and LEX stress conditions at 37°C. Data represent three independent biological replicates (N=3). **c, d** Growth curves and MV quantification of the Δ*dacA* mutant and its derivatives carrying empty pBAD33 or pBAD33 expressing wild-type PBP5 (*dacA*) or the PBP5 interface mutant (T273A/K275A) under the control of the native *dacA* promoter. **e–g** Chloramphenicol-chase–based assessment of PBP5 stability in *acpP* knockdown strain carrying chromosomally 3×FLAG-tagged PBP5. Cells were grown at 37 °C for 6 h under LEX stress, followed by treatment with chloramphenicol (200 µg/mL). PBP5 levels were monitored over time after translational arrest. Data represent two independent biological experiments (N=3). GAPDH was used as a loading control. Representative Western blotting images are shown. Statistical comparisons were performed using one-way ANOVA with Tukey’s multiple comparison test (**P < 0.01, ***P < 0.001, and ****P < 0.0001; ns, not significant).

Given that MV production was reduced in the *acpP* knockdown and *dacA*_T273A/K275A_ interface mutant strains, we reasoned that the PBP5–ACP interaction may prolong the periplasmic residence of PBP5, leading to its accumulation and contributing to periplasmic crowding under LEX stress. To determine whether the PBP5–ACP interaction influences the periplasmic persistence of PBP5, its stability was assessed by in vivo chloramphenicol-chase assays and in vitro half-life measurements in the presence or absence of ACP depletion **(Fig. 6e–g)** (38,39). Our chloramphenicol-chase analyses revealed no significant difference in the half-life of PBP5 between the *acpP* knockdown strain and the corresponding dCas9-uninduced control (2.3 ± 0.6 h vs. 2.6 ± 0.5 h) under control condition **(Fig. 6f)**. *acpP* depletion resulted in a prolonged half-life of PBP5 (3.4 ± 0.4 h) under LEX stress, compared with the half-life of PBP5 in the tested strains under control conditions, indicating reduced turnover of PBP5 under LEX stress **(Fig. 6g)**. Interestingly, under LEX stress, PBP5 signal intensity did not decline over the chase period in the dCas9-uninduced control and instead showed a relative increase **(Fig. 6g)**. This pattern is more likely attributable to the persistence of pre-existing PBP5 than to de novo protein synthesis. Collectively, our MV production and PBP5 half-life analyses using the *acpP* knockdown strain support a model in which cytosolic ACP, upon mislocalization to the periplasm under LEX stress, interacts with and stabilizes PBP5. Increased PBP5 stability may facilitate its accumulation in the periplasm, thereby contributing to periplasmic crowding and subsequent MV formation.

### PBP5-dependent periplasmic crowding–induced OM–PG tethering defects

Although ACP-mediated PBP5 accumulation established a crowded periplasmic state under LEX stress, the physical step linking this state to OM blebbing remained unclear. Because PBP5-dependent periplasmic crowding was associated with increased MV production, this crowded state may interfere with OM–PG tethering mediated by the major linkage factors OmpA and Lpp (40,41). We assessed whether OM–PG tethering is weakened under LEX stress by analyzing OmpA and Lpp at 6 h using Western blotting **(Fig. 7)**. In WT strain, OmpA abundance decreased in both crude extracts and OM-enriched fractions under LEX stress relative to the control condition (∼0.5-fold), whereas Lpp modestly increased in these fractions (∼1.3–1.4-fold) **(Fig. 7a, 7b)**. In contrast, the PG-associated pools of both OmpA and Lpp were reduced under LEX stress (∼0.5-fold) **(Fig. 7c)**. Time-course analyses of free Lpp using NHS-LC-LC-biotin labeling revealed progressive accumulation of detached Lpp under LEX stress **(Fig. S23a, S23b)** (42). As expected, OmpA- and Lpp-mediated OM–PG linkage is reduced under LEX stress, despite increased OM-localized Lpp. However, *acpP* knockdown increased OmpA abundance, whereas Lpp abundance was reduced in these fractions relative to the dCas9-uninduced control **(Fig. 7d, 7e)**. In contrast, the PG-associated pools of both OmpA and Lpp were restored upon *acpP* knockdown **(Fig. 7f)**. These results suggest that ACP depletion reduces PBP5 accumulation and periplasmic crowding, thereby partially restoring OmpA- and Lpp-mediated OM–PG linkage under LEX stress. We next compared the levels of PG-tethered OmpA and Lpp among WT and four *dacA* variants under LEX stress **(Fig. 7g, S24a, S24b)**. Under LEX stress, no significant differences in total OmpA or Lpp abundance were observed among all tested strains in crude extracts **(Fig. S24a)**. In the OM-enriched fraction, however, low-vesiculating strains exhibited higher OmpA abundance (∼2.2-fold) and lower Lpp abundance (∼0.8-fold) than hypervesiculating strains (WT and *dacA*_K242A_) **(Fig. S24b)**. In the PG-linked fraction, both OmpA and Lpp were relatively enriched in the truncation/deletion mutants compared with hypervesiculating strains (OmpA, ∼2.1-fold; Lpp, ∼1.3-fold) **(Fig. 7g)**. Consistently, free Lpp accumulated more rapidly in hypervesiculating strains than in low-vesiculating strains **(Fig. S23c–f)**. Together, these findings suggest that PBP5-dependent periplasmic crowding is associated with reduced OmpA- and Lpp-mediated OM-PG linkage, facilitating MV formation under LEX stress.

**Figure 7.**
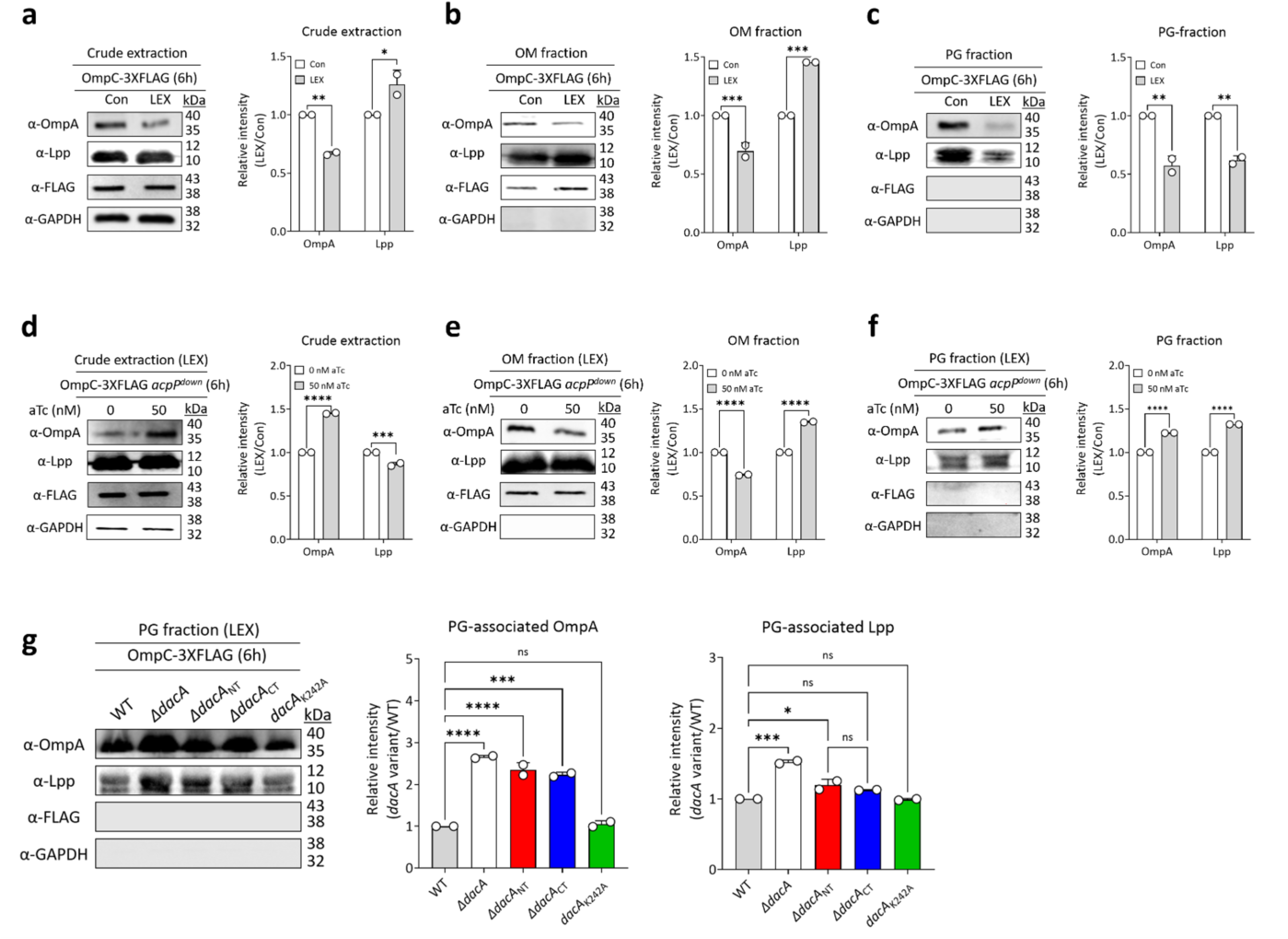
PBP5-dependent periplasmic crowding–induced OM–PG tethering defects. **a–c** Western blot analyses of OmpA and Lpp levels in crude extracts, OM fractions, and PG-associated fractions of the WT strain carrying chromosomally tagged OmpC-3×FLAG under control and LEX stress conditions for 6 h. Band intensities of OmpA and Lpp were quantified and normalized to the untreated control. **d–f** Western blot analyses of OmpA and Lpp levels in crude extracts, OM fractions, and PG-associated fractions of the OmpC-3×FLAG *acpP* knockdown strain under LEX stress, with or without aTc-mediated *acpP* repression. Band intensities were quantified and normalized to the non-induced condition. **g** Western blot analyses of OmpA and Lpp levels in PG-associated fractions of WT and *dacA* variant strains grown under LEX stress for 6 h. Total OmpA and Lpp levels in PG-associated pools were comparable among strains. OmpC-3×FLAG was used as an OM-associated marker, and GAPDH was used as a cytosolic control. Individual points indicate biological replicates, and bars indicate mean values. Statistical comparisons were performed using one-way ANOVA with Tukey’s multiple comparison test (*P < 0.05, **P < 0.01, ***P < 0.001, and ****P < 0.0001; ns, not significant).

## Discussion

In *Salmonella*, PhoPQ-regulated PagL-mediated lipid A deacylation alters lipid A topology and acid-induced PagC conformational reshaping increases lateral pressure in outer leaflet of OM, thereby promoting local OM curvature and vesiculation (43,44). However, PagL was not identified and PagC abundance was reduced (∼0.6-fold), indicating that LEX-induced hypervesiculation does not simply follow the previously described PhoPQ-mediated OMV biogenesis pathway **(Table S6, Fig. S13b)**. Instead, the upregulation of the periplasmic chaperone/proteases DegP and DegQ (∼1.8-fold) suggests an increased envelope proteostasis burden under LEX stress, potentially caused by the accumulation of misfolded envelope proteins **(Table S6, Fig. S13b)**. This pattern is reminiscent of hypervesiculation phenotypes reported in periplasmic quality-control malfunctional mutants (e.g., *AdegP* and *AdegQ*) (6,45). Crowding-driven OMV formation requires coupling with concurrent lytic transglycosylase, which cleave the PG network and thereby enable OMV formation in *Acinetobacter baumannii* (6). In our study, LEX-induced PBP3 inactivation and the concomitant accumulation PBP5 reveals a previously unrecognized route to periplasmic crowding, in which accumulated PBP5 forms a non-canonical complex with mislocalized ACP **(Fig. 4)**. Consistent with this coupled model, our proteomic analyses showed increased abundance (∼1.8-fold) of PG-hydrolases, including lytic transglycosylases RlpA and MltG, under LEX stress **(Table S6, Fig. S13c)**. Our data further reveal that PBP5-dependent periplasmic crowding impairs OM–PG coupling under LEX stress **(Fig. 7, S23, S24)**. Unlike classical models in which hypervesiculation is driven by the loss or dysfunction of canonical OM–PG linkage factors such as OmpA, Lpp, or Tol–Pal components, our MV model suggests a previously unrecognized mechanism in which periplasmic accumulation of the PBP5–ACP complex weakens OmpA–PG and Lpp–PG coupling (1,3). Although we propose that PBP5-dependent periplasmic crowding physically interferes with OM–PG coupling, the possibility remains that this weakening results from altered activity of PG-linking enzymes, such as L,D-transpeptidases, rather than direct physical obstruction.

An unresolved question is how *dacA* expression is induced during LEX stress. BolA represents a plausible candidate regulator for PBP5 induction, as BolA has been reported to contribute to cell-shape homeostasis through regulation of *dacA* expression in *E. coli* (46). However, whether LEX-induced PBP5 upregulation in *Salmonella* is directly mediated by BolA remains unclear. Fine-scale mapping of the *dacA* promoter uncovered three putative transcription factor-binding motifs (BolA, LexA, and NagC) **(Fig. S25a)**. Consistent with a possible link to stress-responsive regulation, *dacA* expression was upregulated (∼1.7-fold) under mitomycin C stress, suggesting that DNA damage/SOS-associated stress may contribute to *dacA* induction **(Fig. S25b, S25c)** (47). Indeed, our proteomic analyses revealed increased abundance of the cell envelope stress regulators (e.g., CpxAR, PhoPQ, and RpoE) under LEX stress **(Fig. S25d)**. Notably, only the *AcpxR* strain exhibited elevated *dacA e*xpression, suggesting that CpxR may indirectly or negatively influence *dacA* expression under LEX stress **(Fig. S25d)**. Although the regulatory basis of LEX-induced PBP5 upregulation remains unresolved, increased PBP5 levels may help maintain permissive PG insertion zones under envelope stress (48). By removing the terminal D-Ala from PG pentapeptide stems, PBP5 generates tetrapeptide stems with reduced crosslinking potential, thereby locally loosening the PG network and maintaining sufficient spatial flexibility for continued PG synthesis and remodeling (48). Given that PBP5 has been linked to both elongasome- and divisome-associated PG synthesis machineries, elevated PBP5 levels may function as an optimizer of local PG architecture, preserving insertion zones that remain accessible to multiple PG synthases and hydrolases (29,49). Under LEX stress, where PBP3 activity is impaired and envelope remodeling demand is increased, this response may help sustain localized PG synthesis and remodeling while preventing excessive structural rigidity.

The incorporation of cytoplasmic proteins into OMVs has long remained one of the most enigmatic aspects of OMV biogenesis. Current models predict that outward bulging of the OM preferentially captures adjacent periplasmic proteins, whereas cytoplasmic proteins lacking signal sequences are expected to be excluded from vesiculation sites by the IM barrier (1–3,11,30). Several scenarios may be envisioned to explain how cytoplasmic proteins appear in OMVs even under normal growth conditions. For example, cytoplasmic proteins may be passively co-captured through transient hydrophobic interactions with envelope proteins during their nascent translocation and folding. In addition, transient envelope remodeling during cell division may create short-lived local defects or compartmental relaxation, allowing cytoplasmic proteins to access periplasm-proximal regions. Alternatively, stress-induced IM remodeling may provide another route for cytoplasmic protein incorporation, as ciprofloxacin-induced activation of the phage shock protein (Psp) response has been shown to promote the production of MVs containing cytosolic Cre recombinase in *E. coli*, potentially through PspA-associated membrane remodeling, given its homology to eukaryotic ESCRT-III proteins involved in membrane curvature and fission (30,50).

Here, our findings suggest that excessive cell elongation may impose mechanical strain on the envelope and transiently perturb IM integrity, providing the basis for the stepwise model of LEX-induced hypervesiculation proposed in **Fig. 8** (51). Under normal growth conditions, the OM remains physically coupled to the PG layer through OmpA and Lpp **(Fig. 8a)**. Upon exposure to sublethal LEX, inhibition of PBP3 impairs cell division, leading to filamentous elongation and concomitant upregulation of PBP5 **(Fig. 8b)**. LEX-mediated accumulation of PBP5 in the periplasm contributes to a crowded envelope state and is accompanied by patchy or irregular IM structures **(Fig. 8c)**. These local IM alterations may allow cytosolic ACP to access the periplasmic or membrane-proximal space, where it can encounter PBP5 and form a PBP5–ACP complex. The accumulation of the complex further increases periplasmic crowding and weakens OmpA- and Lpp-mediated OM–PG connectivity, allowing local OM bulging and ultimately leading to MV release **(Fig. 8d)**. Our native-TEM analyses revealed a small proportion of O-IMVs (0.6%), which may have arisen from elevated autolysin activity or lysis of a minor cell subpopulation **(Fig. 1h)** (1,11). Although Western blot analyses detected all four subcellular markers in the vesicle-enriched fractions, these bulk measurements could not determine whether the markers were associated with OMVs or O-IMVs **(Fig. S5b, S5c)**. Further analyses at the single-vesicle level will therefore be necessary to clarify the origin and molecular composition of these distinct vesicle types. The mislocalization of ACP raises an additional question regarding its biochemical state in the periplasm. Long-chain acylation (e.g., C_16:0_) may provide a hydrophobic contact point that allows mislocalized ACP to associate with IM-proximal regions and encounter PBP5 when IM leakage occurs (52,53). This possibility is consistent with the observation in *E. coli* showing that long-chain acyl-ACP, but not apo/holo-ACP and short-chain acyl-ACP (e.g., C_8:0_), was predominant in the IM fraction (52–54). Regardless of its acylation state, however, the key finding of this study is that transient disruption of normal compartmental boundaries creates an opportunity for ACP and PBP5 to encounter in the periplasm. Given the favorable HADDOCK docking scores, this observation suggests that the PBP5–ACP interaction is restricted primarily by subcellular compartmentalization rather than by structural incompatibility **(Fig. S16)**. Thus, under specific stress conditions that disrupt normal compartmental boundaries, proteins that would otherwise remain spatially separated may form non-canonical pairs and acquire context-dependent functions, as illustrated here by PBP5–ACP-mediated MV biogenesis.

**Figure 8.**
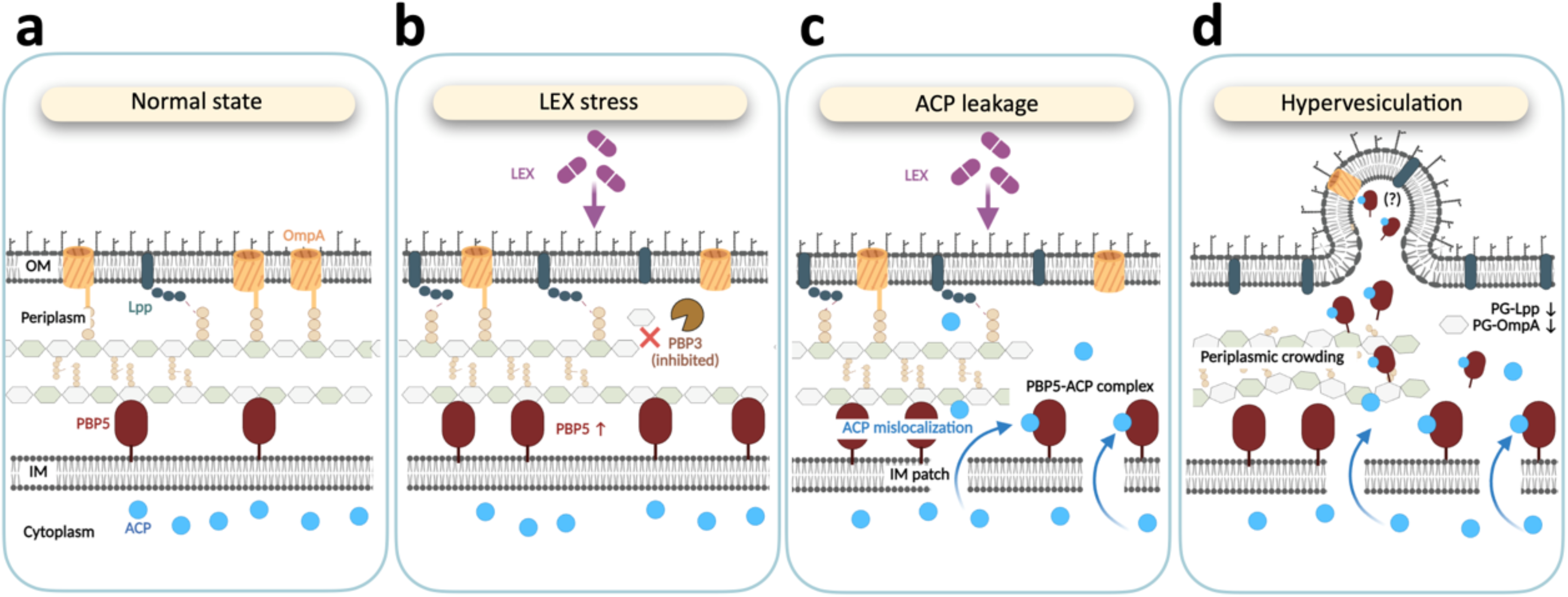
Proposed model for LEX-induced hypervesiculation. **a** Under normal growth conditions, OM is physically coupled to the PG layer through OM–PG tethering factors such as OmpA and Lpp. ACP is confined to the cytoplasm by an intact IM. **b** Exposure to sublethal concentrations of LEX inhibits PBP3, resulting in impaired cell division and filamentation. PBP5 is strongly upregulated under LEX stress. **c** LEX-mediated PBP5 accumulation in the periplasm is associated with local perturbations in IM integrity, allowing cytoplasmic ACP to access the periplasm. Mislocalized ACP forms a PBP5–ACP complex. **d** PBP5–ACP complex stabilizes PBP5 and promotes its accumulation in the periplasm. Accumulation of PBP5 interferes with normal OM–PG tethering mediated by OmpA and Lpp, weakening envelope connectivity and promoting hypervesiculation.

## Materials and methods

### Bacterial strains and culture conditions

The *S.* Typhimurium and *E. coli* used in this study, all isogenic to the WT strain 14028s, are shown in **Table S8**. All *S.* Typhimurium strains were cultured at 37°C in LB broth with static conditions. All strains were cultured overnight for 16 h. Following this, each O/N cell culture was diluted to a ratio of 1:100 in LB broth (5 mL) and then incubated at 37°C during the early phase (optical density, OD_600_, ∼0.3). All experiments were conducted with cell counts adjusted to a final concentration of 1×10^6^ CFU/mL in LB broth. To grow strains (*E. coli* and *S*. Typhimurium) bearing genetic elements conferring antibiotic resistance, the media were supplemented with chloramphenicol (25 or 50 µg/mL); kanamycin (50 or 100 µg/mL); ampicillin (100 or 500 µg/mL); gentamicin (25 or 50 µg/mL).

### Construction of *S.* Typhimurium isogenic mutants using λ-red recombination

All mutant strains were constructed using the one-step λ-red recombination (44). To delete the target genes, the Km^R^ and Cm^R^ cassettes were amplified from pKD3 and pKD4, respectively, using the designed primers **(Table S8)**. The antibiotic cassette was subsequently removed with pCP20 expressing flippase recombinases (55). For the construction of 3 x FLAG-tagged variants, 3 x FLAG sequences flanking target gene-specific sequences (40 bp) were amplified using pSUB11.

### Electron microscopy analysis

Electron microscopy was performed with modifications from previously reported procedures (56,57). WT cells were grown overnight in LB broth at 37°C and then diluted 1:100 into 5 mL of fresh LB medium. Cultures were incubated for 6 h in the presence or absence of LEX, and bacterial cells were collected by centrifugation at 13,000 rpm for 10 min. The resulting pellets were rinsed twice with PBS (pH 7.2) and fixed with 2.5% glutaraldehyde in 0.1 M phosphate buffer (pH 7.2) for 4 h at 4°C. After fixation, samples were briefly centrifuged at 13,000 rpm for 1 min, washed three times with PBS, and post-fixed with 1% osmium tetroxide for 2 h at 4°C to improve membrane contrast. The post-fixed cells were washed again with PBS and dehydrated in 100% ethanol. For FE-SEM analysis, dehydrated samples were resuspended in 30 µL of ethanol, and 5 µL of each suspension was placed onto aluminum stubs. After air drying, the samples were coated with platinum and examined using a field-emission scanning electron microscope (FE-SEM; Quanta 250, FEI, USA).

For thin-section TEM, fixed bacterial pellets were processed for resin embedding, ultrathin sectioning, and mounting on carbon-coated copper grids. Sections were stained with 2% uranyl acetate before observation using a transmission electron microscope (BIO TEM JEM-1011, JEOL, Japan). For negative-stain TEM, fixed cell suspensions were applied to carbon-coated copper grids, gently washed with distilled water, stained with 2% uranyl acetate, and imaged using the same TEM system (58).

### CRISPR interference

The aTc-inducible CRISPRi system was established using the mini-Tn7-based pYDE009 platform with minor modifications from a previously described method (24). Briefly, the mini-Tn7 cassette carrying the dCas9 module was integrated into *S.* Typhimurium 14028s containing pBAD33 by four-parental mating. The recipient *S.* Typhimurium strain, *E. coli* DH5a donor strains carrying pYDE009 or pTNS3, and the *E. coli* HB101 helper strain carrying pRK2013 were cultured overnight at 37°C with shaking at 180 rpm. Equal volumes of the four overnight cultures were mixed, collected by centrifugation at 7,600 × g for 1 min at 4°C, and sequentially washed with 10 mM MgSO_4_ and PBS. The washed cell mixture was resuspended, and 500 µL was applied onto a 0.45-µm-pore-size membrane filter placed on an LB agar plate. After incubation at 37°C for 4 h to allow conjugation, cells were recovered from the filter with 1 mL of 10 mM MgSO_4_. Aliquots of the recovered suspension were plated on LB agar containing chloramphenicol and gentamicin to select transconjugants. Correct insertion of the dCas9 cassette at the *attTn7* site was confirmed by PCR. The pBAD33 plasmid was subsequently removed by serial passage in antibiotic-free medium, and plasmid loss was verified by comparing growth on LB agar plates with or without chloramphenicol. For target-specific CRISPRi, spacer sequences against *ftsI* or *acpP* were inserted into the pSGAb sgRNA vector using Golden Gate Assembly. The resulting sgRNA plasmids were introduced into *S.* Typhimurium 14028s strain carrying chromosomally integrated dCas9, and gene repression was induced by adding anhydrotetracycline at the indicated concentrations.

### Assessment of antibiotic susceptibility in *S.* Typhimurium

Nine β-lactam antibiotics and mitomycin C were used individually in LB liquid to evaluate MIC tests based on the EUCAST guideline and database (https://mic.eucast.org/) **(Table S1)**. All cultured cells were transferred into 96 well plates containing each β-lactam antibiotic, and the final cell counts were adjusted to 1 × 10^6^ CFU/mL. The cells were incubated at 37°C for 24 h to measure the MICs. All experiments were performed in biological five replicates.

### Short-term time–kill curve assay of *S.* Typhimurium under β-lactam antibiotic stress

The time-kill assay of the WT strain was performed in LB added with β-lactam antibiotics (ranging from 1/8 to 1×MIC). Each β-lactam antibiotic-treated cells were harvested at each time (0, 20, 40, and 60 min) and washed twice with PBS, following which aliquots (10 µL) were spotted to LB agar plates for CFU assay. All experiments were performed in biological replicates. To confirm the emergence of mutations in cells exposed to LEX (1/4×MIC), the WT strain cells were exposed to LEX (1/4×MIC, for 6 hours), and then spread on an LB agar plate. Individual single colonies (*n*=10) were isolated, and the growth curves of the isolated individual single cell were measured in LB or LB supplemented with LEX (1/4×MIC).

### Flow cytometric measurement of PI-stained dead cells exposed to β-lactam antibiotics

The WT strain cells were cultured O/N in LB broth added with sub-inhibitory concentration of each β-lactam antibiotic at static conditions for the measurement of the PI-based dead cells. The cells (1×10^6^ CFU/mL) were inoculated into test tubes with a subinhibitory concentration (1/4×MIC) of each β-lactam antibiotic. After 6 h, the cells exposed to each β-lactam antibiotic were washed three times and then stained with PI (Invitrogen, USA; 1 µg/mL) for 30 min at room temperature in the dark. PI staining followed by flow cytometry was used to estimate the proportion of membrane-damaged or dead cells after β-lactam exposure, with modifications from previously described procedures (24,58). Overnight cultures of the WT strain were diluted into LB broth to a final density of 1 × 10^6^ CFU/mL and incubated under static conditions with each β-lactam antibiotic at 1/4×MIC. After 6 h of treatment, bacterial cells were collected, washed three times with PBS, and stained with propidium iodide (PI; Invitrogen, USA) at a final concentration of 1 µg/mL for 30 min at room temperature in the dark. Stained samples were analyzed using an Accuri C6 Plus flow cytometer (BD Biosciences, USA). PI fluorescence was detected using excitation and emission wavelengths of 535 and 615 nm, respectively, and the forward-scatter threshold was set to 10,000. For each condition, three independent biological replicates were analyzed, with approximately 50,000 events collected per replicate. The percentage of PI-positive cells was used to determine the fraction of dead or membrane-permeabilized cells following β-lactam antibiotic treatment.

### Conditioned medium re-inoculation assay

To verify non-lytic MV formation, conditioned medium re-inoculation assay was performed. For the conditioned medium re-inoculation assay, WT cells were exposed to LEX (1/4×MIC) for 6 h. The culture was centrifuged to separate cells from the conditioned medium, and the supernatant was filtered through a 0.22-µm filter to obtain cell-free conditioned medium containing residual LEX and preformed MVs. The harvested cells were washed with PBS and re-inoculated into the cell-free conditioned medium. Cell growth and MV production were monitored every 20 min for up to 2 h.

### Quantification and purification of MVs

Overnight cultures of all strains were diluted 1:100 in fresh LB broth and incubated at 37 °C for 3 h. The precultures were subsequently inoculated into fresh LB broth at a final concentration of 1×10^6^ CFU/mL and incubated for 6 h under the indicated experimental conditions. Following incubation, 1 mL of each bacterial culture was collected and centrifuged to remove bacterial cells. The resulting supernatant was filtered using a Millex-GP syringe filter (0.22 µm, Merck Millipore, USA). A 200 µL aliquot of the filtrate was transferred to each well of A 96-well plate (SPL Life Sciences, South Korea). FM4-64 dye was added to each well at a final concentration of 1 µg/mL. Fluorescence intensity was measured at an excitation wavelength of 510 nm and an emission wavelength of 610 nm using a fluorescence spectrophotometer (TECAN, Männedorf, Switzerland), with a slit width of 5 nm for both excitation and emission (6,14,59).

WT strain cells were grown in LB and LB with LEX (2 L), respectively, for 6 h. Cultures were centrifuged at 10,000×g (for 30 min at 4 °C), and the cell-free supernatants were filtered through 0.22 µm filters. The cell-free supernatants were concentrated using a Vivaspin 20 centrifugal concentrator (100 kDa MWCO, Sartorius AG, Germany) at 4,000×g. The concentrated cell-free supernatant containing MVs was subjected to ultracentrifugation at 300,000 × g (for 2 h at 4 °C), and then the pellet was resuspended in 50% (w/v) OptiPrep™ Density Gradient Medium (Sigma Aldrich, USA). MVs were further purified by density gradient centrifugation (0–50% iodixanol) at 150,000×g (for 16 h at 4 °C) using Optima XE-90 (Beckman Coulter USA) equipped with Swinging bucket SW40Ti rotor (6). MV-containing fractions were collected and then centrifuged again (at 300,000 × g for 2 h at 4 °C). The final MV pellets were resuspended in a storage buffer (500 µL, 10 mM Tris-HCl, pH 7.5). Following centrifugation, fractions corresponding to 20–35% iodixanol, including the whitish MV-enriched bands observed at 20–25%, were subjected to Western blot analysis using OmpA (OM), DsbA–HiBiT (periplasm), TatC–3×FLAG (IM), and GAPDH (cytoplasm) as subcellular markers **(Fig. S5c)**. The purified MV fractions were pooled and concentrated for subsequent analyses. The concentrated MVs were analyzed for particle size distribution using a NanoSight LM10 (Malvern Panalytical, UK). DNA and protein concentrations were measured using a NanoDrop spectrophotometer and Bradford assay, respectively. Additionally, the purified MVs were subjected to native transmission electron microscopy (TEM) to examine vesicle morphology and membrane structures, as well as proteomic analysis **(Dataset 1, Table S2).**

### Bocillin binding assay

WT strain cells cultured in LB or LB supplemented with β-lactam antibiotics were harvested and washed with PBS (58). The washed cells were incubated with fluorescent penicillin Bocillin-FL (BOCILLIN™ FL Penicillin, Sodium Salt, Invitrogen, USA) at a final concentration of 5 µg/mL for 30 min at 37°C. After labeling, cells were washed to remove unbound Bocillin-FL, and membrane fractions containing PBPs were subsequently isolated by centrifugation. Bocillin-FL–labeled PBPs were separated on 12% SDS–polyacrylamide gels (Bio-Rad Laboratories) and visualized using a Typhoon FLA 9500 (GE Healthcare Bio-Sciences AB, Uppsala, Sweden) (excitation at 488 nm and emission at 530 nm). Band intensities were quantified using ImageJ software (National Institutes of Health, USA).

### Western blot analysis

Bacterial cells cultured in LB or LB supplemented with LEX were harvested and disrupted by tip sonication (6,26,31). Depending on the experiment, crude lysates, membrane-enriched fractions, or cytosolic fractions were prepared and used for immunoblot analysis. Protein samples were separated by SDS–PAGE and then transferred onto polyvinylidene difluoride membranes (Thermo Fisher Scientific, USA). After transfer, membranes were blocked overnight at 4°C in TBS-T containing 2.7% skim milk. The blocked membranes were incubated for 2 h at 4°C with the appropriate primary antibodies: rabbit monoclonal anti-FLAG antibody (#ab205606, Abcam; 1:5,000), rabbit anti-OmpA antibody (#CSB-PA387720ZA01AUD, CUSABIO; 1:10,000), rabbit anti-Lpp antibody (#MBS7148276, MyBioSource; 1:10,000), mouse anti-HiBiT Monoclonal Antibody (#N7200, Promega; 1:1,000), or mouse monoclonal anti-GAPDH antibody (#orb627236, Biorbyt; 1:10,000). Membranes were then washed and incubated for 1 h at room temperature with HRP-conjugated secondary antibodies, including goat anti-rabbit IgG (#170-6515, Bio-Rad; 1:10,000) or goat anti-mouse IgG (H+L) (#31430, Thermo Fisher Scientific; 1:10,000). Immunoreactive signals were developed using WEST-SAVESTAR reagent (Republic of Korea) according to the manufacturer’s instructions. After a 1-min incubation at room temperature, chemiluminescent signals were immediately captured with an iBright FL1500 imaging system (Thermo Fisher Scientific, USA). Band intensities were quantified using ImageJ software, and data were obtained from three independent biological replicates.

### qRT-PCR

Total RNA was isolated from bacterial cells using the RNeasy Mini Kit (Qiagen, Germany), following the manufacturer’s protocol with minor modifications from previously reported qRT-PCR procedures (24,59). RNA concentration and purity were determined using a NanoPhotometer N60 (Implen, USA). For each sample, 1 µg of total RNA was used for cDNA synthesis. Quantitative real-time PCR was performed using the primers listed in **Table S8** on a QuantStudio 3 Real-Time PCR System (Thermo Fisher Scientific, USA). The expression level of each target gene was normalized to 16S rDNA, and relative transcript abundance was calculated from at least three independently grown cultures. Each qRT-PCR assay was performed in technical triplicate.

### Observation of cellular morphology using microscopy

Cell morphology was examined by fluorescence microscopy using a modified staining procedure based on a previously described method (57,58). WT and mutant strains were grown in LB or LB supplemented with LEX, and samples were collected at the indicated time points between 2 and 6 h. To monitor active peptidoglycan incorporation, bacterial cultures were adjusted to 1 × 10^6^ CFU/mL and incubated with the fluorescent D-amino acid HADA [7-hydroxycoumarin-amino-D-alanine; 100 µM] (R&D Systems, USA) in LB broth. After HADA labeling, cells were washed and resuspended in 1 mL of PBS. Membranes were then stained with the lipophilic dye FM4-64 at a final concentration of 1 µg/mL for 30 min at 37°C. Stained cells were washed again with PBS and imaged using confocal laser scanning microscopy (CLSM; LSM 700 or LSM 900, Carl Zeiss, Germany). HADA, GFP, and FM4-64 signals were detected using excitation wavelengths of 405, 485, and 515 nm and emission wavelengths of 490, 530, and 645 nm, respectively. Cell length and width were measured using ZEN 3.3 blue edition software.

### FRAP assay

A signal peptide of *dsbA* gene–sfGFP (SP*_dsbA_*–sfGFP) fusion construction was cloned into the IPTG-inducible pSEVA234 vector and transformed into *S.* Typhimurium WT and Δ*dacA* derivative strains (26,28). These strains were cultured in LB or LB supplemented with LEX at 37 °C for 6 h, after which the cultures were shifted to 26 °C and induced with IPTG (0.02 mM) for 30 min. Cells were washed three times with PBS and mounted on agar pads for FRAP assay. FRAP imaging was performed on a CLSM equipped with a 488 nm laser. A region of interest (ROI; ∼ 0.8 µm diameter) within the periplasmic region was photobleached using 25% laser power for a single bleach pulse, followed by time-lapse acquisition every 10 s for 180 s. Fluorescence recovery was recorded using low-intensity 488 nm excitation to minimize acquisition bleaching. Images were processed and normalized using a ZEN 3.3 (blue edition) software.

Periplasmic extracts were prepared by osmotic shock (54). Cell cultures were centrifuged at 4°C and washed three times with Tris-HCl buffer (10 mM, pH 8.0). The cell pellet was resuspended and incubated in a plasmolysis buffer (30 mM Tris-HCl, 1.0 mM EDTA, 20% [wt/vol] sucrose, pH 8.0) on ice for 10 min. After centrifugation at 4°C, the supernatant was collected and considered the periplasmic fraction.

### PG purification and LC/Q-TOF-MS analysis

Peptidoglycan purification and muropeptide analysis were performed with modifications from previously reported protocols (58,61). Briefly, WT cells were grown in 1 L of LB medium to the indicated growth phase and harvested for cell-wall preparation. After washing, cell pellets were boiled in SDS at 100°C for 20 min to remove membrane-associated components. Nucleic acids and proteins were subsequently degraded by sequential enzymatic treatment with DNase/RNase solution [15 µg/mL DNase and 60 µg/mL RNase in 0.1 M Tris-HCl buffer, pH 6.8] and trypsin solution [50 µg/mL trypsin in 0.1 M Tris-HCl]. The remaining cell-wall material was further subjected to acid treatment and extensive washing to obtain purified PG. Purified PG was digested with lysozyme at 3 mg/mL in 12.5 mM sodium phosphate buffer (pH 4.9) for 16 h at 37°C. The resulting muropeptides were analyzed by LC-Q/TOF-MS using an Agilent 1290 Infinity II LC system coupled to an AB SCIEX Q-TOF 5600+ mass spectrometer. Separation was performed on a reverse-phase Acquity LC BEH C18 column (1.7 µm, 2.1 × 100 mm; Waters) at 30°C with a flow rate of 0.3 mL/min. Mass spectra were acquired over an m/z range of 50–2000. The mobile phases consisted of solvent A, 0.1% formic acid in distilled water, and solvent B, 0.1% formic acid in acetonitrile. The chromatographic gradient was run from 100% solvent A to 100% solvent B over 60 min, followed by 5 min at 100% solvent B and re-equilibration to the initial condition for 10 min. Muropeptide peaks were assigned based on their m/z values and comparison with previously reported PG profiles.

### In vivo protein-cross-linking and immunoprecipitation of PBP5 complex

PBP5-associated protein complexes were isolated by DSS-mediated in vivo cross-linking followed by anti-FLAG immunoprecipitation, with modifications from a previously described *Salmonella* cross-linking approach (29). Overnight cultures of the chromosomally FLAG-tagged PBP5 strain were diluted 1:100 into 300 mL of LB broth and grown at 37°C for 4 h in the presence or absence of LEX. Cells were harvested, washed once with ice-cold PBS (pH 8.0), and resuspended in 2 mL of PBS. For in vivo cross-linking, disuccinimidyl suberate (DSS; Thermo Fisher Scientific, USA) was added to a final concentration of 2.5 mM. Samples were incubated for 30 min at room temperature, and the reaction was stopped by adding Tris-HCl (pH 7.5) to a final concentration of 20 mM for 15 min. Cross-linked cells were washed three times with 20 mM Tris-buffered saline (pH 7.5) and resuspended in 50 mM Tris-HCl (pH 7.0) containing DNase I (60 µg/mL) and protease inhibitor cocktail (GenDEPOT, USA). Cells were disrupted by tip sonication, and unbroken cells were removed by centrifugation at 5,000 × g for 10 min at 4°C. The clarified lysate was ultracentrifuged at 160,000 × g for 2 h at 4°C to collect the envelope fraction. The envelope pellet was solubilized in 500 µL of NP-40 buffer [150 mM NaCl, 1% (v/v) Nonidet NP-40, and 50 mM Tris-HCl, pH 8.0] for 30 min at 4°C with gentle rotation at 70 rpm. Insoluble material was removed by centrifugation at 20,000 × g for 1 h at 4°C, and the resulting supernatant containing solubilized cross-linked protein complexes was used for immunoprecipitation. Anti-FLAG antibody was added to the solubilized membrane lysate and incubated for 2 h at 4°C with rotation at 70 rpm. Dynabeads Protein A (1.5 mg; 50 µL; Invitrogen, USA) were then added, and the samples were incubated overnight at 4°C with rotation to capture antibody-bound PBP5 complexes. Beads were collected using a magnetic rack, washed three times with NP-40 buffer, and eluted with 40 µL of Laemmli sample buffer. The eluted material was used for Western blotting and LC/MS-based analysis of PBP5-associated complexes **(dataset 2).**

### In vivo protein-protein interactions using BACTH system

The bacterial adenylate cyclase two-hybrid (BACTH) system was used to test the interaction between PBP5* and ACP, with modifications from a previously described procedure (31). The dacA coding sequence lacking the signal peptide region was amplified and inserted into pUT18C and pUT18 to generate T18-fusion constructs. The acpP coding sequence was cloned into pKNT25 and pKT25 to generate T25-fusion constructs. All plasmids and primers used for BACTH analysis are listed in **Table S9**. The resulting plasmids were first constructed in E. coli DH5a, and the inserted regions were verified by PCR and sequencing. Pairwise combinations of the T18- and T25-fusion plasmids were then co-transformed into the *cyaA*-deficient reporter strain *E. coli* BTH101. Transformants were selected on LB agar plates containing ampicillin (100 µg/mL), kanamycin (50 µg/mL), IPTG (40 µg/mL), and X-Gal (0.5 mM). Plates were incubated at 37°C for 24 h, and protein-protein interaction was assessed based on colony color. Blue colonies were interpreted as positive interactions. The β-galactosidase activity of cells grown in LB broth was quantitatively assessed using a β-Gal Assay Kit (Invitrogen, USA) according to the manufacturer’s instructions with minor modifications (31). For each reaction, 10 µL of cell lysate (200 µg/mL total protein) was mixed with 50 µL of 1×cleavage buffer containing β-mercaptoethanol and 17 µL of ortho-nitrophenyl-β-D-galactopyranoside (ONPG; 4 mg/mL). After incubation at 37°C for 30 min, the reaction was terminated by adding 125 µL of stop buffer.Specific activity was calculated based on the amount of ONPG hydrolyzed, reaction time, and total protein content. β-Galactosidase activity was quantified as nmol/min/mg protein.

### Overexpression and purification of PBP5 and ACP

Genomic DNA was extracted from the *S.* Typhimurium 14028s strain to amplify the *dacA* gene (without the signal peptide and anchoring region) and the *acpP* gene using the Wizard® Genomic DNA Purification Kit (Promega) according to the manufacturer’s protocol. The amplified sequences were cloned into pET28b(+) for recombinant protein overproduction. The resulting plasmids were transformed into *E. coli* BL21 (DE3). Overnight cultures were diluted into LB broth supplemented with kanamycin (50 µg/mL) and incubated at 37°C. After 3 h, gene expression was induced by adding IPTG (0.5 mM), followed by incubation for 16 h at 20°C. Cells were harvested by centrifugation (4,000 × g) and washed three times with PBS. The cell pellets were resuspended in a lysis buffer (Tris-HCl, pH 7.6, containing 25 µg/mL protease inhibitor) and lysed by tip sonication. The lysates were centrifuged, and the supernatants were collected. Both PBP5^#^ and ACP were purified using the HisPur™ Ni–NTA Purification Kit (Thermo Fisher Scientific, USA) (62). After equilibration with an equilibration buffer (PBS containing 10 mM imidazole), the clarified lysates were loaded onto the column and allowed to bind for 1 h. Unbound proteins were removed by washing five times with a washing buffer (PBS containing 25 mM imidazole). The bound proteins were eluted using an elution buffer (PBS containing 250 mM imidazole). To remove excess imidazole, the purified PBP5-His and ACP-His proteins were buffer-exchanged using Amicon Ultra centrifugal filters (3 kDa or 10 kDa cutoff; Merck KGaA, Germany) into either MST buffer (25 mM HEPES, 150 mM NaCl, 0.05% Tween 20, pH 7.5) or reaction buffer (25 mM Tris-HCl, 10 mM NaCl, pH 7.5). For ACP, the His-tag was removed by thrombin digestion (1,000 U, MP Biomedical, USA) at 4°C overnight prior to use.

### MST-based in vitro protein-protein interaction

Direct interaction between PBP5^#^ and ACP was validated using MST on a Monolith NT.115 instrument (NanoTemper Technologies, Germany) (32,33). PBP5^#^ (400 nM) was fluorescently labeled with a histidine-reactive dye (MO-L018 Tris-NTA) according to the manufacturer’s instructions. The labeled protein was diluted to an appropriate working concentration, and a serial dilution of ACP (32.5-19,000 nM) was prepared. Capillary scans were performed prior to each measurement to ensure consistent fluorescence counts and to exclude potential artifacts arising from pipetting errors, protein adsorption, or dilution effects. MST measurements were conducted at 40% LED power and 50% MST power using the red fluorescence filter. Data were analyzed using NT Analysis software (version 1.5.41). Dissociation constants (K_d_) were calculated based on the thermophoretic response.

### In silico docking and PBP5-ACP binding prediction

A complete PBP5 and ACP model for *Salmonella enterica subsp. enterica serovar* Typhimurium str. 14028s was generated using AlphaFold3, and protein-protein docking was conducted using HADDOCK (35,36). PBP5-ACP complexes were visualized using ChimeraX. The promoter regions, transcription factors, and Nucleoid-associated protein (NAP)-binding sites in the upstream region of *dacA* were predicted using the web-based tools BPROM (http://www.softberry.com) and P2RP (http://www.p2rp.org) (24).

### OM and IM separation for LC/Q-TOF-MS–based membrane proteomics

Separation of the OM and IM was performed as previously described (6,63,64). WT cells grown in LB broth (1 L) at 37 °C for 6 h was harvested by centrifugation at 7,000 × g (for 10 min at 4 °C). The pellet was resuspended in 10 mM Tris-HCl (pH 7.5) with 0.5 M sucrose and lysozyme (10 mg/mL) solution (12.5 mL) and then was incubated 10 min. Subsequently, 1.5 mM EDTA solution (12.5 mL) was added, and the mixture was stirred for 10 min. The periplasmic fraction was collected by centrifugation at 11,000 ×g for 10 min at 4 °C. The remaining pellet was resuspended in 25 mL of 0.2 M sucrose, 10 mM Tris-HCl (pH 7.5), and 2 mM MgCl2 containing 1 µL each of RNase/DNase mix and protease inhibitor cocktail. Harvested cells were lysed by sonication (2 s on / 4 s off cycles, 1 min total, 40% amplitude). Unbroken cells were removed by centrifugation at 6,169 × g for 10 min at 4 °C, and membranes were collected by ultracentrifugation at 184,500 × g for 2 h at 4 °C. The membrane pellet was resuspended in 1 mL of 20% (w/v) sucrose, 1 mM EDTA, and 1 mM Tris-HCl (pH 7.5). Membranes were further separated using a sucrose density gradient (20%, 45%, and 73%) and ultracentrifuged for 23 h at 288,000 × g. The IM and OM were collected, washed with 10 mM Tris-HCl (pH 7.5), and pelleted again by centrifugation (at 184,500 ×g for 1 h) **(Fig. S26a-b)**. To verify the successful separation of OM and IM fractions, an NADH dehydrogenase activity assay was performed **(Fig. S26c-d)** (6). Protein concentrations of each membrane fraction were normalized to 200 µg/mL, and NADH (10 mg/mL) was used as the substrate. Enzyme activity was monitored by measuring the decrease in absorbance at 340 nm for 5 min. The recovered OM and IM fractions were subsequently subjected to LC-MS-based proteomic analyses to characterize their protein compositions **(Dataset 3).**

### Cryo-electron tomography sample preparation and data acquisition

For cryo-TEM and tomogram, WT and *AdacA* strain cells were grown overnight in LB medium. Overnight cultures were diluted (1/100) and incubated for 6 h in LB or LB with LEX conditions. Cells were harvested by centrifugation (4,000 × g, 10 min) and washed three times using PBS. Bacterial cells were applied (3 µL) onto glow-discharged Quantifoil R1.2/1.3 holey carbon copper grids (200 mesh, Quantifoil Micro Tools GmbH). Prior to sample application, grids were glow discharged using a PELCO easiGlow system (Ted Pella, USA) at 15 mA for 60 s. Vitrification was performed using a Vitrobot Mark IV (Thermo Fisher Scientific, USA) operated at 4°C and 95% relative humidity. Grids were plunge-frozen into liquid ethane and stored in liquid nitrogen until imaging. Cryo-electron tomography data were acquired using a Titan Krios G2 transmission electron microscope (Thermo Fisher Scientific, USA) operated at 300 kV and equipped with a Falcon 4i direct electron counting camera and a Selectris X energy filter. Images were recorded in electron-event representation format using a 15-eV energy-selecting slit. Tilt series were acquired using Tomography 5 software (Thermo Fisher Scientific, USA) following a bidirectional tilt scheme spanning from -60° to +60° with 2° angular increments. Acquisition started at 0° and proceeded sequentially to -60°, followed by collection from +2° to +60°. Data were collected at a calibrated pixel size of 8.33 Å/pixel and a nominal defocus of approximately -5 µm. The electron dose was approximately 1.4 ē/Å² per tilt image, resulting in a cumulative dose of ∼81.7 ē/Å² per tilt series. To obtain cryo-electron tomogram data, movie frames were first aligned and motion-corrected in cryoSPARC v4.7.0 using a fractionation value of 20. Tilt series were then aligned by fiducial-less patch tracking in IMOD v5.1.2, and tomograms were reconstructed from 4 × binned tilt images using weighted back-projection with a SIRT-like filter. Three-dimensional visualization and volume rendering of the reconstructed tomograms were performed using Amira (Thermo Fisher Scientific, USA).

### HiBiT-LgBiT system-based bioluminescence assay

Because ACP is a small cytoplasmic protein, fusion to bulky fluorescent proteins such as GFP or mCherry may interfere with its translocation to the periplasm under LEX-induced envelope stress. We employed a HiBiT-LgBiT system (Nano-Glo^®^ HiBiT Extracellular Detection System, Promega, #N2420, USA)-based bioluminescence assay, in which ACP was tagged with HiBiT, an 11-amino-acid peptide tag (37). Chromosomally tagged *dsbA*-HiBiT and *acpP*-HiBiT strains were used together with plasmids designed to express LgBiT either in the cytoplasm or in the periplasm. Cytoplasmic LgBiT was expressed from pLumi-LgBiT (Addgene, #188319), in which LgBiT expression is driven by the constitutive *rpoD* promoter, whereas periplasmic LgBiT was expressed from pBAD33-*P_dsbA_*-SP*_dsbA_*-LgBiT to direct LgBiT, which was constructed in this study to direct LgBiT to the periplasm. *dsbA*-HiBiT- and *acpP*-HiBiT-tagged strains were transformed with each LgBiT plasmid to validate cytoplasmic and periplasmic LgBiT detection. For each measurement, bacterial cultures were washed with PBS and then mixed with Nano-Glo HiBiT buffer containing the Nano-Glo substrate furimazine in white 96-well plates. HiBiT-LgBiT Luminescence was measured using a luminometer (Luminoskan Ascent, Thermo Fisher Scientific, USA). After validation, pBAD33-*P_dsbA_*-SP*_dsbA_*-LgBiT was introduced into *acpP*-HiBiT-tagged WT and *dacA* variants to measure LEX-induced periplasmic accessibility of ACP by luminescence detection.

### NHS-LC-LC-biotin labeling on free-Lpp

Overnight-grown WT and *dacA* variants were inoculated into LB broth (50 mL) in the presence or absence of LEX and cultured to the indicated time point (2, 4, and 6 h). Cells were harvested and washed three times with PBS. NHS-LC-LC-biotin (Thermo Fisher Scientific, USA) dissolved in DMSO was added to a final concentration [1% (v/v)], and samples were incubated at room temperature (3 min) (42). The reaction was quenched by adding Tris-HCl (pH 7.5) to a final concentration (250 mM). Labeled cells were washed five times with PBS, followed by cell lysis using sonication. The harvested insoluble and soluble fractions were normalized based on total protein concentration, resolved by SDS-PAGE (15%), and subjected to Western blotting. LC-LC-biotin–labeled proteins were detected using HRP conjugated Streptavidin (#N100, Thermo Fisher Scientific, USA, diluted 1:10,000).

### Purification of PG-associated proteins

Overnight cultures of strains carrying 3 X FLAG-tagged OmpC in either *acpP* knockdown or non-knockdown strains were inoculated into LB broth (1L) supplemented with KM and GM, in the presence or absence of LEX. The cultures were then adjusted to a final cell density of 1×10ၭ CFU/mL prior to harvesting by centrifugation (13,000 × g, 30 min). The cell pellets were washed three times with PBS and resuspended in 1.5 mL PBS. To isolate proteins tightly associated with PG, samples were slowly mixed with 4% SDS and incubated at 80 °C for 16 h to obtain purified PG sacculi (65). The samples were then washed extensively (three times) with PBS to completely remove residual SDS. Proteins tightly associated with PG were released by incubation with lysozyme (10 mg/mL in 50 mM phosphate buffer, pH 4.9). Following centrifugation (20,000 × g, 30 min, 4 °C), the pellet containing the precipitated proteins was resuspended in 300 µL PBS, mixed with 100 µL Laemmli sample buffer, and directly subjected to Western blot analysis.

### Subcellular fractionation for Western blotting

Overnight cultures of strains carrying 3 × FLAG-tagged OmpC in either *acpP* knockdown or non-knockdown backgrounds were inoculated into LB broth supplemented with KM and GM, with or without LEX, and grown at 37°C. After 6 h, cells were harvested and washed three times with cold PBS. Washed cell pellets were resuspended in PBS containing Triton X-100 and incubated for 3 h at 4°C (64). The samples were centrifuged at 15,000 × g for 30 min at 4°C, and the supernatant and pellet were collected as the IM-enriched and OM-enriched fractions, respectively. Lipopolysaccharide (LPS) and GAPDH were used as outer membrane and cytoplasmic markers, respectively. Equal protein loading was verified by Coomassie Brilliant Blue staining **(Fig. S27)**.

### In vivo to determine PBP5 half-lives

An in vivo chloramphenicol chase assay was performed to determine the half-lives of PBP5 in the presence or absence of ACP (38,39). Overnight cultures of strains carrying FLAG-tagged PBP5 in either *acpP* knockdown or non-knockdown strains were inoculated into LB broth supplemented with KM and GM, in the presence or absence of LEX. Arabinose (50 nM) was added where indicated to induce ACP knockdown. After 6 h of incubation, CAP (200 µg/mL) was added to inhibit protein synthesis. Samples were collected at 1 h intervals for a total of 4 h. PBP5 protein levels were analyzed by Western blotting, as described above. The normalized signal at the time of CAP addition was set as 1, and relative changes in protein levels at each time point were quantified accordingly.

### Quantification and statistical analysis

Statistical analyses were performed using GraphPad Prism 11.0 (GraphPad Software).

Significant differences were determined using Student’s *t*-test (two-tailed) for comparisons between two groups. To evaluate pairwise differences in relative gene expression within each stage, statistical significance was determined using one-way ANOVA followed by Tukey’s multiple comparisons test. The *P-*values are indicated on each figure (*P < 0.05, **P < 0.01, ***P < 0.001, and ****P< 0.0001. ns, not significant).

## Data availability statement

The datasets used in this study are available in the NCBI sequence archives under the accession numbers CP001363.1 (*Salmonella enterica subsp. enterica serovar* Typhimurium str. 14028s). The proteomic data of all identified peptides and protein lists have been deposited in the ProteomeXchange Consortium (http://proteomecentral.proteomexchange.org) via the PRIDE partner repository, with the dataset identifier PXD047524 (MVs derived from LEX-treated cells), PXD074933 (PBP5-associated complex using in vivo crosslinking assays using a chromosomally FLAG-tagged PBP5 strain cultured in LB and LB with LEX), PXD047522 (IM fractions of WT strain cultured in LB and LB with LEX), and PXD047523 (OM fractions of WT strain cultured in LB and LB with LEX). Dataset 1 is available under PXD047524; Dataset 2 is available under PXD074933; Dataset 3 is available under PXD047522 and PXD047523.

## Acknowledgments

We thank a member of Juhyun Kim’s lab (Kyungpook National University) for providing the plasmids pKIKO–msfGFP/sfGFP, pCP20, and pSEVA234. We also thank a member of Jinki Yeom’s lab (Seoul National University) for providing the plasmids pKD3, pKD4, and pKD46 as well as *AphoP* strain.

## Author contributions

**Yongjun Son and Jihye Yang**: These authors contributed equally. Conceptualization, Methodology, Resources, Data curation, Software, Validation, Visualization, Formal analysis, Investigation, Writing-Original draft, Writing – Review & Editing. **Wonjae Kim**: Validation and Data curation. **Jinwon Park:** Methodology and Validation. **Hyosung Choi**: Software and Visualization. **Hyeongseop Jeong:** Validation and data curation. **Woojun Park**: Conceptualization, Methodology, Resources, Data curation, Investigation, Writing-Original draft, Writing – Review & Editing, Supervision, Project administration, and Funding acquisition.

---

**Table S1.** MIC values of β-lactam antibiotics against *S.* Typhimurium. MICs of β-lactam antibiotics were determined using *S.* Typhimurium cells cultured in LB broth in 96-well plates at 37°C. Cells were adjusted to 1 × 10^6^ cells/mL and incubated with each β-lactam antibiotic for 24 h.

| **β-lactam antibiotics** |  | **MIC (µg/mL)** |
|---|---|---|
| **Monobactam** | Aztreonam (AZT) | 0.06 |
| **Penicillin** | Ampicillin (AMP) | 64 |
|  | Amoxicillin (AMX) | 16 |
|  | Mecillinam (MEC) | 256 |
| **Cephalosporin** | Cephalexin (LEX) | 256 |
|  | Cefotaxime (CTX) | 1 |
|  | Ceftriaxone (CRO) | 1 |
| **Carbapenem** | Meropenem (MEM) | 128 |
|  | Doripenem (DOR) | 32 |

\*Aztreonam, AZT; Ampicillin, AMP; Amoxicillin, AMX; Mecillinam, MEC; Cephalexin, LEX; Cefotaxime, CTX; Ceftriaxone, CRO; Meropenem, MEM; Doripenem, DOR.

**Table S2.**
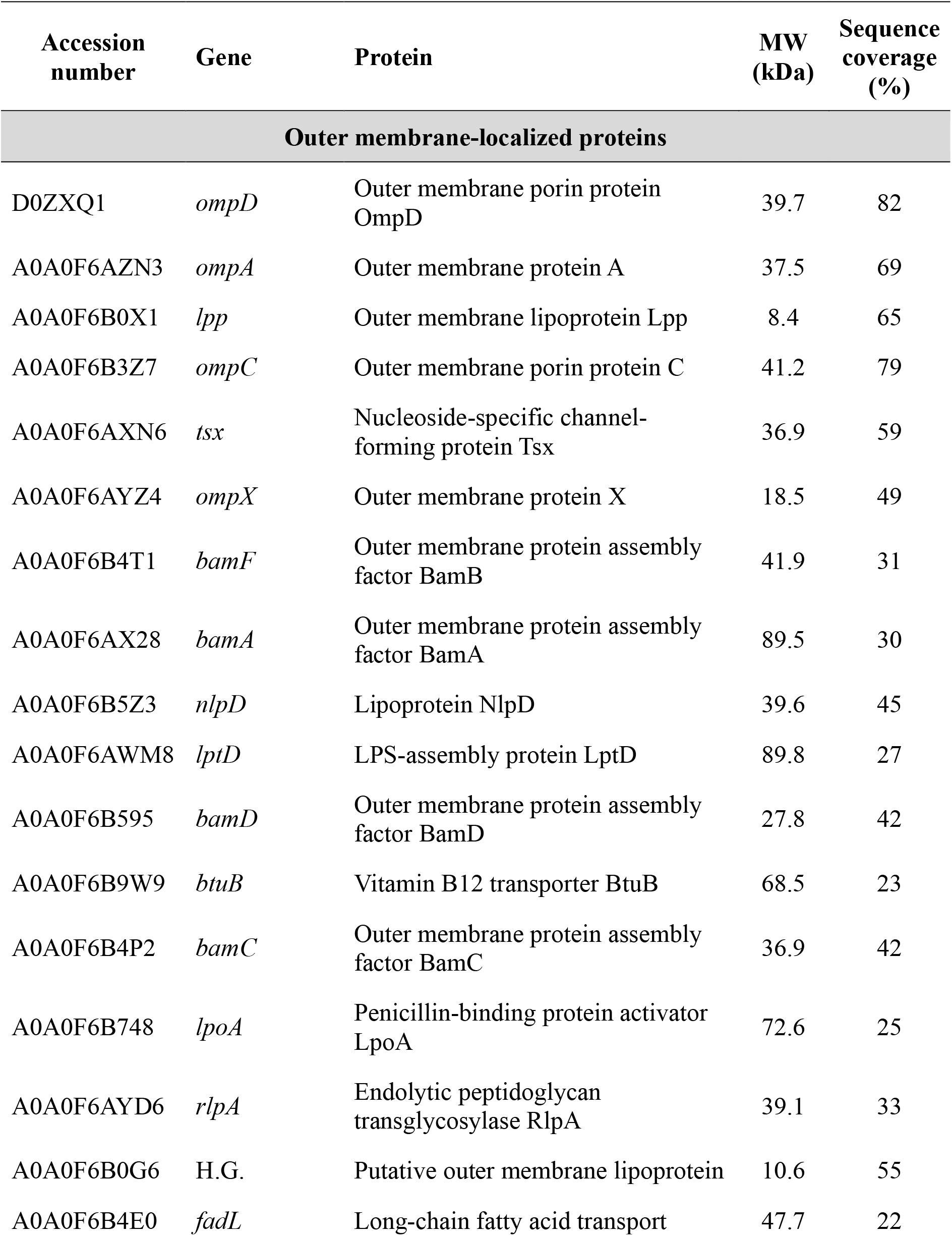
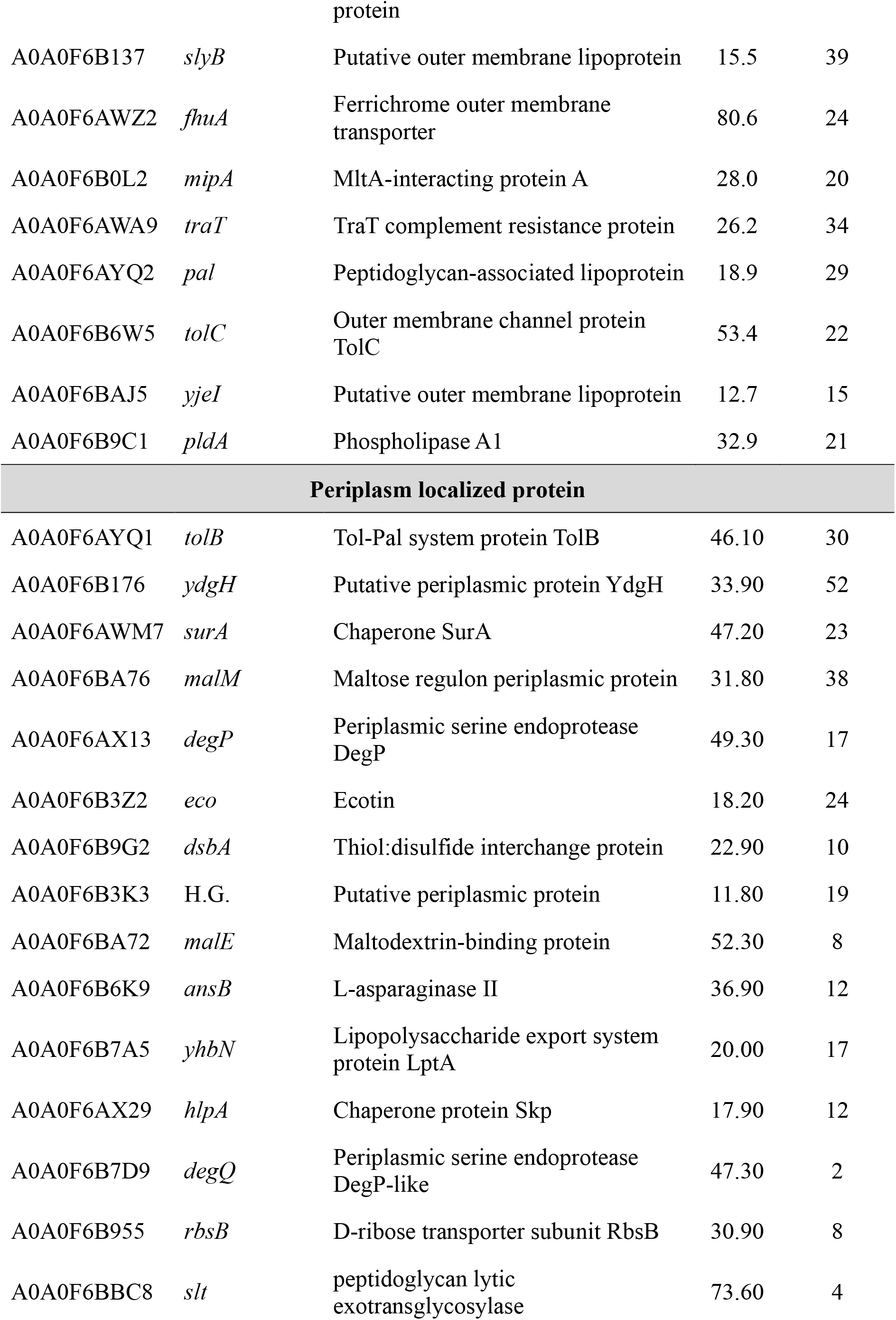
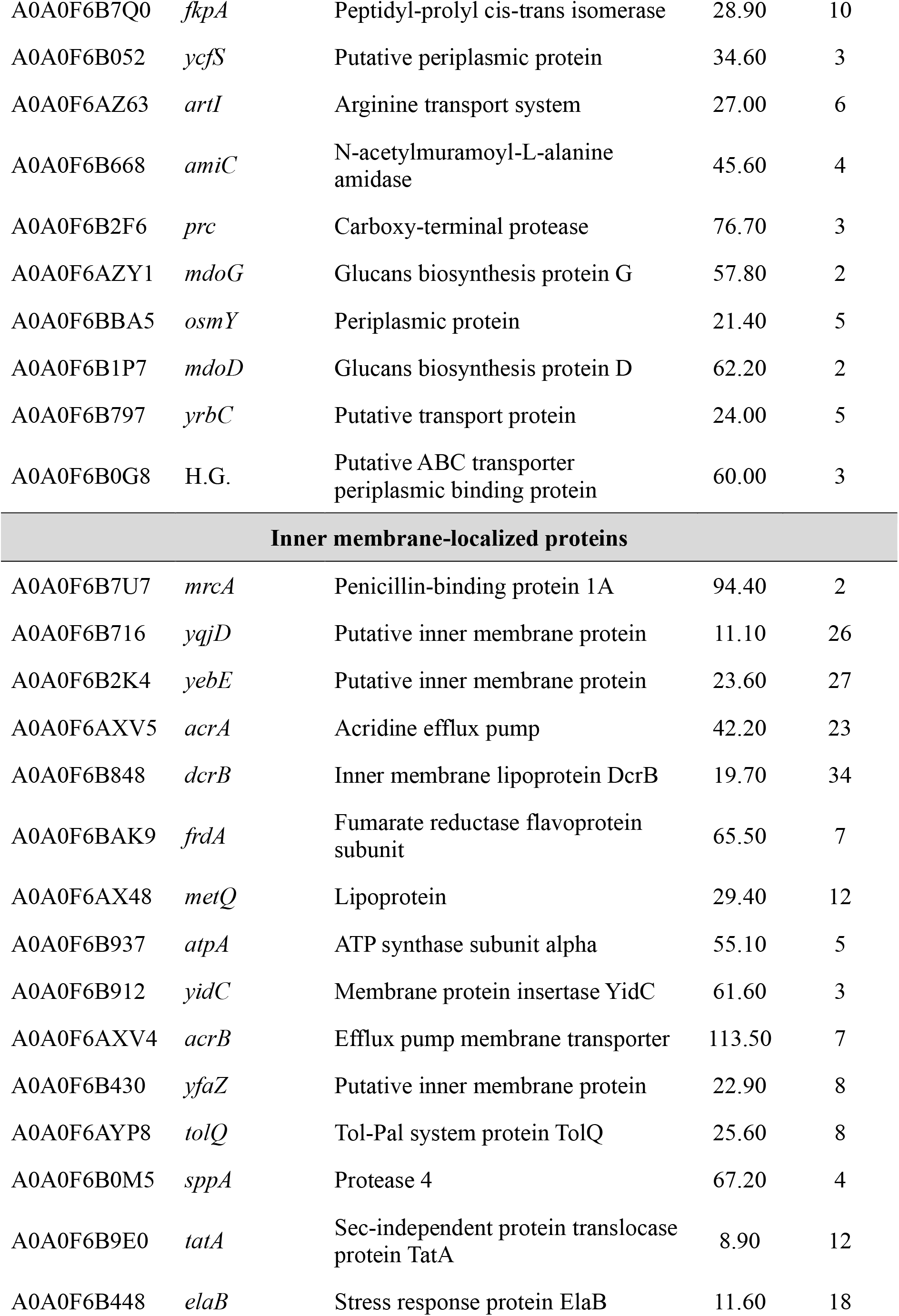
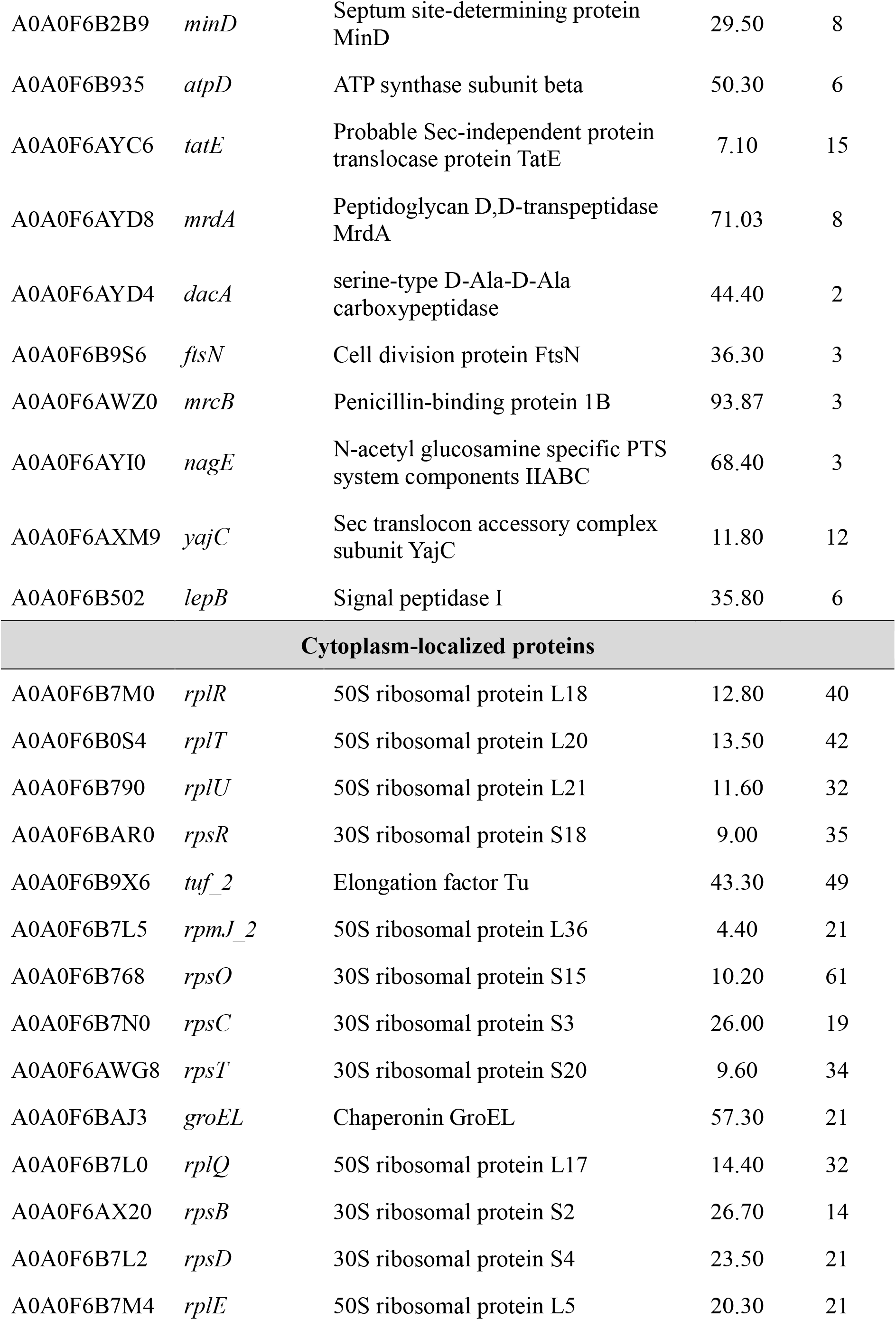
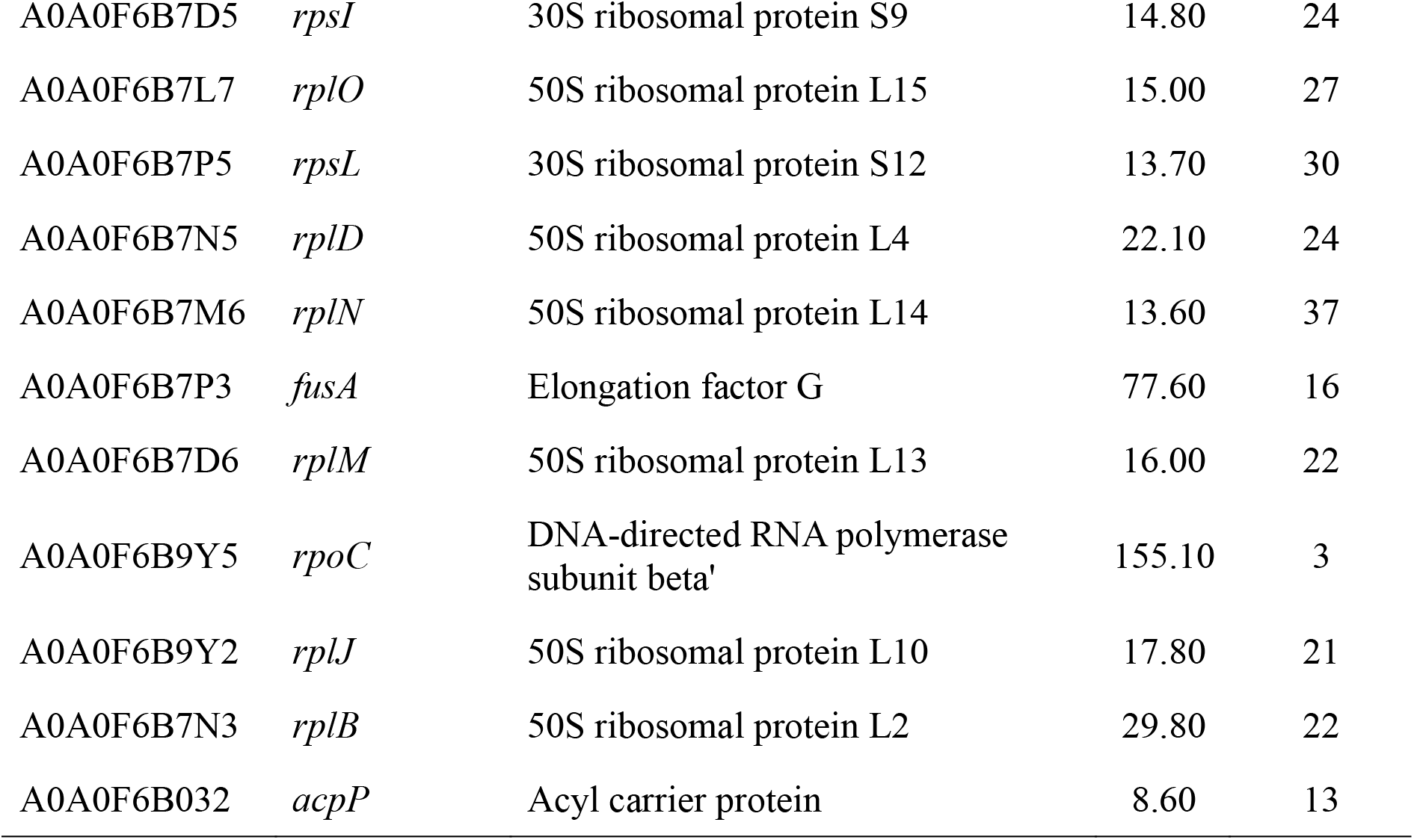
Top 25 most abundant proteins in each subcellular localization category identified in vesicles of *S.* Typhimurium 14028s under LEX stress. A total of 235 proteins identified in vesicles under LEX stress were classified into four categories based on their subcellular localization: outer membrane, periplasm, inner membrane, and cytoplasm. The top 25 proteins in each category were selected based on their abundance under LEX stress. H.G., hypothetical gene.

**Table S3.** Annotated penicillin binding proteins in the genome of *S.* Typhimurium 14028s.

| **Functional category** | **Gene** | **Protein** | **Function** | **Size (kDa)** | **a.a similarity of *E. coli* (%)** | **Accession number** |
|---|---|---|---|---|---|---|
| **Penicillin binding protein** | *mrcA* | PBP1a | glycosyltransferase/transpeptidase PBP1a | 94.4 | 92.6 | P02918 |
|  | *mrcB* | PBP1b | glycosyltransferase/transpeptidase PBP1b | 93.9 | 92.6 | P02919 |
|  | *pbpC* | PBP1c | glycosyltransferase/transpeptidase PBP1c | 85.1 | 82.4 | P76577 |
|  | *mrdA* | PBP2 | DD-transpeptidase PBP2 | 71.0 | 96.2 | P0AD65 |
|  | *mrdA*<sub>SAL</sub> | PBP2<sub>SAL</sub><sup>*</sup> | DD-transpeptidase PBP2<sub>SAL</sub> | 69.8 | 58.6 | P0AD65 |
|  | *ftsI* | PBP3 | DD-transpeptidase PBP3 | 63.8 | 96.3 | P0AD68 |
|  | *ftsI*<sub>SAL</sub> | PBP3<sub>SAL</sub><sup>#</sup> | DD-transpeptidase PBP3<sub>SAL</sub> | 60.1 | 64.4 | P0AD68 |
| **Carboxypeptidase, Endopeptidase** | *dacB* | PBP4 | DD-carboxypeptidase/endopeptidase PBP4 | 52.0 | 93.9 | P24228 |
|  | *dacA* | PBP5 | DD-carboxypeptidase PBP5 | 44.5 | 95.0 | P0AEB2 |
|  | *dacC* | PBP6a | DD-carboxypeptidase PBP6a | 43.7 | 92.2 | P08506 |
|  | *dacD* | PBP6b | DD-carboxypeptidase PBP6b | 43.1 | 87.4 | P33013 |
|  | *pbpG* | PBP7 | DD-endopeptidase PBP7 | 34.3 | 92.4 | P0AFI5 |

\*PBP2<sub>SAL</sub><sup>\*</sup>, pathogen-specific PBP2<sub>SAL</sub>; PBP3<sub>SAL</sub><sup>#</sup>, pathogen-specific PBP3<sub>SAL</sub>; a.a, amino acid .

**Table S4.** Muropeptide composition of peptidoglycan from *S.* Typhimurium cells under control and LEX stress conditions. M, D, and T indicate monomeric, dimeric, and trimeric muropeptides, respectively. M2, monomeric disaccharide-dipeptide muropeptide; M3, monomeric disaccharide-tripeptide muropeptide; M3-Gly, monomeric disaccharide-tripeptide muropeptide with glycine as the fourth stem residue; M4, monomeric disaccharide-tetrapeptide muropeptide; M5, monomeric disaccharide-pentapeptide muropeptide; D33, dimeric muropeptide composed of two tripeptide-containing disaccharide units; D43, dimeric muropeptide composed of one tetrapeptide-containing and one tripeptide-containing disaccharide units; D43_Lpp_, dimeric muropeptide composed of tetrapeptide- and tripeptide-containing disaccharide units covalently linked to Lpp; D43_Gly_, dimeric muropeptide composed of tetrapeptide- and tripeptide-containing disaccharide units, with glycine at the fourth stem position; D44_N_, dimeric muropeptide composed of two tetrapeptide-containing disaccharide units containing an anhydro-MurNAc residue; T443, trimeric muropeptide composed of two tetrapeptide-containing and one tripeptide-containing disaccharide units.

| **Peak** | **RT (min)** | **Muropeptide annotation** | **Observed *m/z*** |
|---|---|---|---|
| 1 | 1.73 | M3<sub>Gly</sub> | 926.3 |
| 2 | 1.83 | D43<sub>Lpp</sub> | 747.4 |
| 3 | 1.95 | M4 | 940.3 |
| 4 | 2.78 | D43<sub>Gly</sub> | 962.4 |
| 5 | 3.70 | T443 | 932.4 |
| 6 | 4.02 | M2 | 697.1 |
| 7 | 6.93 | D43 | 931.4 |
| 8 | 7.48 | D43 | 959.1 |
| 9 | 7.68 | D33 | 860.1 |
| 10 | 8.23 | D43 | 931.4 |
| 11 | 8.96 | D43 | 931.4 |
| 12 | 10.05 | D44<sub>N</sub> | 922.3 |

**Table S5.** Membrane proteomic identification of cytosolic proteins in *S.* Typhimurium 14028s under control and LEX stress conditions. WT strain culture were harvested at 6 h and fractionated into OM and IM fractions before proteomic analyses.

| **Accession number** | **Gene** | **Protein** | **log<sub>2</sub> fold change (LEX/Con)** |  | **Size (kDa)** | **Sequence coverage (%)** |  |
|  |  |  | **OM** | **IM** |  | **OM** | **IM** |
|---|---|---|---|---|---|---|---|
| **Fatty acid biosynthesis** |  |  |  |  |  |  |  |
| A0A0F6B032 | *acpP* | Acyl carrier protein | 1.8 | 0.6 | 8.6 | 33 | 13 |
| A0A0F6AZN0 | *fabA* | 3-hydroxydecanoyl-[acyl-carrier-protein] dehydratase | N.D | 0.2 | 19 | - | 8 |
| A0A0F6B4C5 | *fabB* | 3-oxoacyl-(Acyl carrier protein) synthase I | N.D | 0.6 | 42.3 | - | 6 |
| A0A0F6B033 | *fabF* | 3-oxoacyl-[acyl-carrier-protein] synthase 2 | N.D | 11.1 | 42.9 | - | 9 |
| A0A0F6AX31 | *fabZ* | 3-hydroxyacyl-[acyl-carrier-protein] dehydratase FabZ | 0.9 | 1.2 | 20.5 | 52 | 52 |
| **Nucleoid organization, translation machinery** |  |  |  |  |  |  |  |
| A0A0F6BA04 | *hupA* | Transcriptional regulator HU subunit alpha | 1.0 | 1.3 | 9.5 | 47 | 63 |
| A0A0F6AXT0 | *hupB* | Transcriptional regulator HU subunit beta | 0.1 | 2.1 | 9.2 | 34 | 34 |
| A0A0F6AYY9 | *dps* | DNA protection during starvation protein | 2.1 | 0.4 | 18.7 | 72 | 72 |
| A0A0F6B907 | *dnaA* | Chromosomal replication initiator protein DnaA | 5.6 | 2.3 | 52.6 | 2 | 6 |
| A0A0F6AZ95 | *infA* | Translation initiation factor IF-1 | N.D | 1.3 | 8.2 | - | 22 |
| A0A0F6B771 | *infB* | Translation initiation factor IF-2 | N.D | 1.5 | 97.3 | - | 33 |
| A0A0F6B0S2 | *infC* | Translation initiation factor IF-3 | 0.9 | 1.8 | 20.6 | 21 | 58 |
| A0A0F6AX21 | *tsf* | Elongation factor Ts | 0.4 | N.D | 30.3 | 3 | - |
| A0A0F6B9X6 | *tsf_2* | Elongation factor Tu | 2.2 | 1.1 | 43.3 | 60 | 44 |
| A0A0F6B7P3 | *fusA* | Elongation factor G | 2.9 | 0.9 | 77.6 | 7 | 11 |
| A0A0F6B503 | *lepA* | Elongation factor 4 | 3.0 | 0.3 | 66.5 | 6 | 6 |
| A0A0F6B981 | *rho* | Transcription termination factor Rho | 10.6 | 3.1 | 47 | 26 | 35 |
| A0A0F6B9X9 | *nusG* | Transcription termination/antitermination protein NusG | N.D | 17 | 20.5 | N.D | 20.5 |
| **30S ribosomal protein** |  |  |  |  |  |  |  |
| A0A0F6AZD0 | *rpsA* | 30S ribosomal protein S1 | 3.2 | 0.4 | 61.1 | 24 | 37 |
| A0A0F6AX20 | *rpsB* | 30S ribosomal protein S2 | 3.5 | 5.4 | 26.7 | 52 | 59 |
| A0A0F6B7N0 | *rpsC* | 30S ribosomal protein S3 | 6.3 | 4.1 | 26 | 58 | 29 |
| A0A0F6B7L2 | *rpsD* | 30S ribosomal protein S4 | 1.7 | 2.0 | 23.5 | 45 | 55 |
| A0A0F6B7L9 | *rpsE* | 30S ribosomal protein S5 | 3.0 | 8.6 | 17.6 | 62 | 79 |
| A0A0F6BAQ8 | *rpsF* | 30S ribosomal protein S6 | 1.6 | 1.9 | 15.2 | 21 | 66 |
| A0A0F6B7P4 | *rpsG* | 30S ribosomal protein S7 | 5.9 | 10.9 | 17.6 | 52 | 70 |
| A0A0F6B7M2 | *rpsH* | 30S ribosomal protein S8 | 1.3 | 9.3 | 14.1 | 26 | 57 |
| A0A0F6B7D5 | *rpsI* | 30S ribosomal protein S9 | 1.0 | 4.0 | 14.8 | 49 | 57 |
| A0A0F6B7N7 | *rpsJ* | 30S ribosomal protein S10 | 37.3 | 15.2 | 11.8 | 43 | 60 |
| A0A0F6B7L3 | *rpsK* | 30S ribosomal protein S11 | 7.6 | 40.9 | 13.8 | 9 | 47 |
| A0A0F6B7P5 | *rpsL* | 30S ribosomal protein S12 | 6.5 | 25.3 | 13.7 | 23 | 48 |
| A0A0F6B7L4 | *rpsM* | 30S ribosomal protein S13 | 2.8 | 5.1 | 13.2 | 54 | 68 |
| A0A0F6B7M3 | *rpsN* | 30S ribosomal protein S14 | 73.1 | 10.1 | 11.1 | 30 | 55 |
| A0A0F6B768 | *rpsO* | 30S ribosomal protein S15 | 18.4 | 2.1 | 10.2 | 34 | 70 |
| A0A0F6B5A9 | *rpsP* | 30S ribosomal protein S16 | 9.4 | 2.8 | 9.2 | 52 | 59 |
| A0A0F6B7M7 | *rpsQ* | 30S ribosomal protein S17 | 16.4 | 2.0 | 9.7 | 10 | 42 |
| A0A0F6BAR0 | *rpsR* | 30S ribosomal protein S18 | 56.8 | 3.6 | 9 | 40 | 51 |
| A0A0F6B7N2 | *rpsS* | 30S ribosomal protein S19 | 12.0 | 6.4 | 10.4 | 37 | 58 |
| A0A0F6AWG8 | *rpsT* | 30S ribosomal protein S20 | 2.6 | 1.9 | 9.6 | 38 | 44 |
| A0A0F6B6Z2 | *rpsU* | 30S ribosomal protein S21 | 34.5 | 21.6 | 8.5 | 25 | 48 |
| **50S ribosomal protein** |  |  |  |  |  |  |  |
| A0A0F6B9Y1 | *rplA* | 50S ribosomal protein L1 | 3.5 | 3.6 | 24.7 | 54 | 59 |
| A0A0F6B7N3 | *rplB* | 50S ribosomal protein L2 | 3.8 | 8.1 | 29.8 | 59 | 59 |
| A0A0F6B7N6 | *rplC* | 50S ribosomal protein L3 | 28.5 | 30.2 | 22.2 | 35 | 44 |
| A0A0F6B7N5 | *rplD* | 50S ribosomal protein L4 | 13.5 | 8.7 | 22.1 | 37 | 37 |
| A0A0F6B7M4 | *rplE* | 50S ribosomal protein L5 | 9.3 | 1.3 | 20.6 | 74 | 75 |
| A0A0F6B7M1 | *rplF* | 50S ribosomal protein L6 | 4.5 | 19.0 | 18.8 | 59 | 72 |
| A0A0F6B9Y3 | *rplL* | 50S ribosomal protein L7/L12 | 4.0 | 6.7 | 12.3 | 30 | 19 |
| A0A0F6BAR1 | *rplI* | 50S ribosomal protein L9 | 4.1 | 2.7 | 15.8 | 64 | 64 |
| A0A0F6B9Y2 | *rplJ* | 50S ribosomal protein L10 | 8.4 | 13.0 | 17.8 | 52 | 56 |
| A0A0F6B9Y0 | *rplK* | 50S ribosomal protein L11 | 5.8 | 5.2 | 14.9 | 28 | 39 |
| A0A0F6B7D6 | *rplM* | 50S ribosomal protein L13 | 5.7 | 7.6 | 16 | 55 | 63 |
| A0A0F6B7M6 | *rplN* | 50S ribosomal protein L14 | 8.0 | 8.3 | 13.6 | 45 | 45 |
| A0A0F6B7L7 | *rplO* | 50S ribosomal protein L15 | 1.8 | 1.7 | 15 | 59 | 59 |
| A0A0F6B7M9 | *rplP* | 50S ribosomal protein L16 | 2.1 | 2.0 | 15.2 | 45 | 45 |
| A0A0F6B7L0 | *rplQ* | 50S ribosomal protein L17 | 1.0 | 1.8 | 14.4 | 40 | 40 |
| A0A0F6B7M0 | *rplR* | 50S ribosomal protein L18 | 0.9 | 4.9 | 12.8 | 21 | 45 |
| A0A0F6B5A6 | *rplS* | 50S ribosomal protein L19 | 5.5 | 3.4 | 13.1 | 60 | 69 |
| A0A0F6B0S4 | *rplT* | 50S ribosomal protein L20 | 8.6 | 1.5 | 13.5 | 42 | 48 |
| A0A0F6B790 | *rplU* | 50S ribosomal protein L21 | 32.8 | 1.0 | 11.6 | 36 | 36 |
| A0A0F6B7N1 | *rplV* | 50S ribosomal protein L22 | 4.1 | 3.5 | 12.2 | 62 | 62 |
| A0A0F6B7N4 | *rplW* | 50S ribosomal protein L23 | 19.1 | 21.1 | 11.2 | 19 | 28 |
| A0A0F6B7M5 | *rplX* | 50S ribosomal protein L24 | 10.4 | 4.4 | 11.3 | 57 | 76 |
| A0A0F6B3V1 | *rplY* | 50S ribosomal protein L25 | N.D | 13.3 | 10.5 | - | 53 |
| A0A0F6B789 | *rpmA* | 50S ribosomal protein L27 | 13.8 | 6.0 | 9.1 | 21 | 25 |
| A0A0F6B8M6 | *rpmB* | 50S ribosomal protein L28 | 27.0 | 3.9 | 9 | 45 | 36 |
| A0A0F6B7M8 | *rpmC* | 50S ribosomal protein L29 | 5.0 | 2.7 | 7.6 | 46 | 49 |
| A0A0F6B7L8 | *rpmD* | 50S ribosomal protein L30 | 7.8 | 13.9 | 6.5 | 19 | 19 |
| A0A0F6B9S9 | *rpmE* | 50S ribosomal protein L31 | 30.6 | N.D | 7.7 | 23 | - |
| A0A0F6B027 | *rpmF* | 50S ribosomal protein L32 | N.D | 12.4 | 6.4 | - | 18 |
| A0A0F6B8M5 | *rpmG* | 50S ribosomal protein L33 | 42.9 | 35.3 | 6.4 | 27 | 56 |

\*OM, outer membrane; IM, inner membrane; N.D, not detected.

**Table S6.**
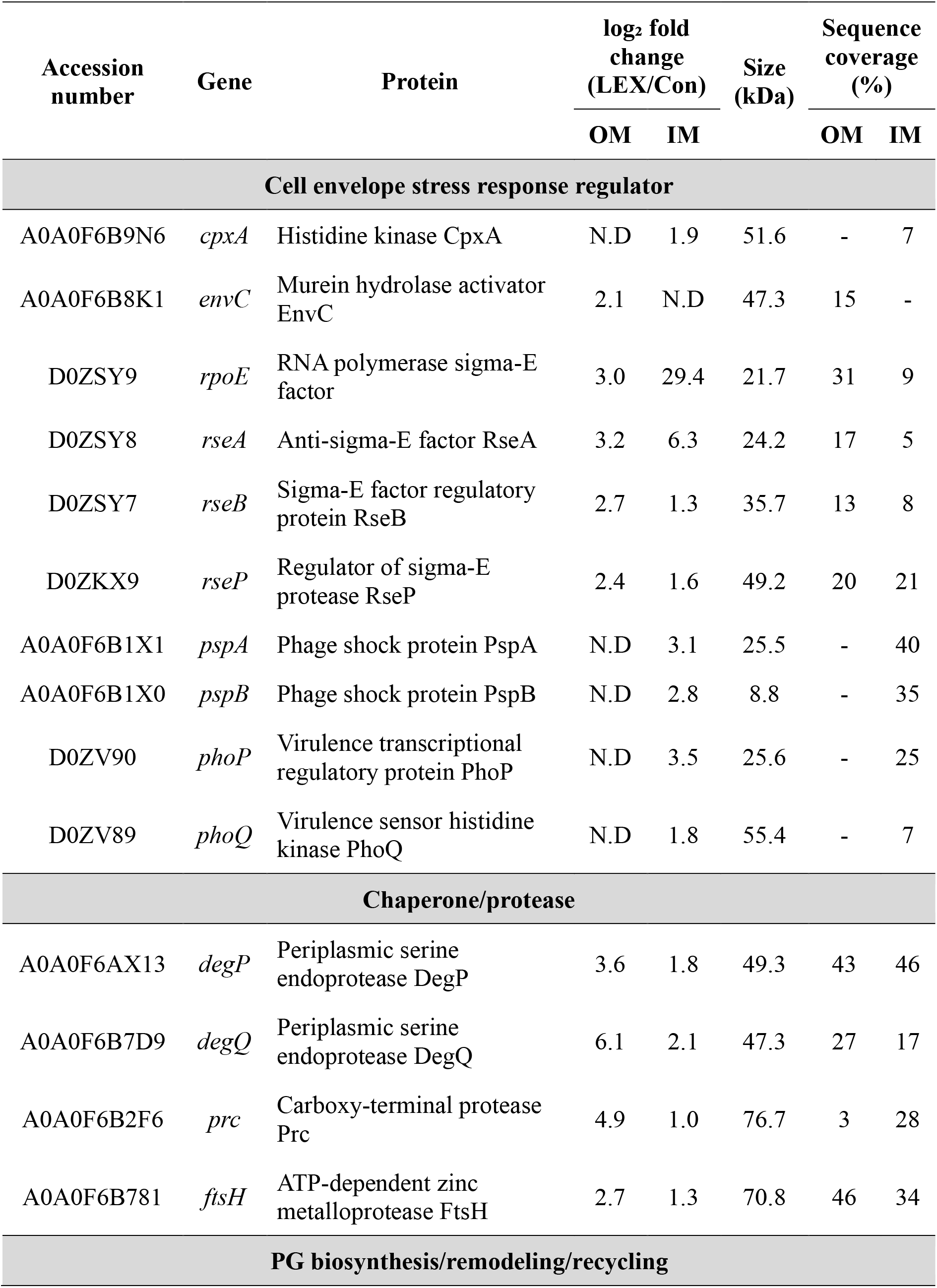
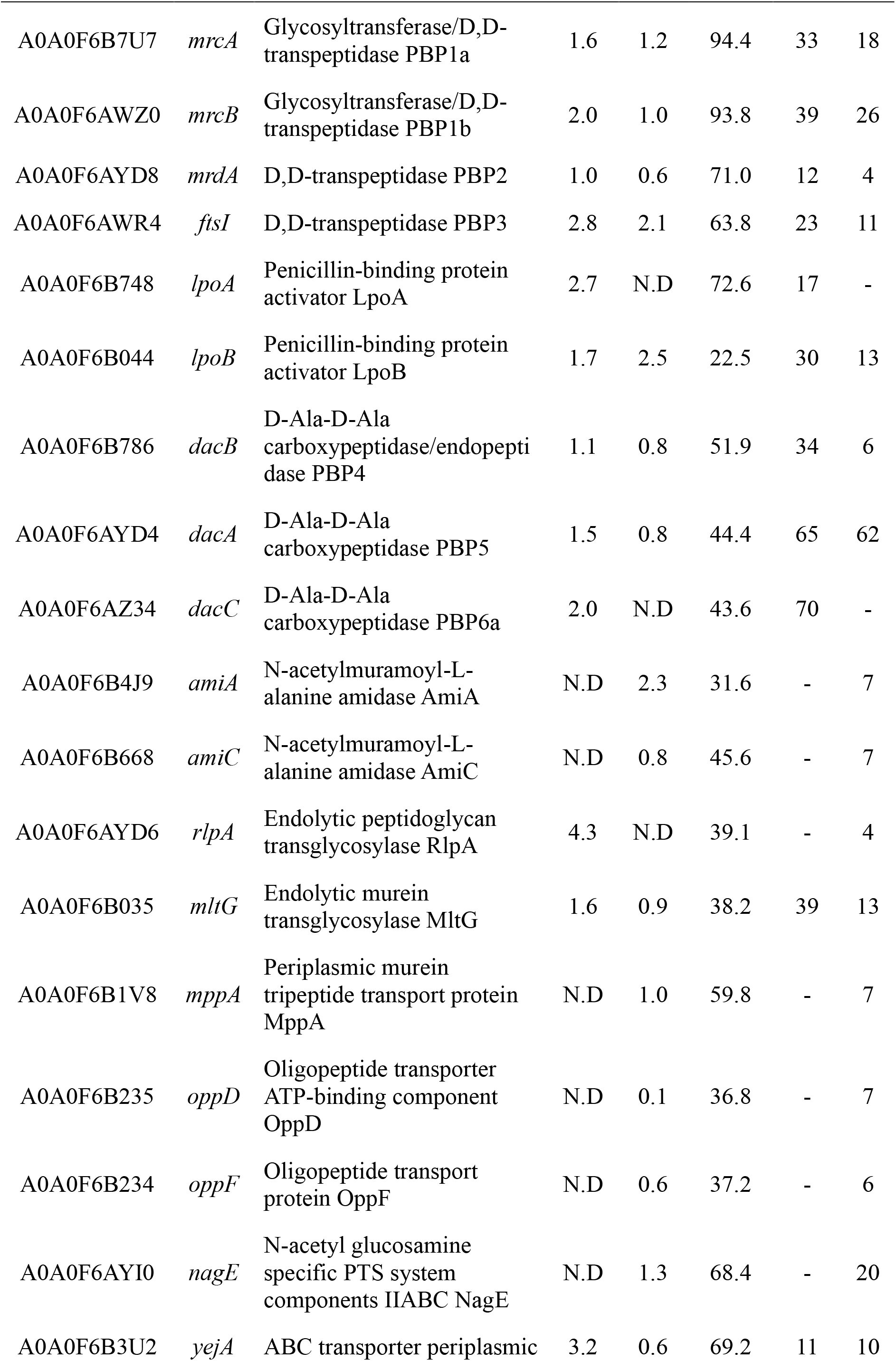
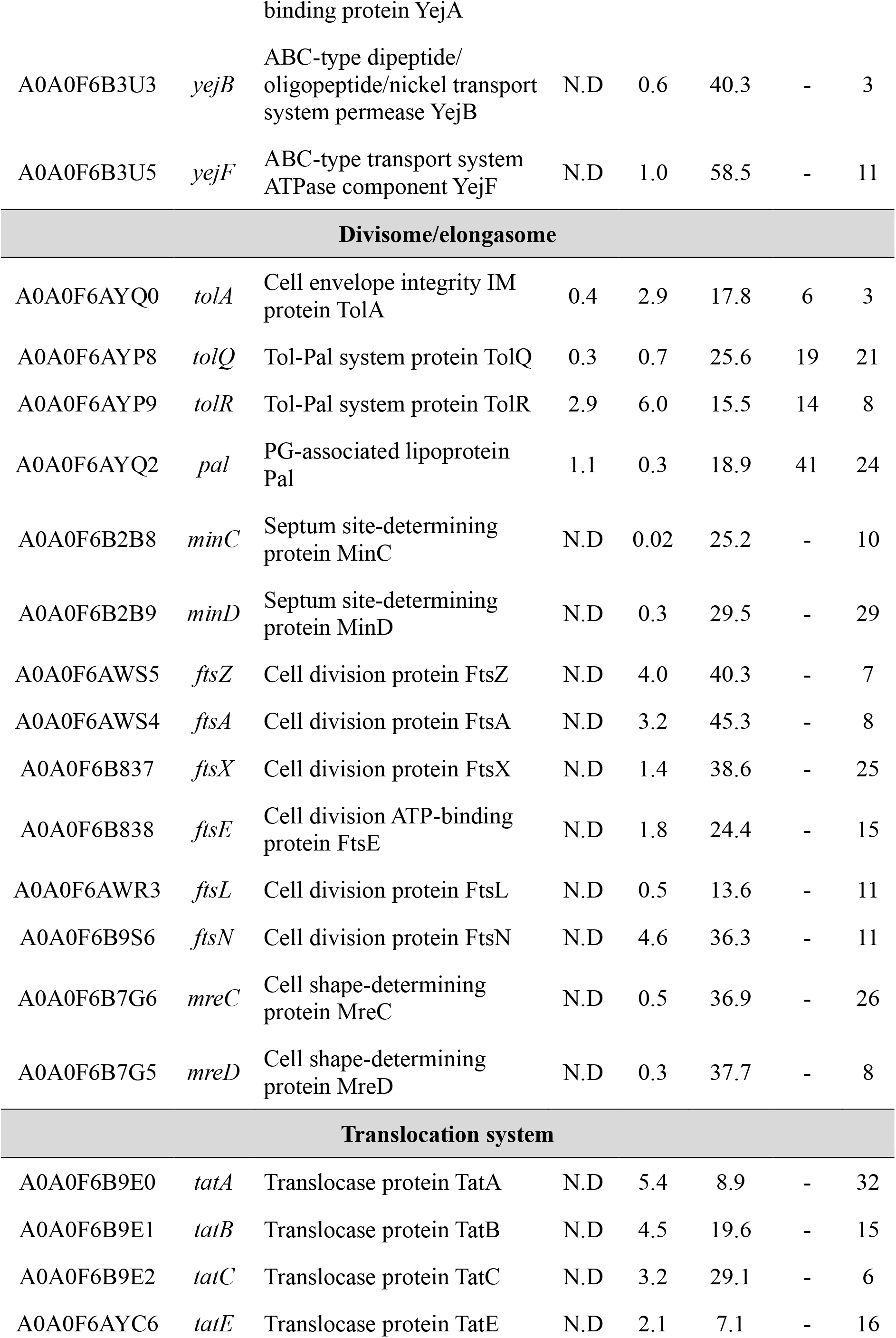
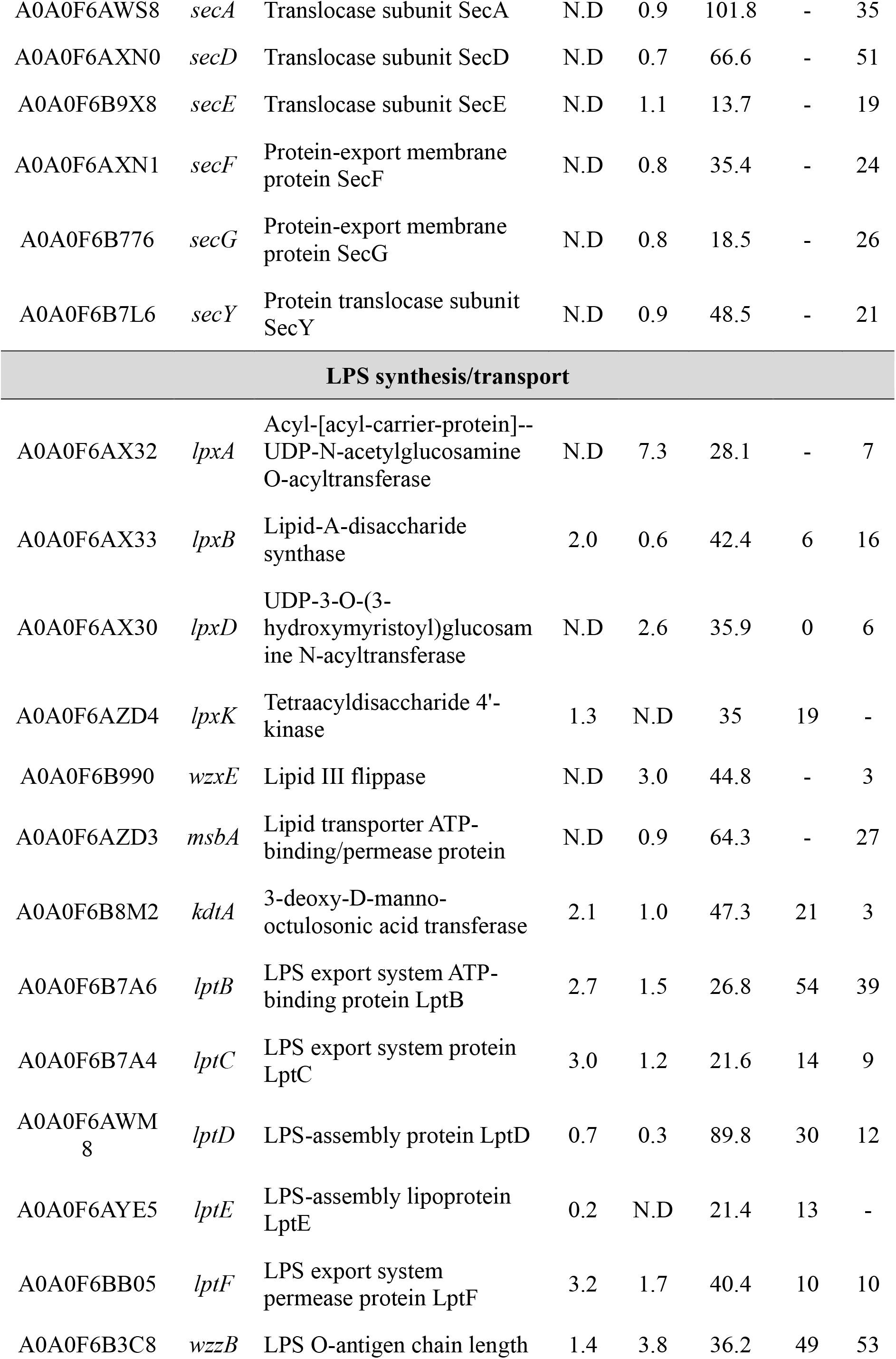
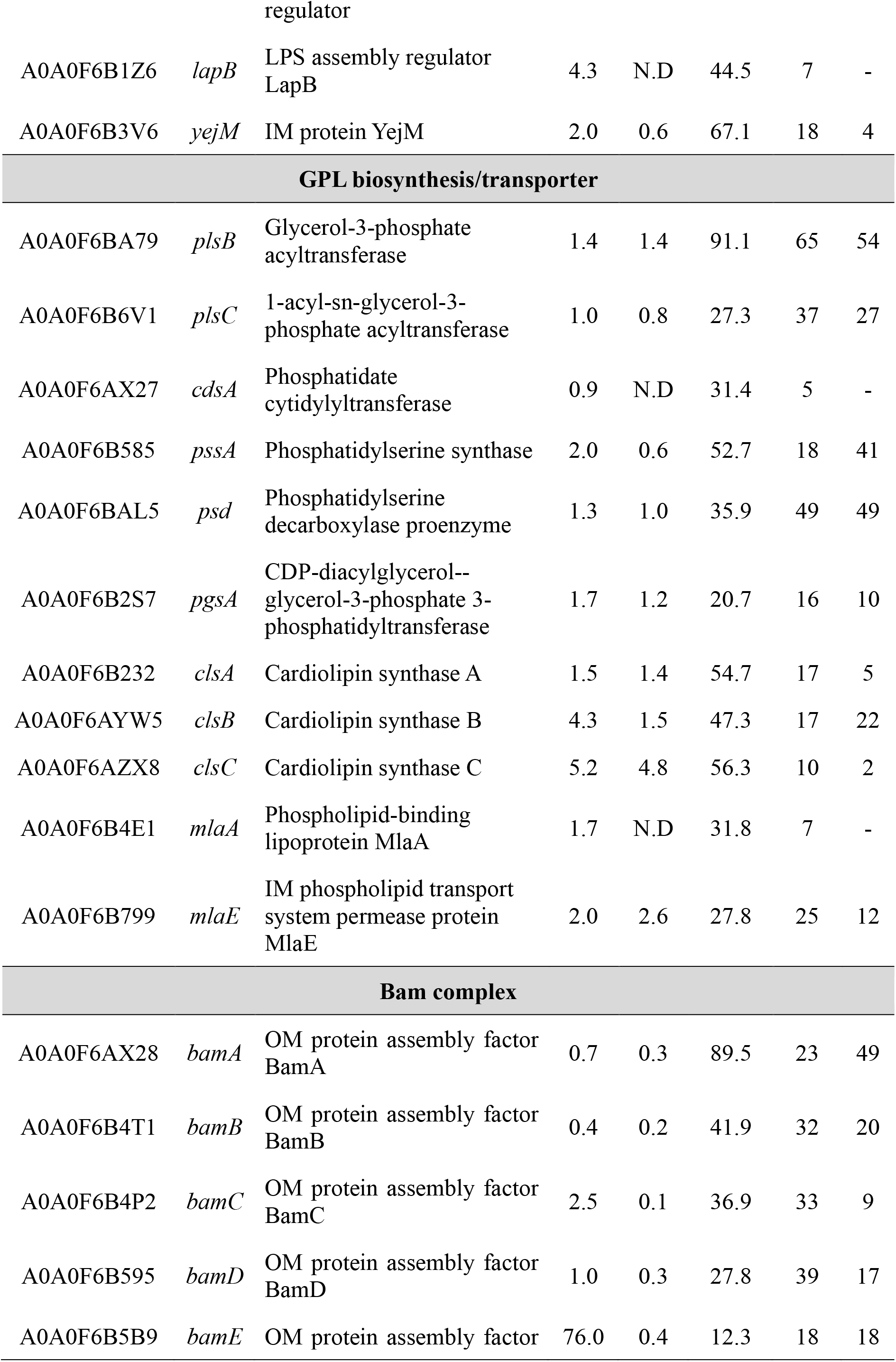
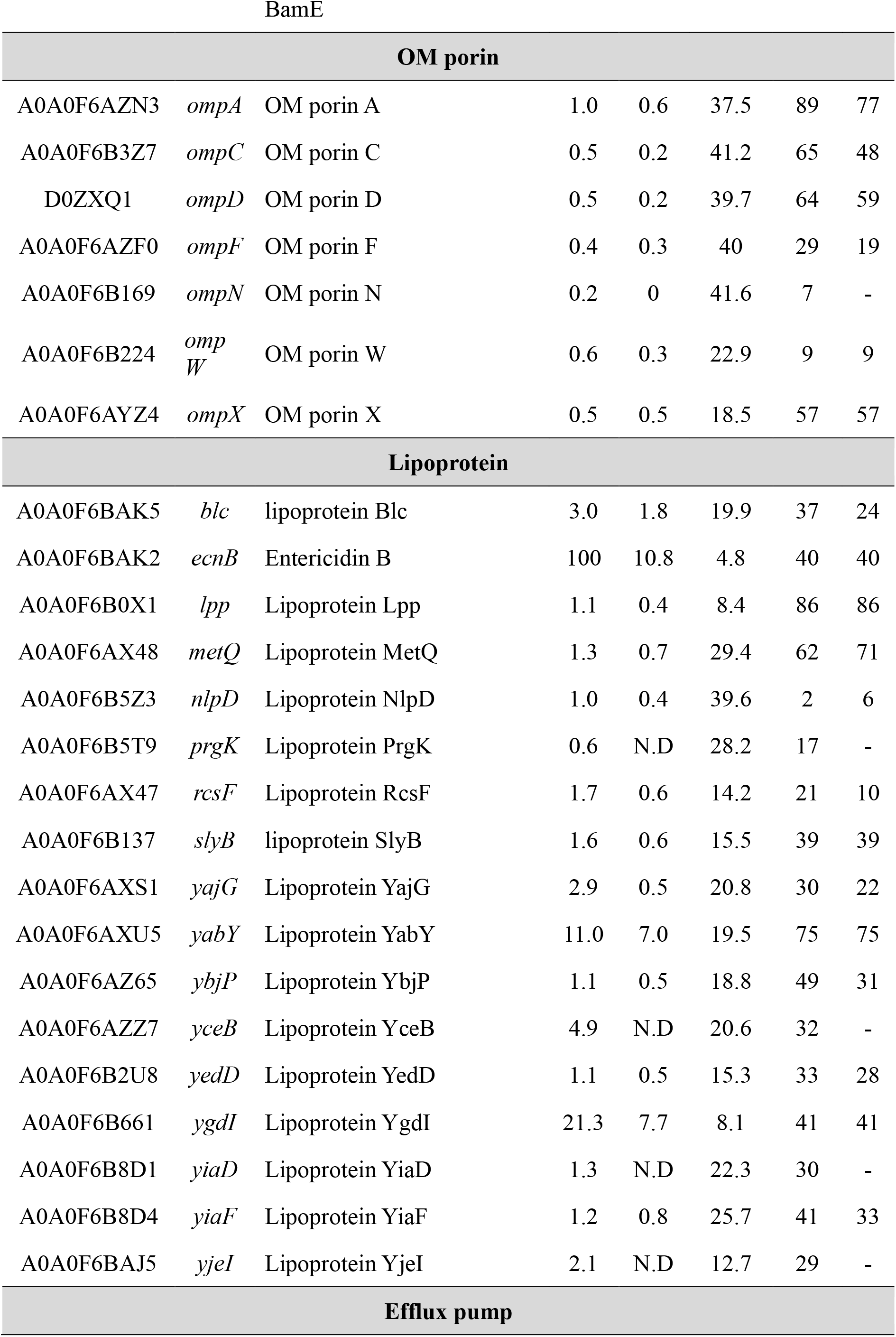
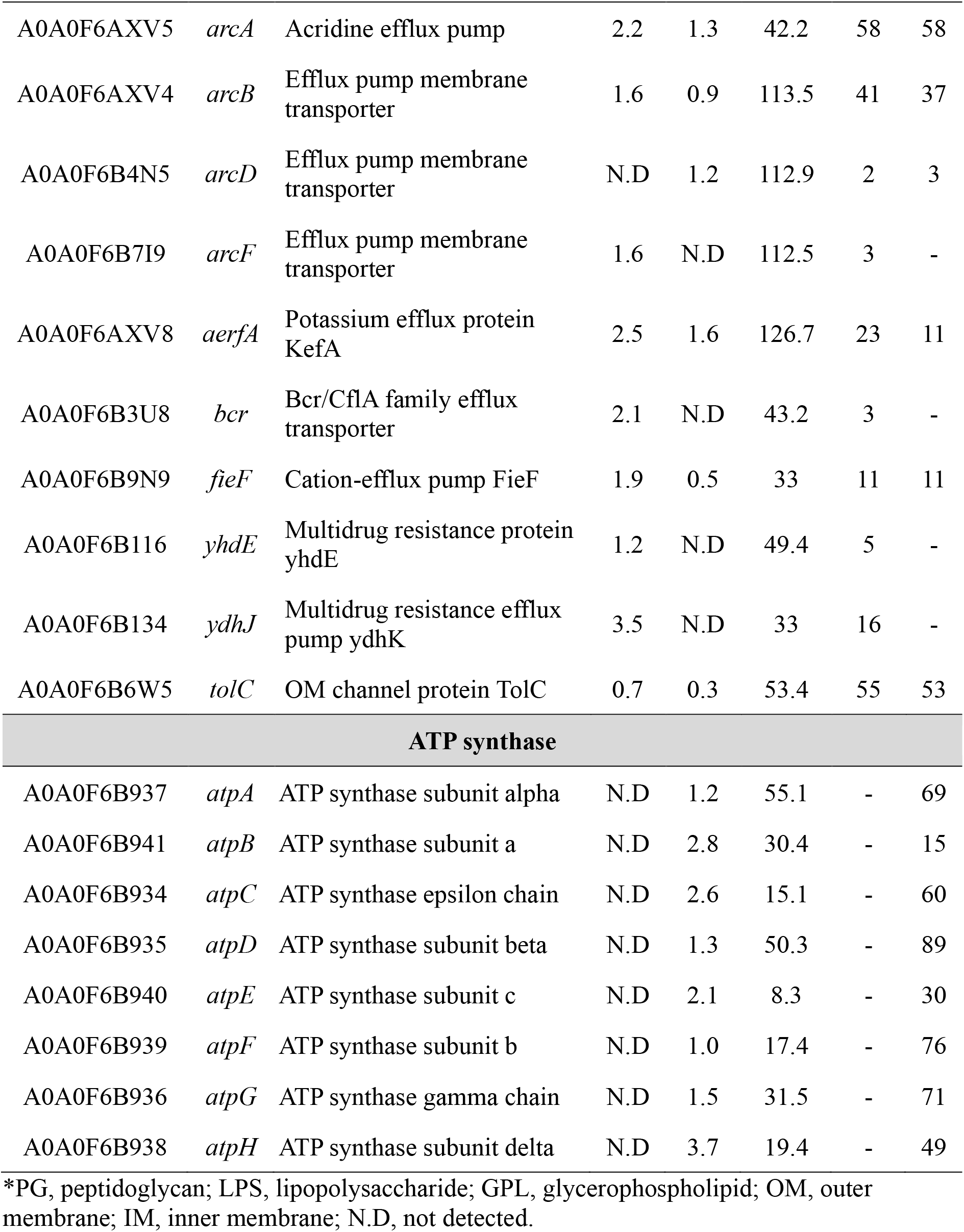
Membrane proteomic identification of envelope-localized proteins in *S.* Typhimurium 14028s under control and LEX stress conditions. WT strain culture were harvested at 6 h and fractionated into OM and IM fractions for proteomic analyses.

**Table S7.**
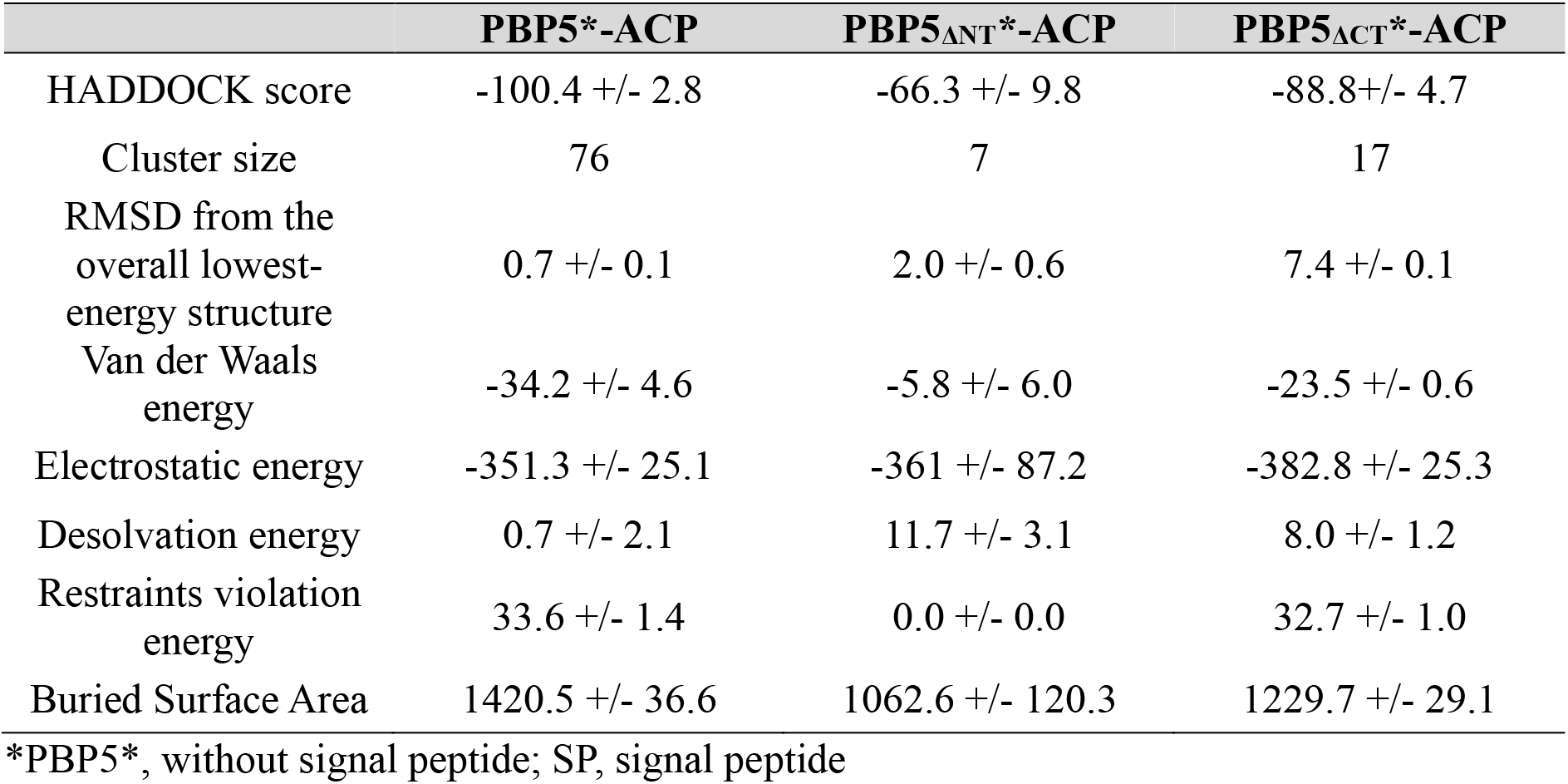
In silico HADDOCK docking scores of PBP5-ACP complexes.

**Table S8.** Bacterial strains and plasmids used in this study.

| Strains | Description* | Reference |
|---|---|---|
| ATCC 14028s | Wild type,<br>*S*. Typhimurium ATCC 14028s | This study |
| *dsbA*-HiBiT, *tatC*-3xFLAG | C-terminal of *dsbA* fused with 3xFLAG, C-terminal of *tatC* fused with 3xFLAG | This study |
| *ΔdacA* | *dacA* knockout mutant | This study |
| *ΔdacC* | *dacC* knockout mutant | This study |
| *ΔdacD* | *dacD* knockout mutant | This study |
| *ΔdacA*/pBAD33::*dacA* | Complementation strain; insertion of pBAD33-*dacA* with its promoter, into *ΔdacA* (Cm<sup>R</sup>) | This study |
| *ΔdacC*/pBAD33::*dacC* | Complementation strain; insertion of pBAD33-*dacC* with its promoter, into *ΔdacC* (Cm<sup>R</sup>) | This study |
| *ΔdacD*/pBAD33::*dacD* | Complementation strain; insertion of pBAD33-*dacD* with its promoter, into *ΔdacD* (Cm<sup>R</sup>) | This study |
| *ΔdacA*/pBAD33::*dacA<sub>OE</sub>* | Overexpression strain; insertion of pBAD33-*dacA*, into Δ*dacA* (Cm<sup>R</sup>) | This study |
| *ΔdacA*<sub>K242A</sub> | Point mutation of catalytic active site | This study |
| *ΔdacA*<sub>NT</sub> | Deletion of PBP5 N-terminal domain | This study |
| *ΔdacA*<sub>CT</sub> | Deletion of PBP5 C-terminal domain | This study |
| *dacA*-3xFLAG | C-terminal of *dacA* fused with 3xFLAG | This study |
| *dacC*-3xFLAG | C-terminal of *dacC* fused with 3xFLAG | This study |
| *dacD*-3xFLAG | C-terminal of *dacD* fused with 3xFLAG | This study |
| *mreA*-3xFLAG | C-terminal of *mreA* fused with 3xFLAG | This study |
| *ftsI*-3xFLAG | C-terminal of *ftsI* fused with 3xFLAG | This study |
| *dacA*-msfGFP | C-terminal of *dacA* fused with msfGFP | This study |
| WT/pSEVA234::SP<sub>*dsbA*</sub>-sfGFP | *P<sub>tac</sub>*-SP<sub>*dsbA*</sub>-sfGFP (Km<sup>R</sup>) | This study |
| *ΔdacA*/pSEVA234::SP<sub>*dsbA*</sub>-sfGFP | *P<sub>tac</sub>*-SP<sub>*dsbA*</sub>-sfGFP (Km<sup>R</sup>) | This study |
| *ΔdacA*<sub>NT</sub>/pSEVA234::SP<sub>*dsbA*</sub>-sfGFP | *P<sub>tac</sub>*-SP<sub>*dsbA*</sub>-sfGFP (Km<sup>R</sup>) | This study |
| *ΔdacA*<sub>CT</sub>/pSEVA234::SP<sub>*dsbA*</sub>-sfGFP | *P<sub>tac</sub>*-SP<sub>*dsbA*</sub>-sfGFP (Km<sup>R</sup>) | This study |
| *dacA*<sub>K242A</sub>/pSEVA234::SP<sub>*dsbA*</sub>-sfGFP | *P<sub>tac</sub>*-SP<sub>*dsbA*</sub>-sfGFP (Km<sup>R</sup>) | This study |
| *ftsI*<sup>down</sup> | *attTn7*::dCas9/pSGAb::*ftsI*<sub>spacer</sub> (Gm<sup>R</sup>, Km<sup>R</sup>) | This study |
| WT/pLumi-LgBiT | pLumi::*P<sub>rpo70</sub>*-LgBiT (AMP<sup>R</sup>) | This study |
| WT/pPBAD33::LgBiT | pBAD33::*P<sub>dsbA</sub>*-SP<sub>*dsbA*</sub>-LgBiT (CAP<sup>R</sup>) | This study |
| *acpP*-HiBiT/pLumi-LgBiT | C-terminal of *acpP* fused with HiBiT, pLumi::*P<sub>rpo70</sub>*-LgBiT (AMP<sup>R</sup>) | This study |
| *acpP*-HiBiT/pPBAD33::LgBiT | C-terminal of *acpP* fused with HiBiT, pBAD33::*P<sub>dsbA</sub>*-SP<sub>*dsbA*</sub>-LgBiT (CAP<sup>R</sup>) | This study |
| *dsbA*-HiBiT/pLumi-LgBiT | C-terminal of *dsbA* fused with HiBiT, pLumi::*P<sub>rpo70</sub>*-LgBiT (AMP<sup>R</sup>) | This study |
| *dsbA*-HiBiT/pPBAD33::LgBiT | C-terminal of *dsbA* fused with HiBiT, pBAD33::*P<sub>dsbA</sub>*-SP<sub>*dsbA*</sub>-LgBiT (CAP<sup>R</sup>) | This study |
| WT, *acpP*-HiBiT/pPBAD33::LgBiT | C-terminal of *acpP* fused with HiBiT, pBAD33::*P<sub>dsbA</sub>*-SP<sub>*dsbA*</sub>-LgBiT (CAP<sup>R</sup>) | This study |
| *ΔdacA, acpP*-HiBiT/pPBAD33::LgBiT | C-terminal of *acpP* fused with HiBiT, pBAD33::*P<sub>dsbA</sub>*-SP<sub>*dsbA*</sub>-LgBiT (CAP<sup>R</sup>) | This study |
| *ΔdacA*<sub>NT</sub>, *acpP*-HiBiT/pPBAD33::LgBiT | C-terminal of *acpP* fused with HiBiT, pBAD33::*P<sub>dsbA</sub>*-SP<sub>*dsbA*</sub>-LgBiT (CAP<sup>R</sup>) | This study |
| *ΔdacA*<sub>CT</sub>, *acpP*-HiBiT/pPBAD33::LgBiT | C-terminal of *acpP* fused with HiBiT, pBAD33::*P<sub>dsbA</sub>*-SP<sub>*dsbA*</sub>-LgBiT (CAP<sup>R</sup>) | This study |
| *dacA*<sub>K242A</sub>, *acpP*-HiBiT/pPBAD33::LgBiT | C-terminal of *acpP* fused with HiBiT, pBAD33::*P<sub>dsbA</sub>*-SP<sub>*dsbA*</sub>-LgBiT (CAP<sup>R</sup>) | This study |
| *acpP*<sup>down</sup> | *attTn7*::dCas9/pSGAb::*acpP*<sub>spacer</sub> (Gm<sup>R</sup>, Km<sup>R</sup>) | This study |
| *acpP*-3xFLAG | C-terminal of *ompC* fused with 3xFLAG | This study |
| WT, *ompC*-3xFLAG | C-terminal of *ompC* fused with 3xFLAG | This study |
| *ΔdacA*<sub>NT</sub>, *ompC*-3xFLAG | C-terminal of *ompC* fused with 3xFLAG | This study |
| *ΔdacA*<sub>CT</sub>, *ompC*-3xFLAG | C-terminal of *ompC* fused with 3xFLAG | This study |
| *ΔdacA*<sub>K242A</sub>, *ompC*-3xFLAG | C-terminal of *ompC* fused with 3xFLAG | This study |
| *acpP*<sup>down</sup>, *ompC*-3xFLAG | *attTn7*::dCas9/pSGAb::*acpP*<sub>spacer</sub> (Gm<sup>R</sup>, Km<sup>R</sup>), C-terminal of *ompC* fused with 3xFLAG | This study |
| *ΔcpxR* | *cpxR* knockout mutant | This study |
| *ΔompR* | *ompR* knockout mutant | This study |
| *ΔrpoE* | *rpoE* knockout mutant | This study |
| *ΔrcsB* | *rcsB* knockout mutant | This study |
| *ΔphoP* | *phoP* knockout mutant | This study |
| **Plasmid** | **Description** | **Reference** |
| pKD46 | Amp<sup>R</sup> | This study |
| pKD3 | Amp<sup>R</sup>, Cm<sup>R</sup> | This study |
| pKD4 | Amp<sup>R</sup>, Km<sup>R</sup> | This study |
| pCP20 | 3xFLP<sup>+</sup>, Amp<sup>R</sup>, Cm<sup>R</sup> | This study |
| pSUB11 | 3xFLAG sequence, Amp<sup>R</sup>, Km<sup>R</sup> | This study |
| pTNS3 | Transposase, Amp<sup>R</sup> | This study |
| pRK2013 | *trfA*, *traJ*, Km<sup>R</sup> | This study |
| pYDE009 | dCas9, Amp<sup>R</sup>, Gm<sup>R</sup> | This study |
| pSGAb | gRNA (spacer), Km<sup>R</sup> | This study |
| pKIKO | Knock-in/out, Amp<sup>R</sup> | This study |
| pKIKO::msfGFP | msfGFP, Km<sup>R</sup> | This study |
| pKIKO::sfGFP | sfGFP, Km<sup>R</sup> | This study |
| pKIKO::HiBiT | HiBiT sequence, Km<sup>R</sup> | This study |
| pLumi::LgBiT (cytosolic LgBiT) | LgBiT, Amp<sup>R</sup> | This study |
| pBAD33::*P<sub>dsbA</sub>*-SP<sub>*dsbA*</sub>_LgBiT (Periplasmic LgBit) | LgBiT, Cm<sup>R</sup> | This study |

\*All *S*. Typhimurium strains listed are isogenic and derivatives of parental strain 14028s.

**Table S9.**
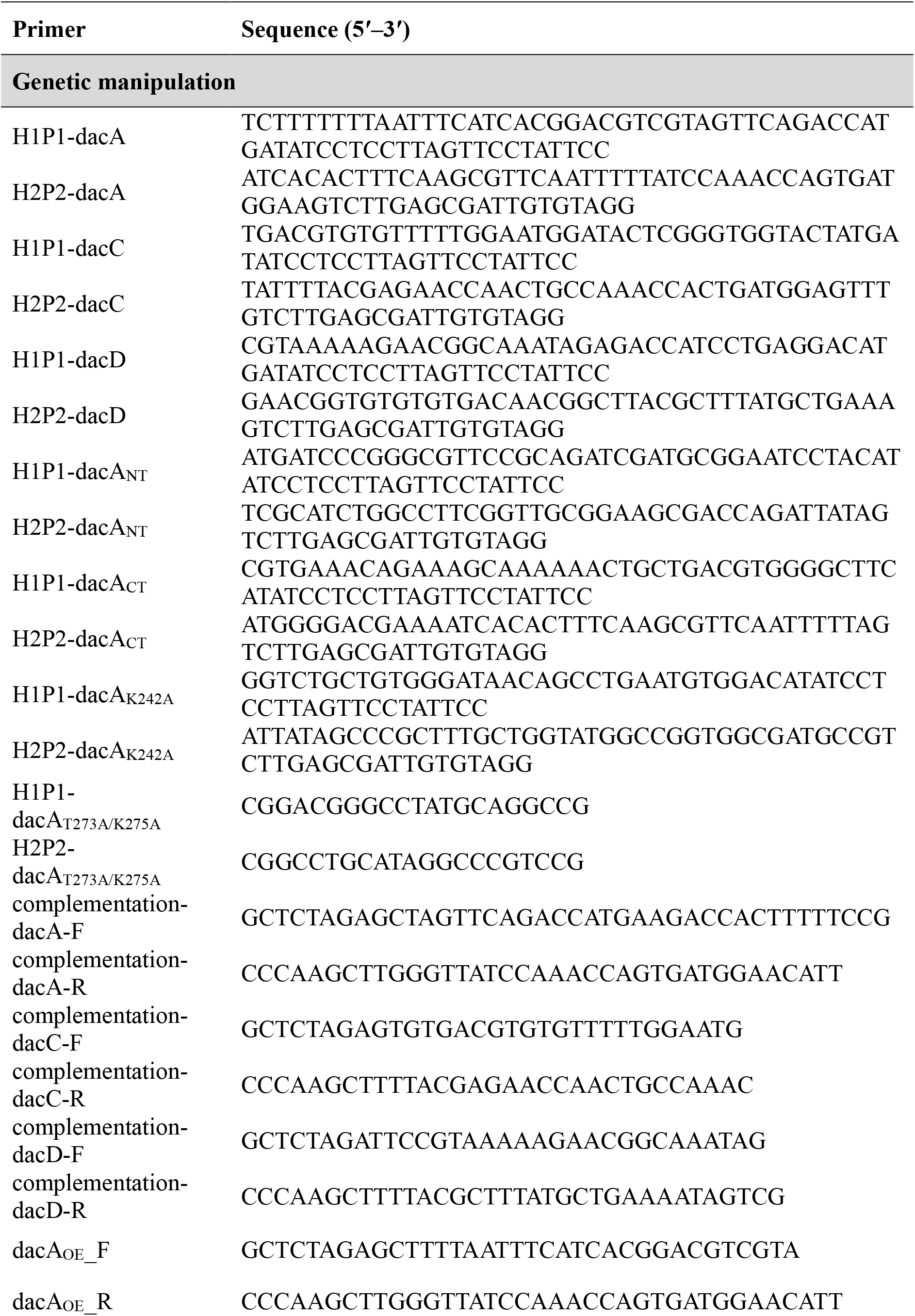
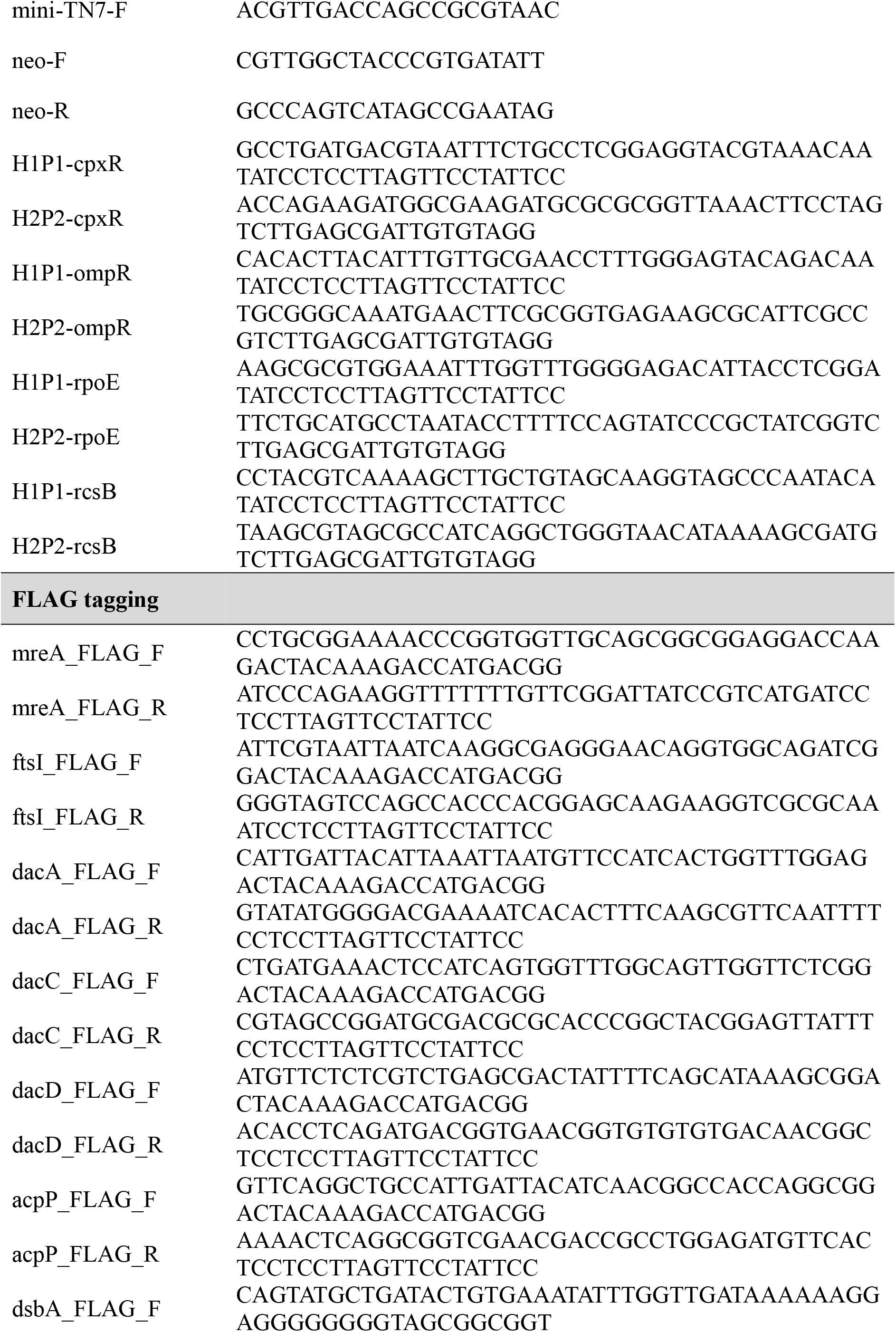
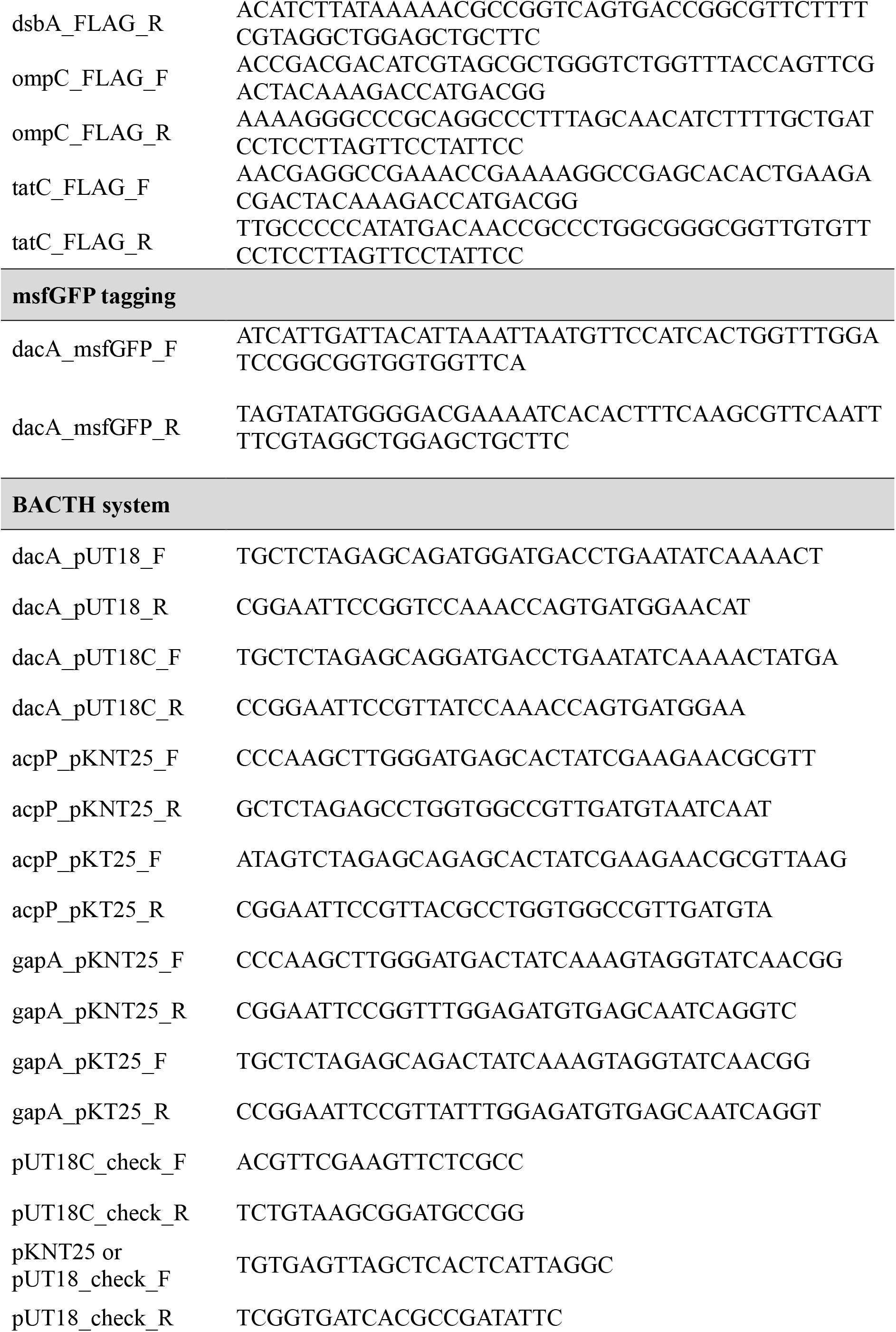
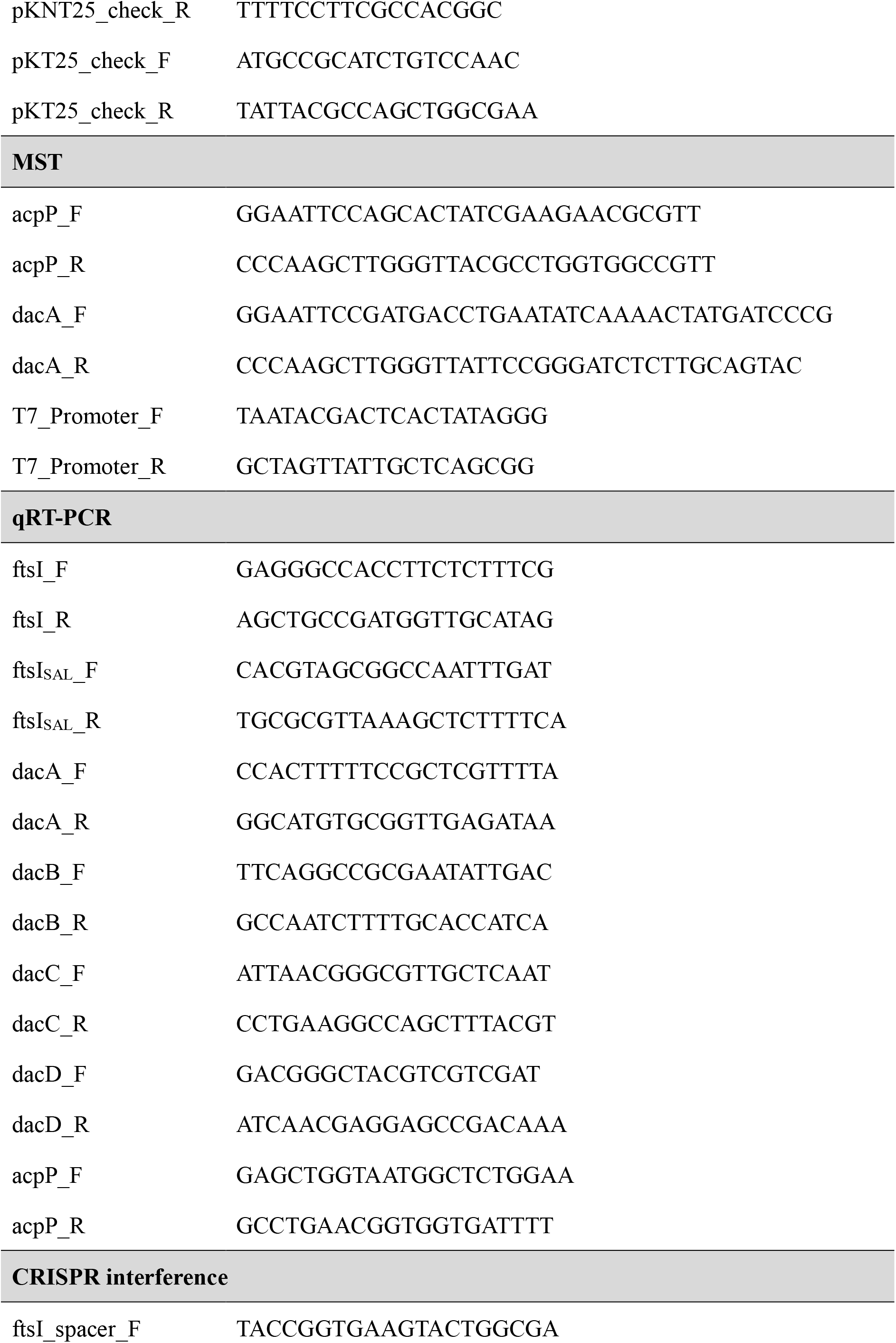
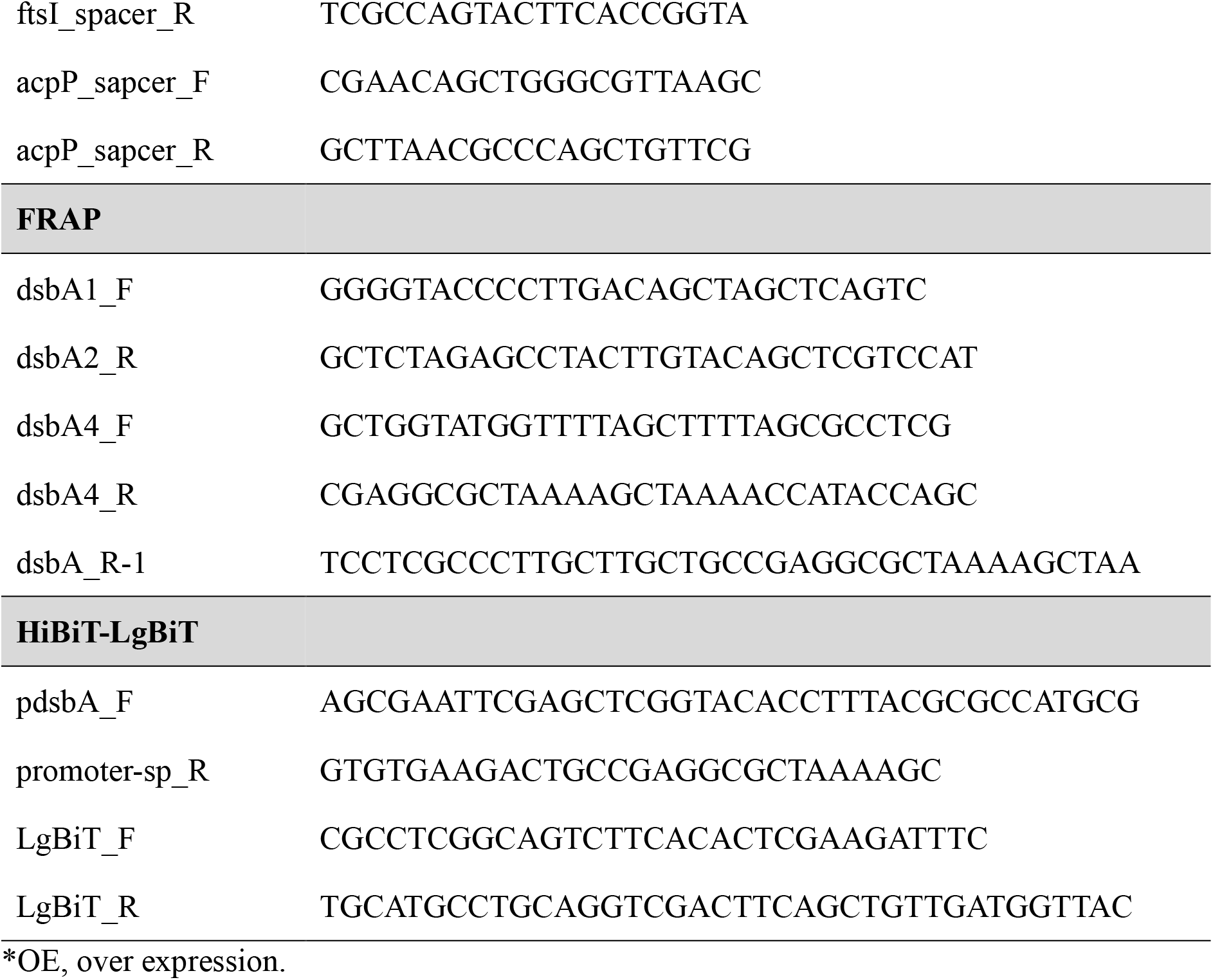
Oligonucleotides used in this study.

**Figure S1.**
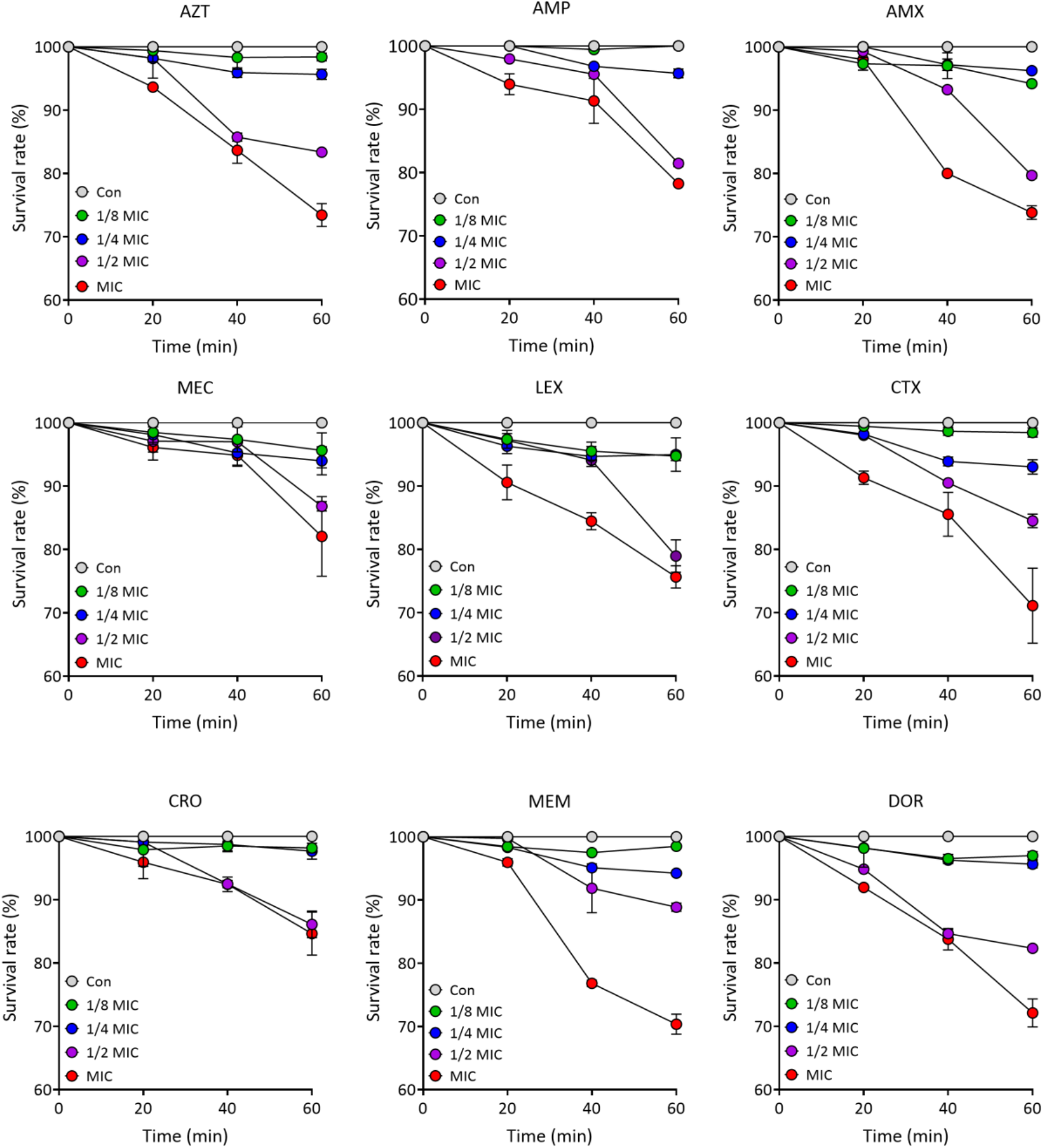
Short-term killing curves of β-lactam-treated *S.* Typhimurium cells. Exponentially growing *S.* Typhimurium cells (for 3 h) were adjusted to an initial density of 1 × 10^6^ cells/mL in LB broth and exposed to β-lactam antibiotics at concentrations ranging from 1 × MIC to 1/8 × MIC under static conditions at 37°C. Cell numbers were measured by CFU assay every 20 min for 60 min, and viability loss was calculated relative to the initial cell number at 0 min. *Aztreonam, AZT; Ampicillin, AMP; Amoxicillin, AMX; Mecillinam, MEC; Cephalexin, LEX; Cefotaxime, CTX; Ceftriaxone, CRO; Meropenem, MEM; Doripenem, DOR.

**Figure S2.**
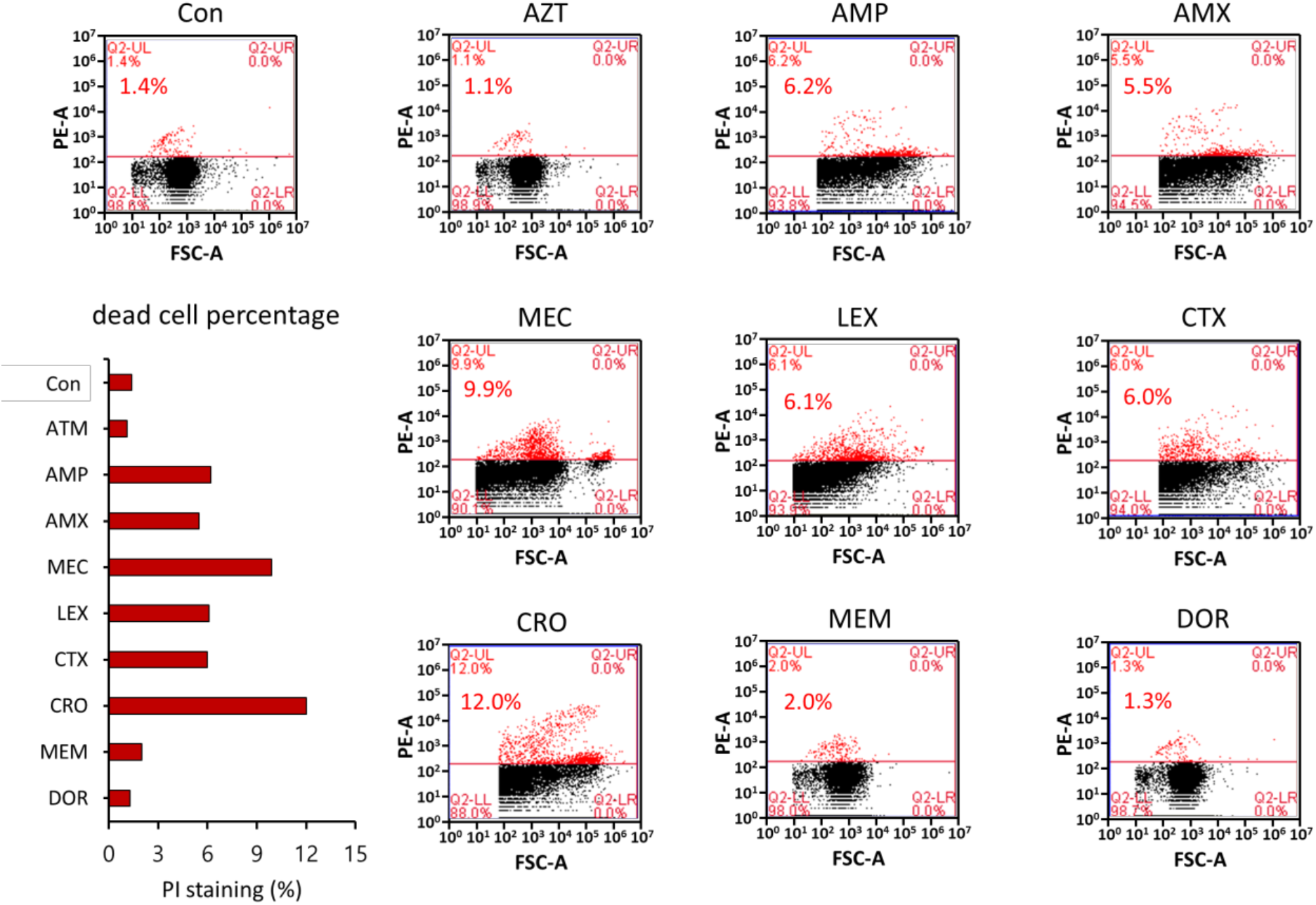
Measurement of the dead cells using FACS. *S*. Typhimurium cells were exposed to β-lactam antibiotics at 1/4 × MIC in LB broth under static conditions at 37°C for 6 h. After 6 hours, cells were harvested, washed three times with PBS, and stained with propidium iodide (PI) in the dark for 20 min. After three additional PBS washes, PI-stained dead cells were quantified by FACS analyses, with 50,000 cells analyzed per sample. *Aztreonam, AZT; Ampicillin, AMP; Amoxicillin, AMX; Mecillinam, MEC; Cephalexin, LEX; Cefotaxime, CTX; Ceftriaxone, CRO; Meropenem, MEM; Doripenem, DOR.

**Figure S3.**
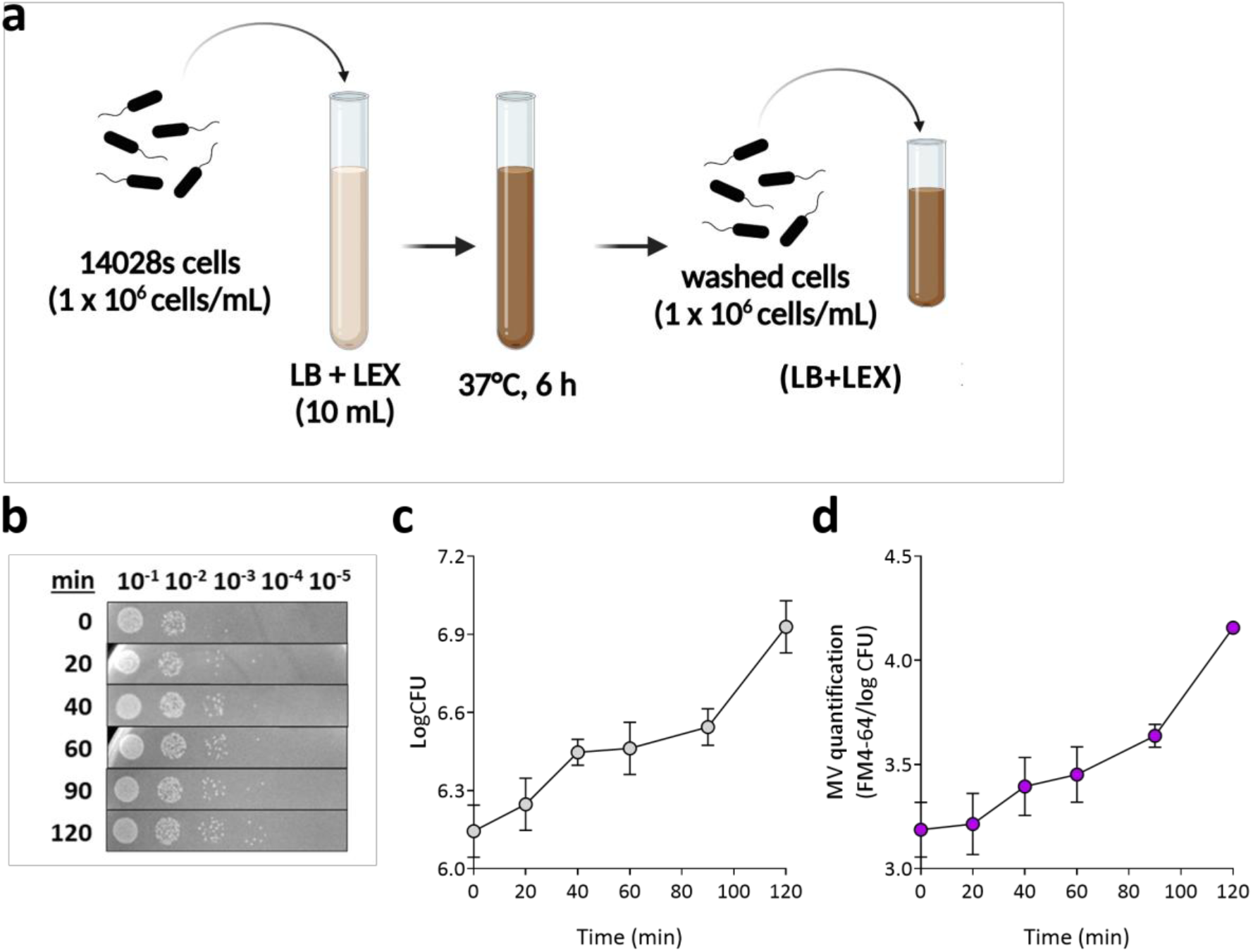
Conditioned medium re-inoculation assay. **a** Schematic of the conditioned medium re-inoculation assay. *S.* Typhimurium cells were exposed to LEX in LB broth under static conditions at 37°C for 6 h. Cells were then harvested, washed, and re-inoculated at 1 ×10^6^ cells/mL into cell-free conditioned medium prepared by filtration through a 0.2-µm membrane. **b–d** After re-inoculation, cell numbers and MV production were measured at the indicated time points by CFU assay and FM4-64 dye, respectively.

**Figure S4.**
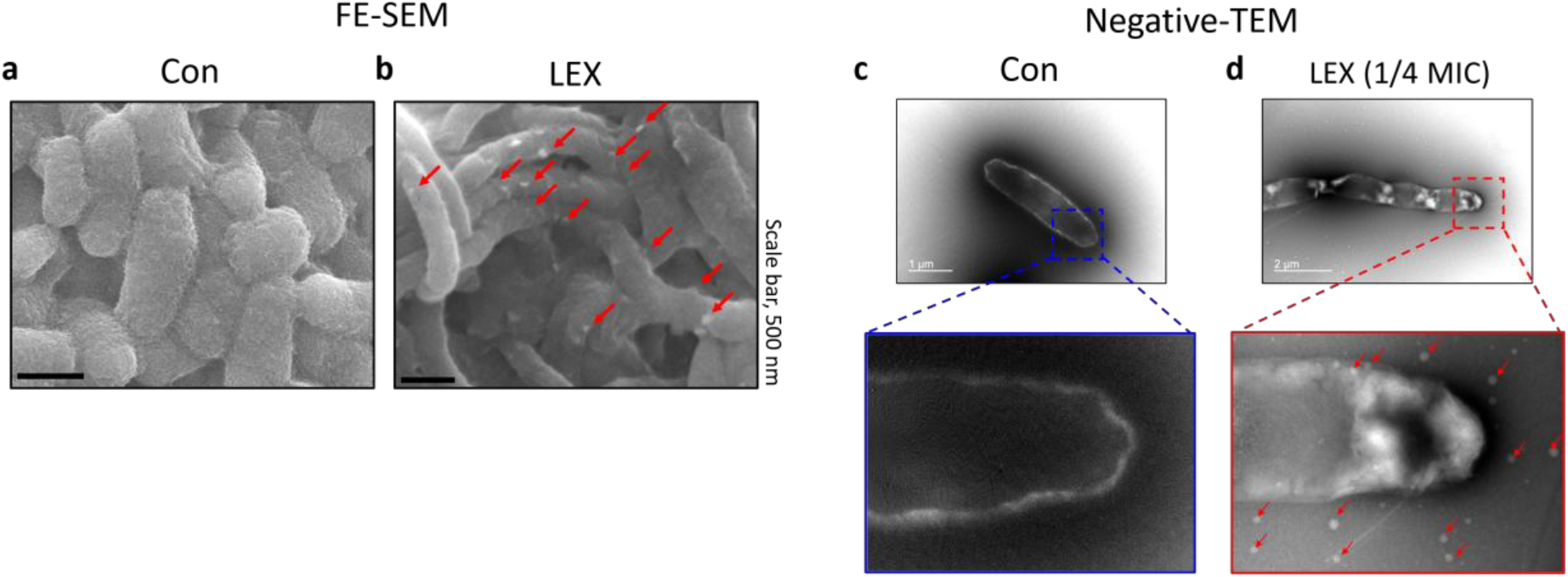
FE-SEM and negative-stain TEM observation of LEX-treated and untreated *S.* Typhimurium cells. *S.* Typhimurium cells were cultured in LB broth with or without under static conditions at 37°C for 6 h and analyzed by negative-stain transmission electron microscopy (TEM). FE-SEM and negative-stain TEM images show **a, c** control cells and **b, d** LEX-treated cells with OMV formation. Red arrows indicate OMVs.

**Figure S5.**
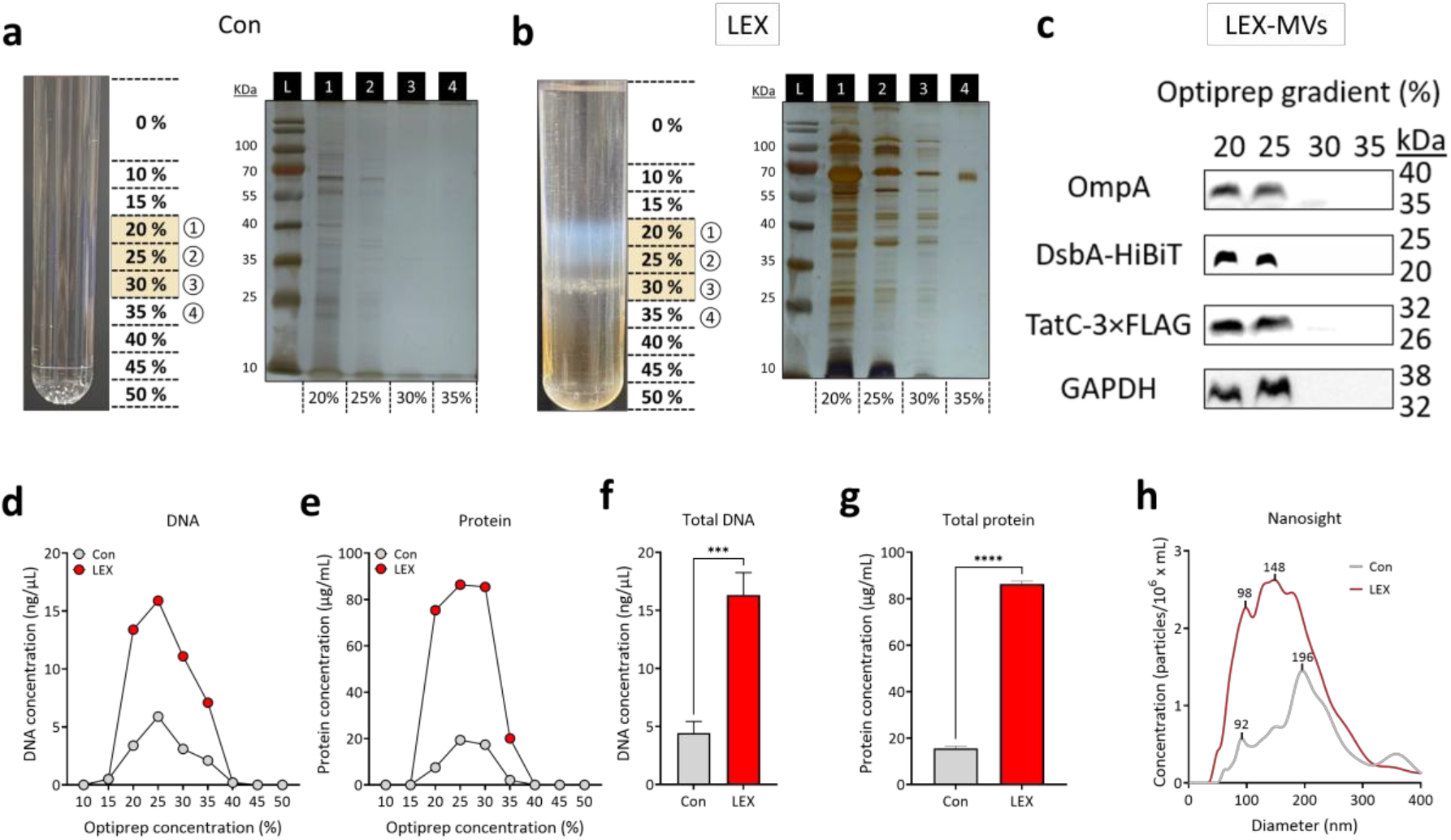
Purification and biomolecular analysis of MVs derived from LEX-treated and untreated *S.* Typhimurium cells. OMVs were purified from 2 L culture supernatants of untreated control and LEX-treated *S.* Typhimurium cells by OptiPrep density gradient centrifugation. **a, b** OptiPrep gradient images of MV fractions isolated from **a** control and **b** LEX-treated cultures. Fractions were recovered from each OptiPrep density range and analyzed by SDS-PAGE followed by silver staining. The white band in **b** indicates the OMV-enriched fraction recovered for further analysis. **c** Western blot analysis of MVs isolated from LEX-treated S. Typhimurium cells carrying chromosomal *dsbA*–HiBiT and *tatC*–3×FLAG fusions following OptiPrep density-gradient fractionation. Fractions corresponding to 20%, 25%, 30%, and 35% OptiPrep were analyzed for OmpA (OM marker), DsbA–HiBiT (periplasmic marker), TatC–3×FLAG (IM marker), and GAPDH (cytoplasmic marker). **d, e** DNA and protein contents of individual OptiPrep gradient fractions. **f, g** Total DNA and protein contents of purified OMV. **h** Particle size distribution of purified MVs was analyzed using NanoSight LM10. Statistical comparisons were performed using one-way ANOVA with Tukey’s multiple comparison test (***P < 0.001 and ****P < 0.001).

**Figure S6.**
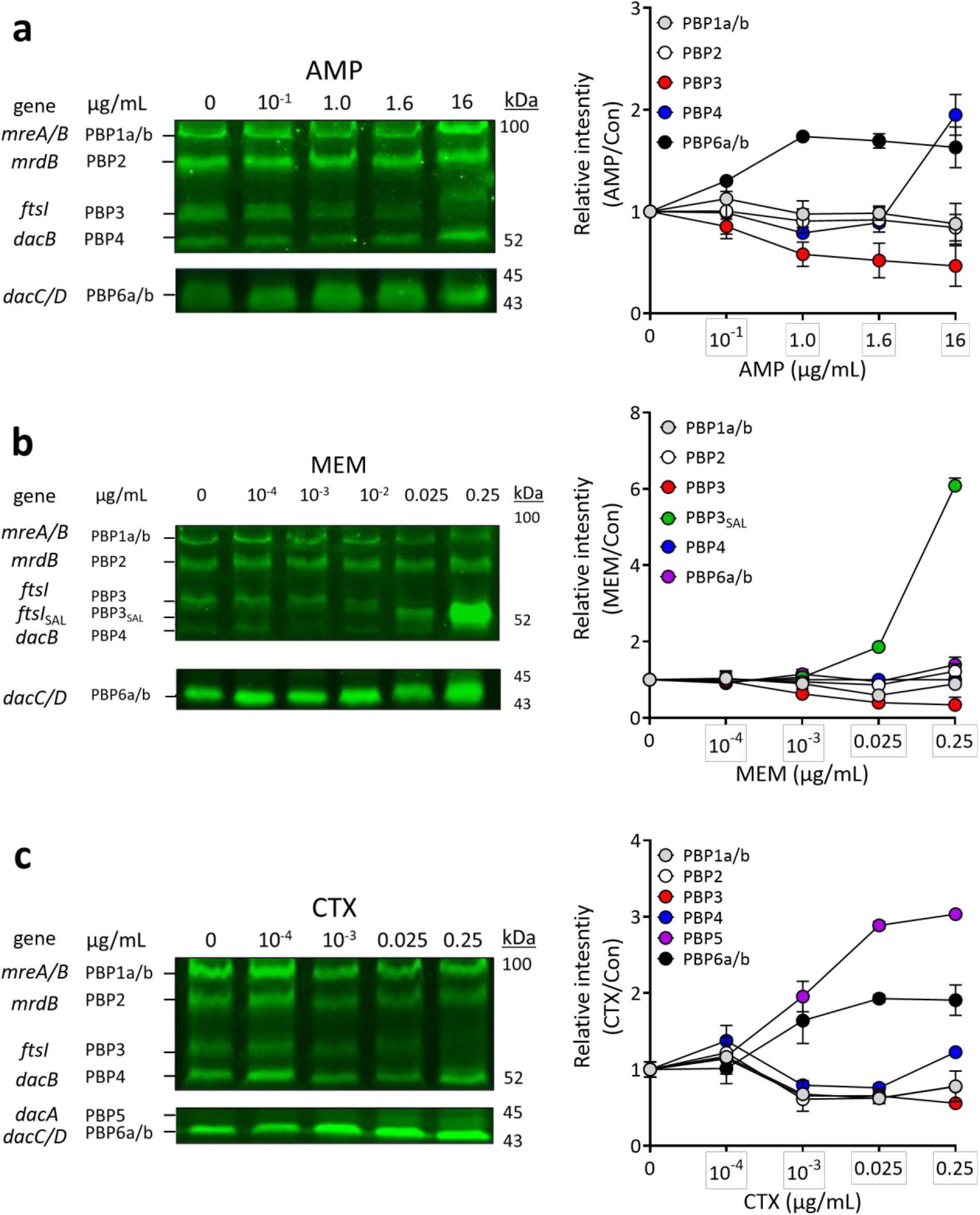
PBP profiling of β-lactam-treated *S.* Typhimurium cells using Bocillin-FL binding assay. *S.* Typhimurium cells were exposed to different concentrations of β-lactam antibiotics and subjected to Bocillin-FL binding assay to assess active PBP profiles. **a–c** SDS-PAGE gel images of Bocillin-FL–labeled PBPs and ImageJ-based quantification of Bocillin-FL signals in cells treated with **a** AMP, **b** MEM, and **c** CTX. Relative PBP levels were calculated by normalizing each signal intensity to the 0 µg/mL condition.

**Figure S7.**
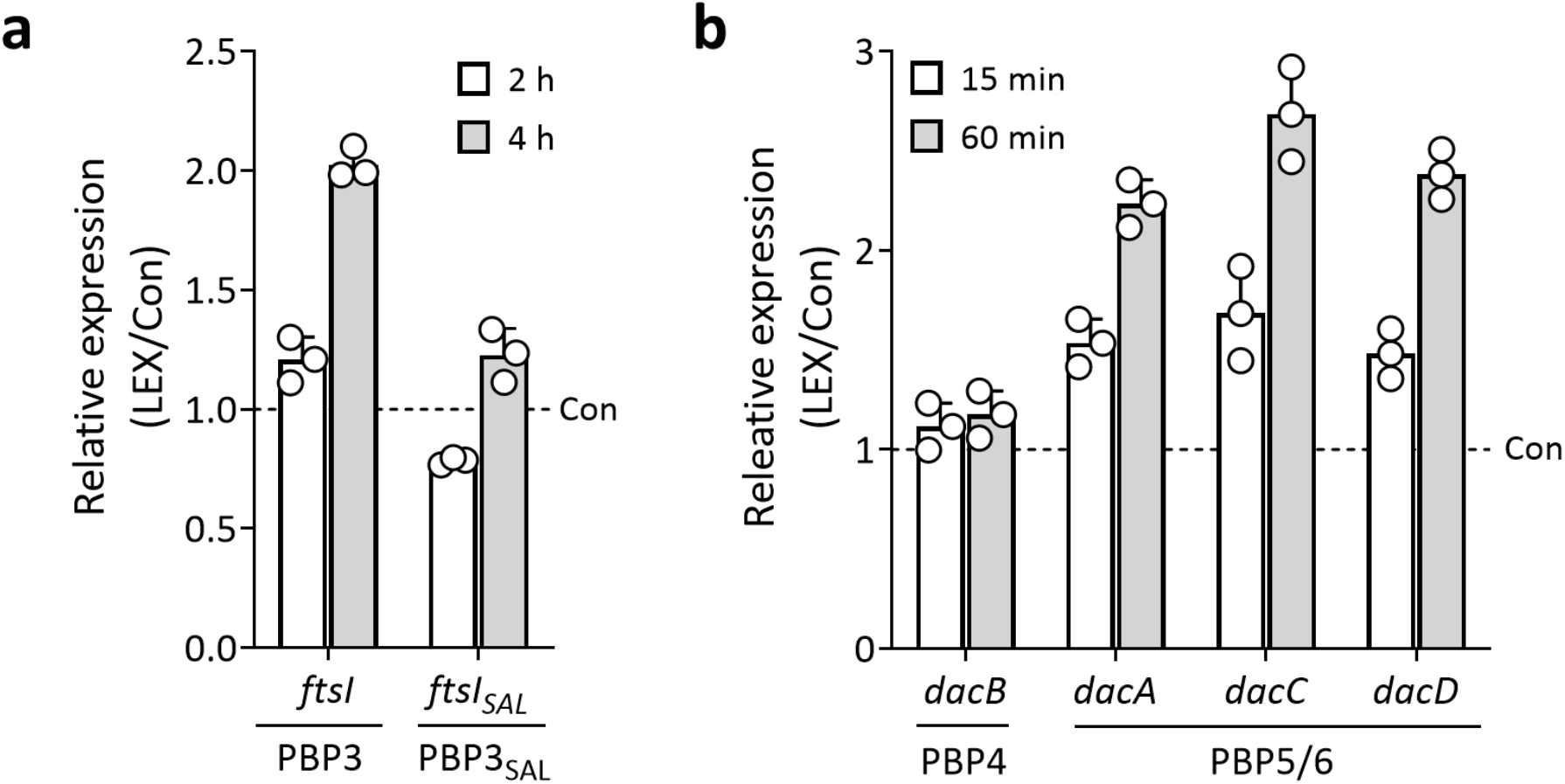
Relative expression of genes encoding PBPs in *S.* Typhimurium under LEX stress. Relative expression levels of genes encoding PBPs were measured by qRT-PCR in *S*. Typhimurium cells exposed to LEX. **a** Relative expression of *ftsI* and *ftsI_SAL_*, encoding PBP3 and PBP3_SAL_, respectively, was measured after 2 and 4 h of LEX exposure. **b** Relative expression of *dacB, dacA*, *dacC*, and *dacD*, encoding PBP4, PBP5, PBP6a, and PBP6b, respectively, was measured after 15 and 60 min of LEX exposure. Relative expression levels were normalized to 16S rRNA and calculated relative to the untreated control (Con) condition at each corresponding time point.

**Figure S8.**
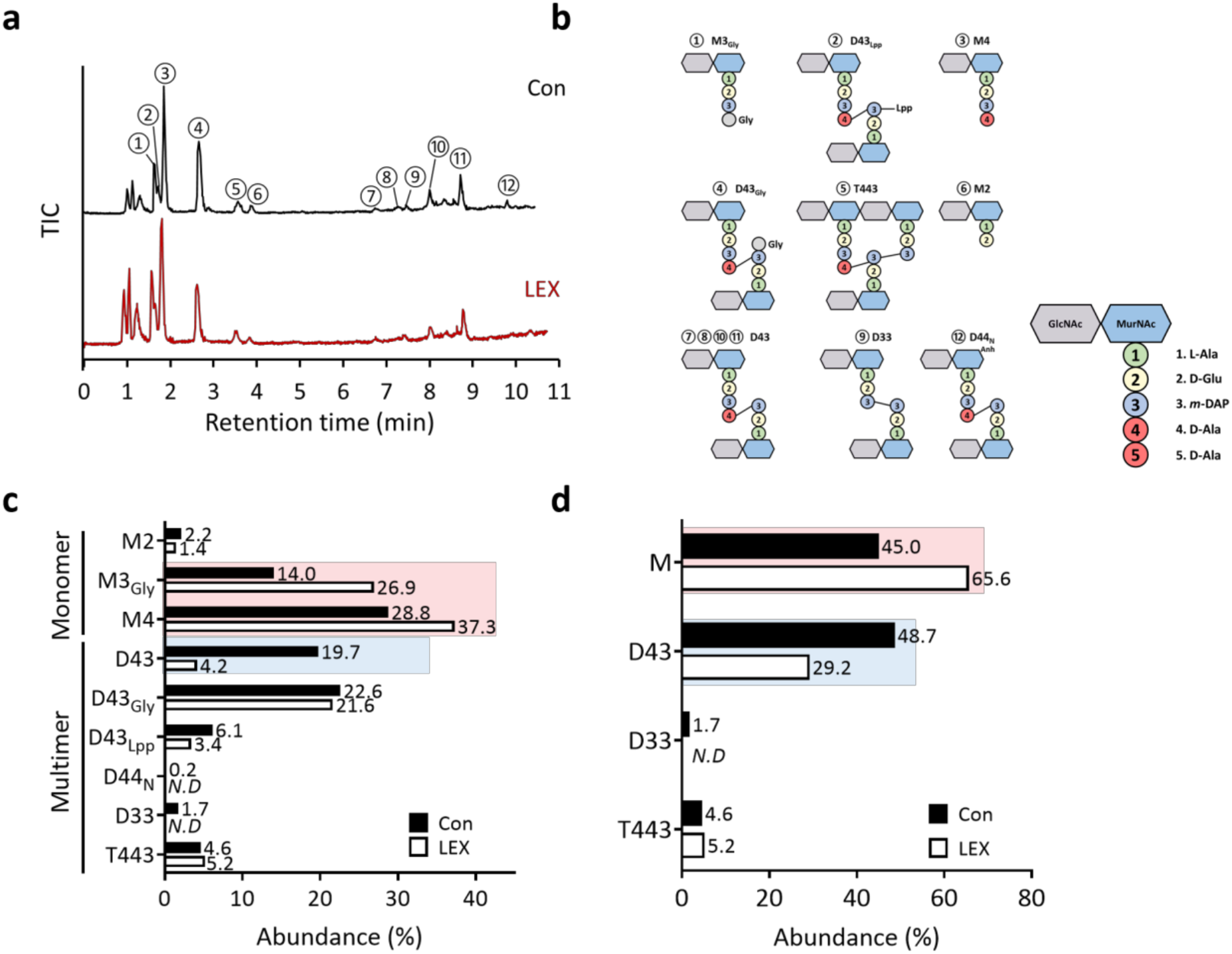
Muropeptide profiling of peptidoglycan from LEX-treated and untreated *S*. Typhimurium cells. Muropeptides were analyzed by LC-Q/TOF-MS after 6 h of culture in LB broth (1 L) with or without LEX. **a** Total ion chromatogram of muropeptides from control and LEX-treated cells. **b** Schematic structures of the detected muropeptide species. **c** Relative abundance of individual muropeptide species in control and LEX-treated cells. Red and blue background shading indicate muropeptide species increased and decreased under LEX stress, respectively. **d** Relative proportions of selected muropeptide classes, including monomeric muropeptides, D46, D33, and T443, calculated from the data shown in **c**. M, D, and T indicate monomeric, dimeric, and trimeric muropeptides, respectively. M2, monomeric disaccharide-dipeptide muropeptide; M3, monomeric disaccharide-tripeptide muropeptide; M3_Gly_, monomeric disaccharide-tripeptide muropeptide with glycine as the fourth stem residue; M4, monomeric disaccharide-tetrapeptide muropeptide; M5, monomeric disaccharide-pentapeptide muropeptide; D33, dimeric muropeptide composed of two tripeptide-containing disaccharide units; D43, dimeric muropeptide composed of one tetrapeptide-containing and one tripeptide-containing disaccharide units; D43_Lpp_, dimeric muropeptide composed of tetrapeptide- and tripeptide-containing disaccharide units covalently linked to Lpp; D43_Gly_, dimeric muropeptide composed of tetrapeptide- and tripeptide-containing disaccharide units, with glycine at the fourth stem position; D44_N_, dimeric muropeptide composed of two tetrapeptide-containing disaccharide units containing an anhydro-MurNAc residue; T443, trimeric muropeptide composed of two tetrapeptide-containing and one tripeptide-containing disaccharide units.*N. D, not detected.

**Figure S9.**
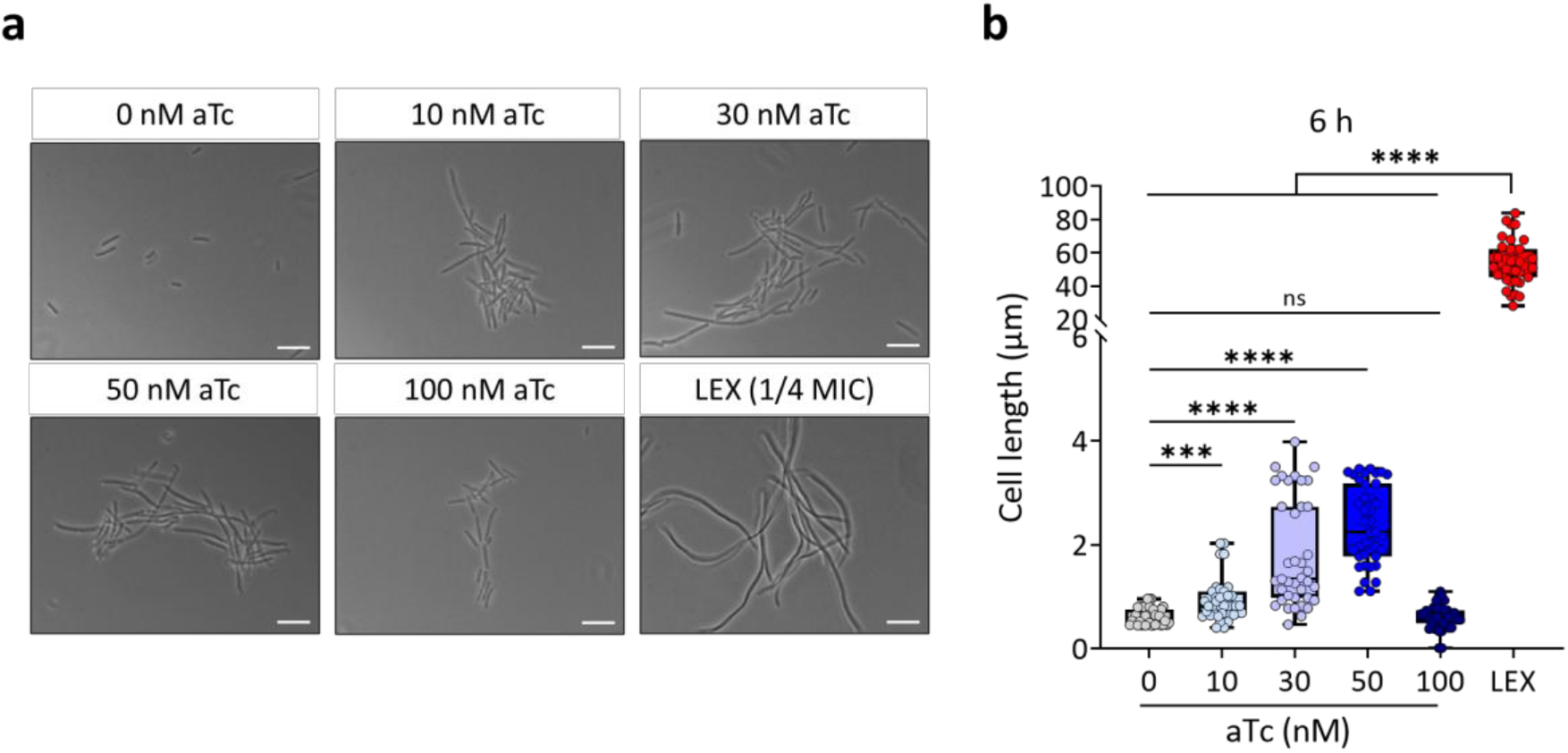
Microscopic analyses of cellular morphology in *ftsI*^down^ strain. The *ftsI^down^* strain, carrying chromosomally integrated *attTn7*::dCas9 and the pSGAb::*ftsI*_spacer_, was cultured in LB broth under static conditions at 37°C and treated with different concentrations of aTc to induce dCas9-dependent repression of *ftsI* gene, and cellular morphology was examined by microscopy. LEX-treated cells were included as a comparison. **a** Representative microscopy images of cells treated with 0, 10, 30, 50, or 100 nM aTc, and cells treated with LEX. Scale bars, 1 µm. **b** Cell length was measured using ZEN Blue edition software. Statistical comparisons were performed using one-way ANOVA with Tukey’s multiple comparison test (***P < 0.001 and ****P < 0.001; ns, not significant).

**Figure S10.**
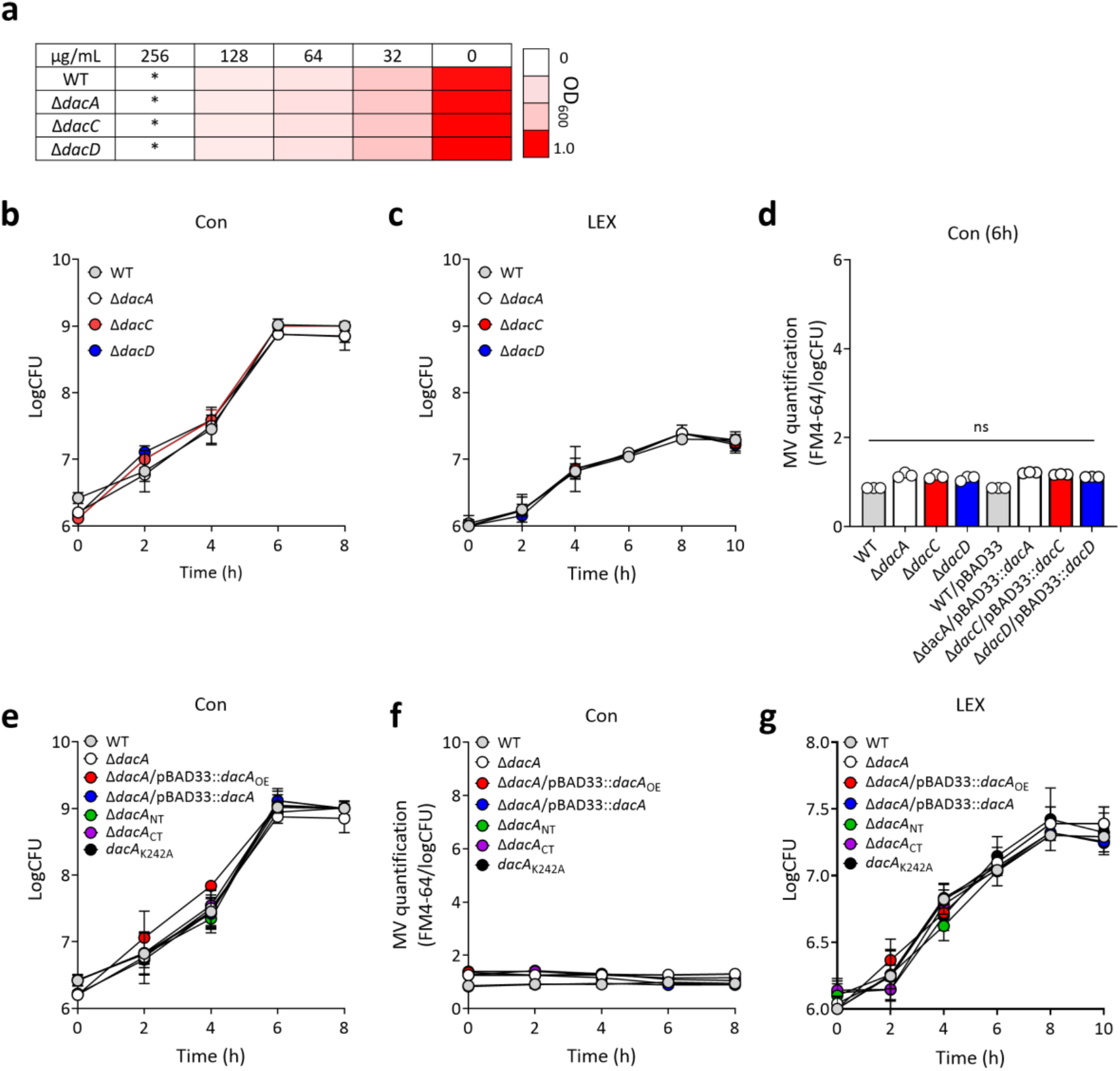
Growth and MV production of DD-carboxypeptidase single-knockout strains and *dacA* variants. WT, DD-carboxypeptidase single-knockout strains, and *dacA* variants were cultured in LB broth under static conditions at 37°C. **a** Evaluation of MIC level in WT, *AdacA, AdacC*, and *AdacD* single-knockout strains under LEX stress. **b, c** Growth curves of WT, *AdacA, AdacC*, and *AdacD* single-knockout strains under control and LEX stress conditions, respectively. **d** MV production of WT, *AdacA*, *AdacC*, and *AdacD* single-knockout strains under control conditions. **e, f** Growth curves and OMV production of WT and *dacA* variants under control conditions, respectively. **g** Growth curves of WT and *dacA* variants under LEX stress. Statistical comparisons were performed using one-way ANOVA with Tukey’s multiple comparison test (ns, not significant). *OE, overexpression; ΔNT, N-terminal domain deletion; ΔCT, C-terminal domain deletion.

**Figure S11.**
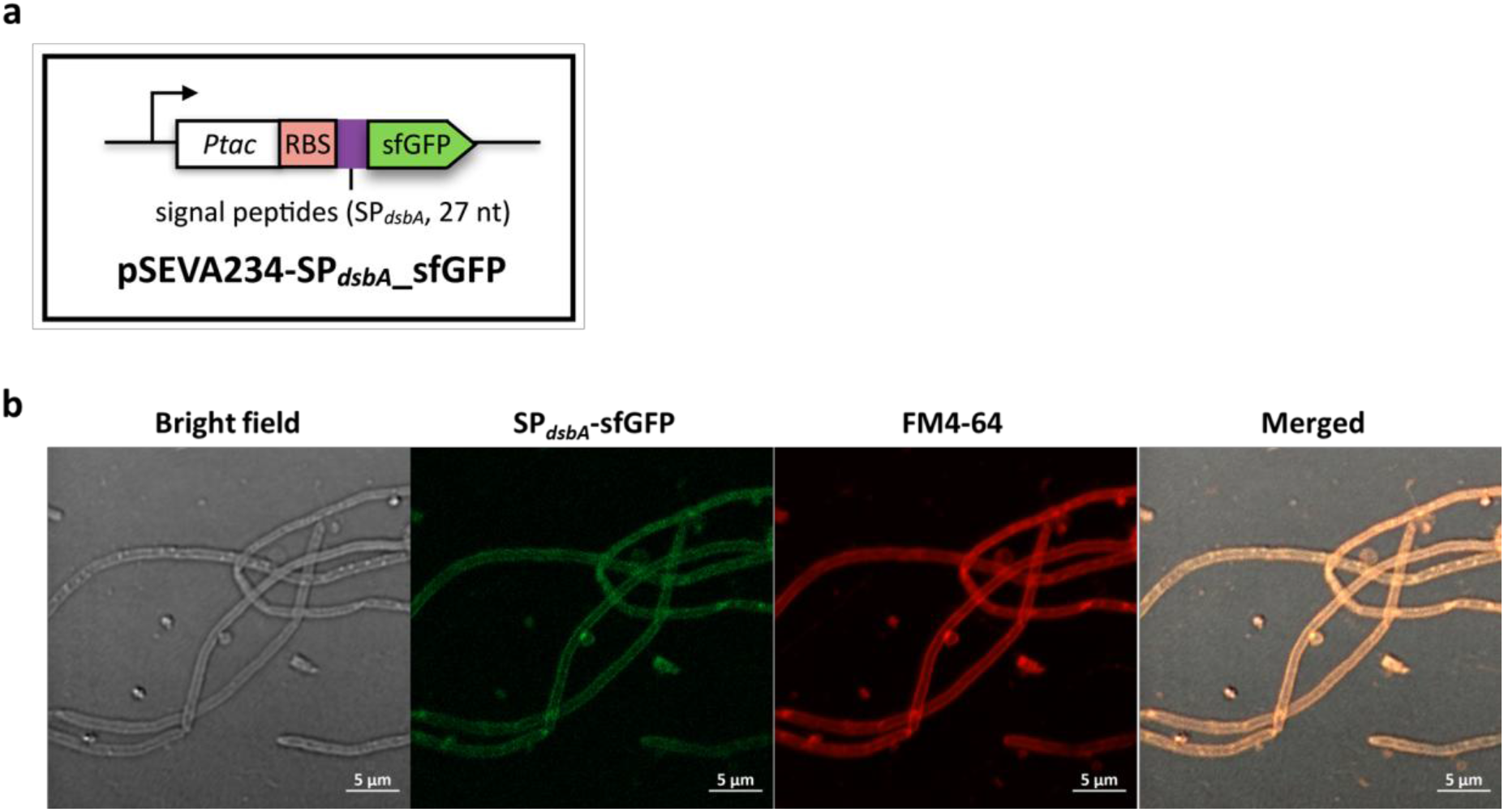
Construction of SP*_dsbA_*-sfGFP for FRAP assay to assess periplasmic crowding. **a** Schematic representation of the pSEVA234-SP*_dsbA_*–sfGFP construct. The signal peptide sequence of *dsbA* gene was fused to sfGFP to the cell envelope/periplasmic region. **b** Fluorescence microscopy images showing that sfGFP fused to the DsbA signal peptide localized to the periplasmic/cell envelope region. Cells were stained with FM4-64 to visualize OM. *Scale bars, 5 µm. SP, signal peptide.

**Figure S12.**
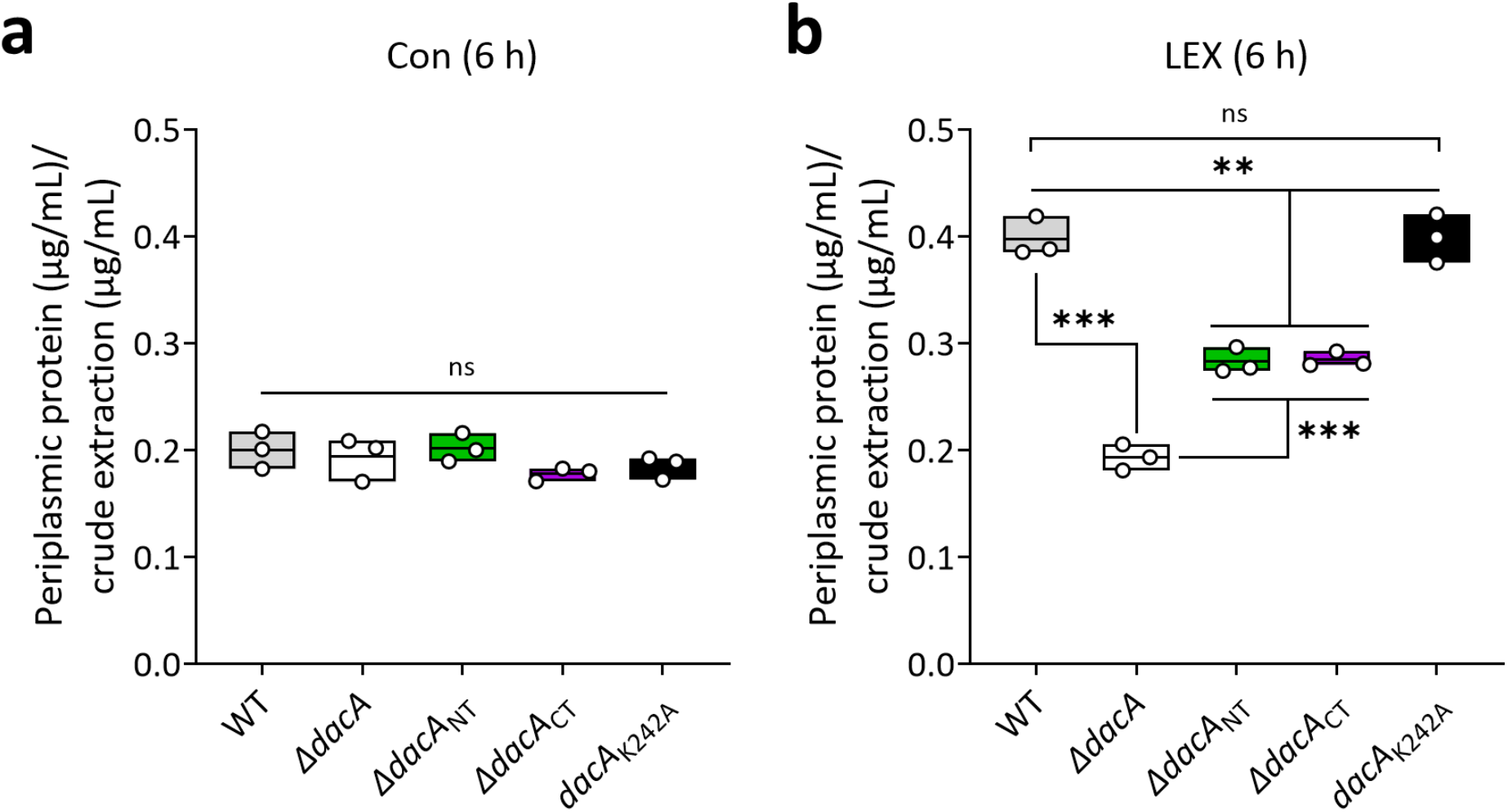
Quantification of periplasmic protein level in WT and *dacA* variants. WT and *dacA* variant cells were cultured in LB broth with or without LEX under static conditions at 37°C for 6 h. Protein concentrations in the periplasmic fractions were measured by Bradford assay and normalized to the corresponding crude extract protein levels. **a** Relative periplasmic protein levels under control conditions. **b** Relative periplasmic protein levels under LEX stress. Statistical comparisons were performed using one-way ANOVA with Tukey’s multiple comparison test (**P < 0.01 and ***P < 0.001; ns, not significant).

**Figure S13.**
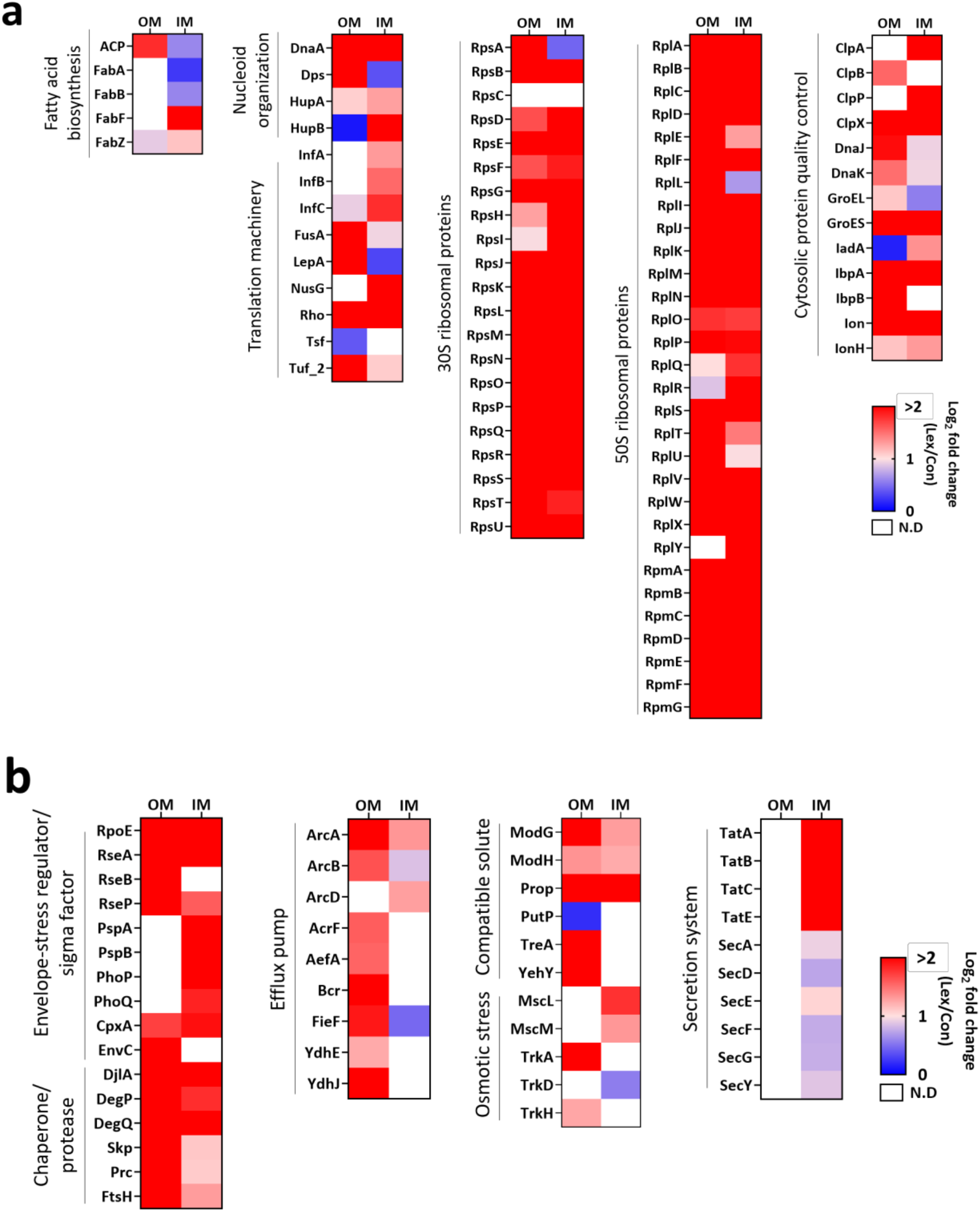
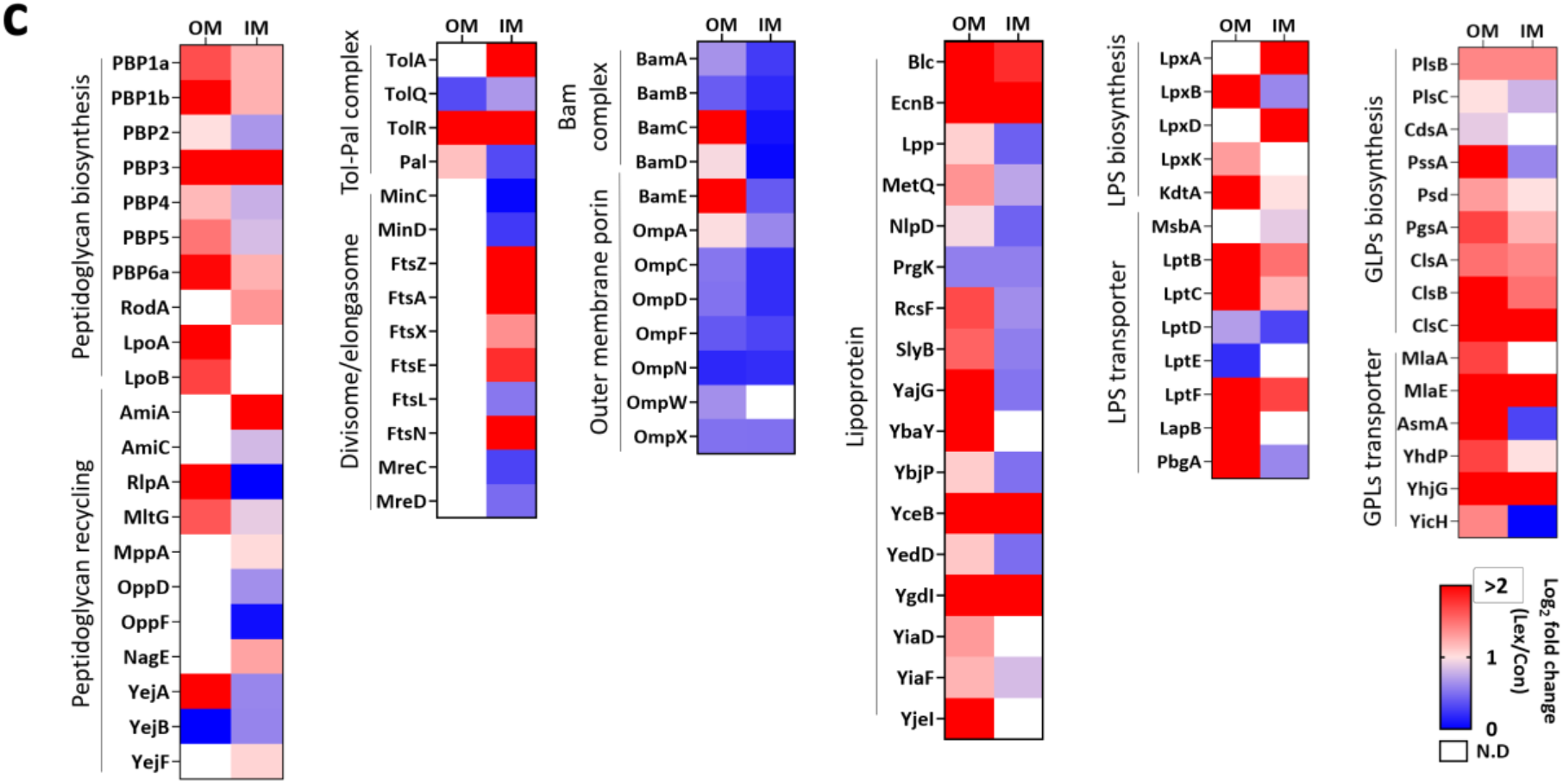
Membrane proteomic analyses of OM and IM fractions from LEX-treated and untreated *S.* Typhimurium cells. Proteins detected in OM and IM fractions were grouped by functional category and compared between control and LEX-treated cells. **a** Cytosolic proteins. **b** Stress response and protein homeostasis-related proteins. **c** Cell envelope biogenesis-related proteins. The heatmap represents log_2_ fold changes in protein abundance under LEX stress relative to the untreated control. Red and blue indicate increased and decreased abundance under LEX stress, respectively.

**Figure S14.**
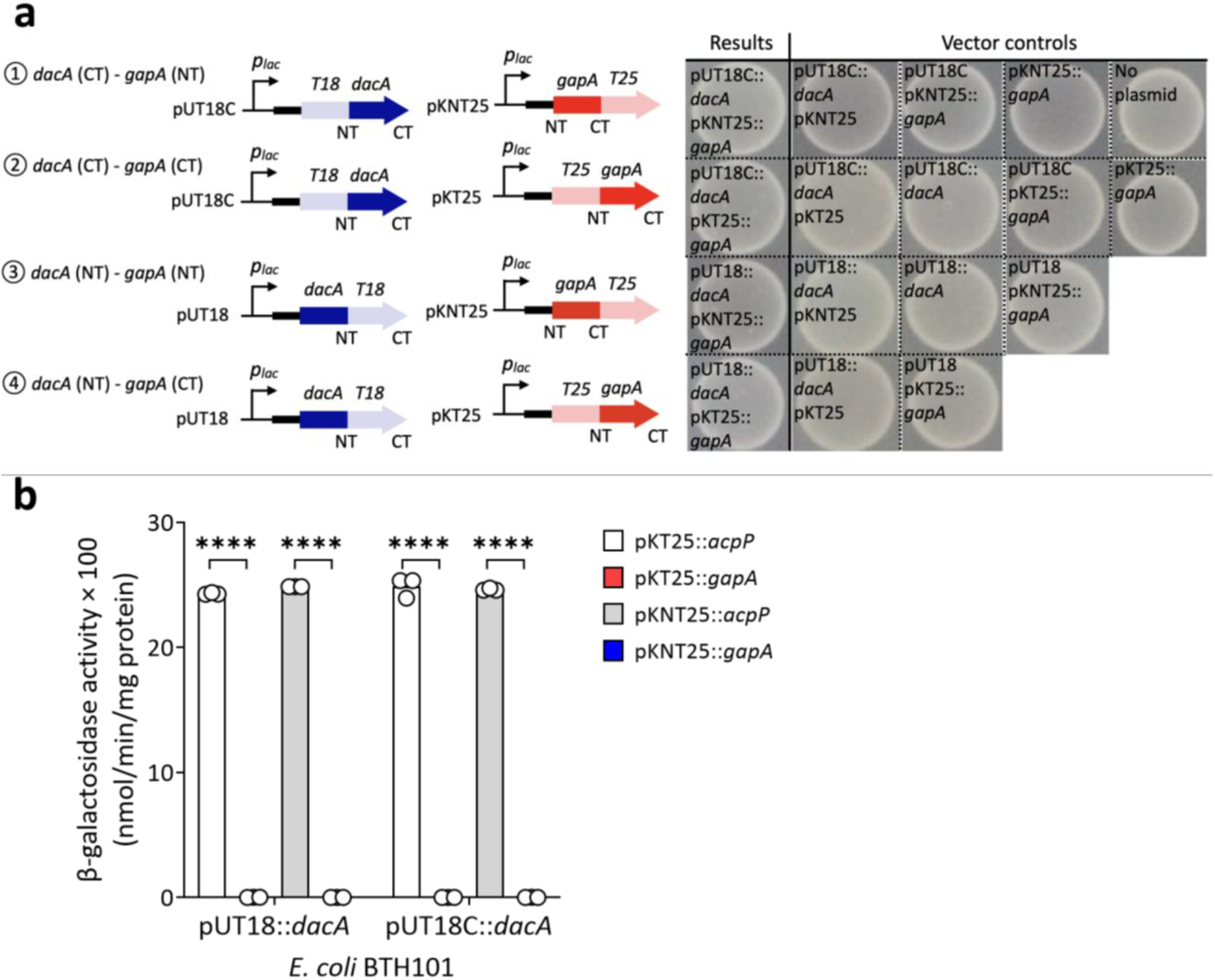
Specificity of the PBP5–ACP interaction assessed by BACTH assays. **a** Schematic representation of the four fusion configurations and representative BACTH assay results for PBP5* (*dacA*) and GAPDH (*gapA*), including the corresponding vector controls. **b** β-Galactosidase activities of *E. coli* BTH101 cells expressing PBP5* (*dacA*) fused to T18 or T18C in combination with ACP (*acpP*) or GAPDH (*gapA*) fused to T25 or NT25. ****P < 0.0001.

**Figure S15.**
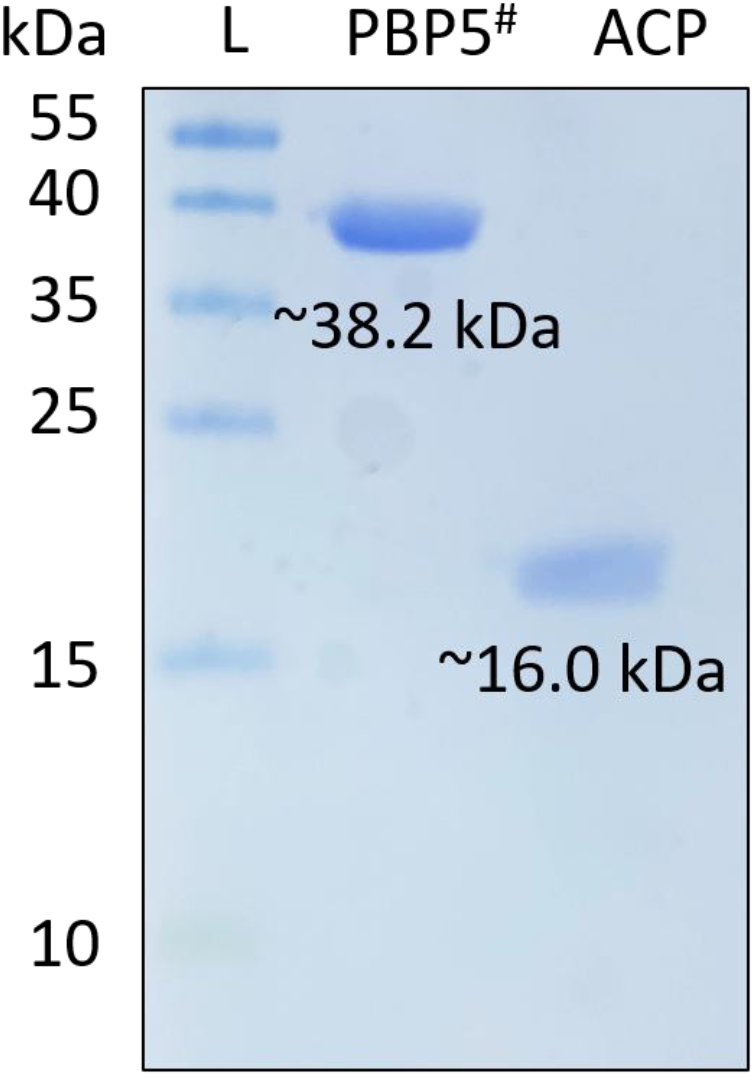
Purification of PBP5 and ACP for MST analyses. PBP5^#^ and ACP proteins expressed in *E. coli* BL21 (DE3) were purified and loaded on 12% SDS-PAGE. PBP5^#^ was expressed and purified as a truncated form lacking both the signal peptide and membrane-anchoring sequence. The purified PBP5 and ACP proteins were detected at approximately 38.2 kDa and 16.0 kDa, respectively.

**Figure S16.**
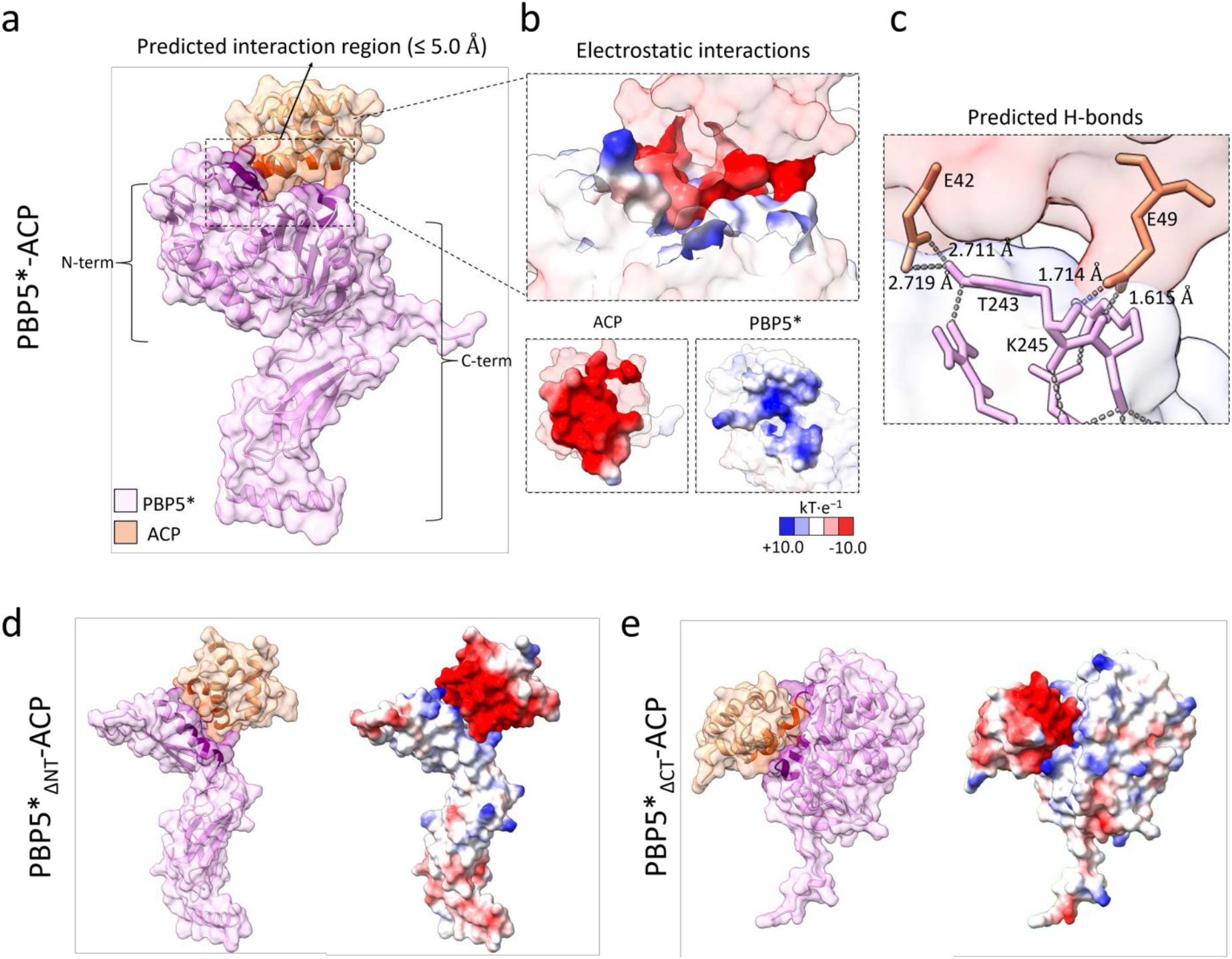
In silico modeling and docking prediction of the interaction between PBP5 and ACP. In silico docking analysis was performed using HADDOCK (ver. 2.4) to predict the interaction between ACP and PBP5 variants lacking the signal peptide. PBP5* indicates PBP5 without its signal peptide sequence. **a–c** Predicted interaction model between full-length PBP5* and ACP. PBP5* and ACP are shown in pink and orange, respectively. **A** Overall docking model of the PBP5*–ACP complex rendered using ChimeraX. The predicted interaction region within 5.0 Å is indicated. **b** Putative electrostatic interaction surface at the predicted PBP5*–ACP interface. **c** Predicted hydrogen bonds between PBP5* and ACP at the interaction interface. **d** Predicted docking model of ACP with PBP5*_ΔNT_. **e** Predicted docking model of ACP with PBP5*_ΔCT_. PBP5*_ΔNT_ and PBP5*_ΔCT_ indicate signal peptide–deleted PBP5 variants lacking the N-terminal and C-terminal regions, respectively.

**Figure S17.**
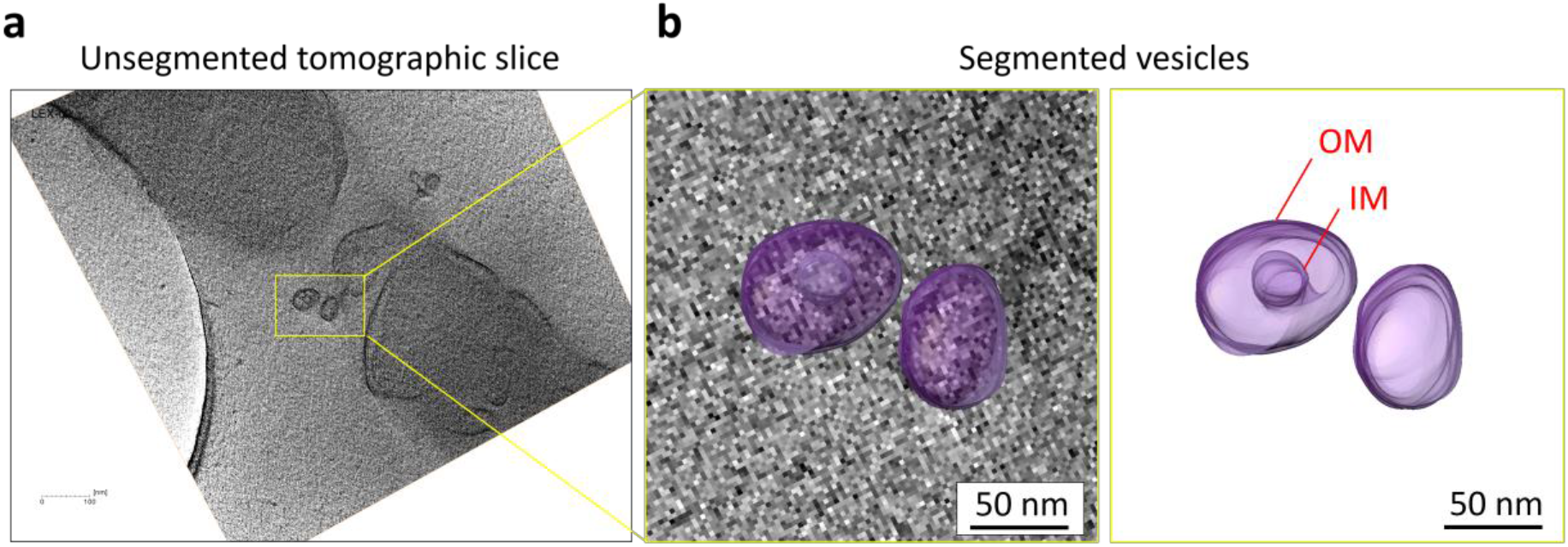
Cryo-electron tomographic visualization and 3D rendering of vesicles under LEX stress. **a** Representative cryo-electron tomogram of *S.* Typhimurium under LEX stress. **b** Magnified view and 3D rendering of the vesicles within the yellow box in a. The vesicles were selectively segmented and rendered in purple, with the OM and IM indicated. Scale bars, 50 nm (b).

**Figure S18.**
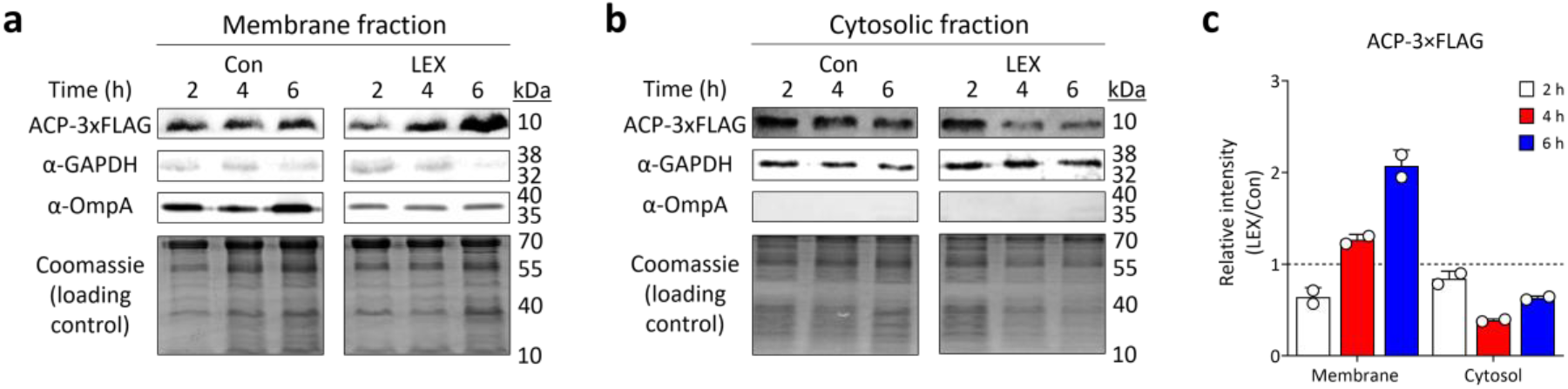
Quantification of ACP in membrane and cytosolic fractions in WT strain under control and LEX stress conditions. **a, b** Western blot analyses of ACP-3×FLAG in membrane and cytosolic fractions under control and LEX stress conditions. GAPDH and OmpA were used as cytosolic and membrane markers, respectively, and Coomassie staining was used as a loading control. **c** Quantification of ACP-3×FLAG levels in membrane and cytosolic fractions.

**Figure S19.**
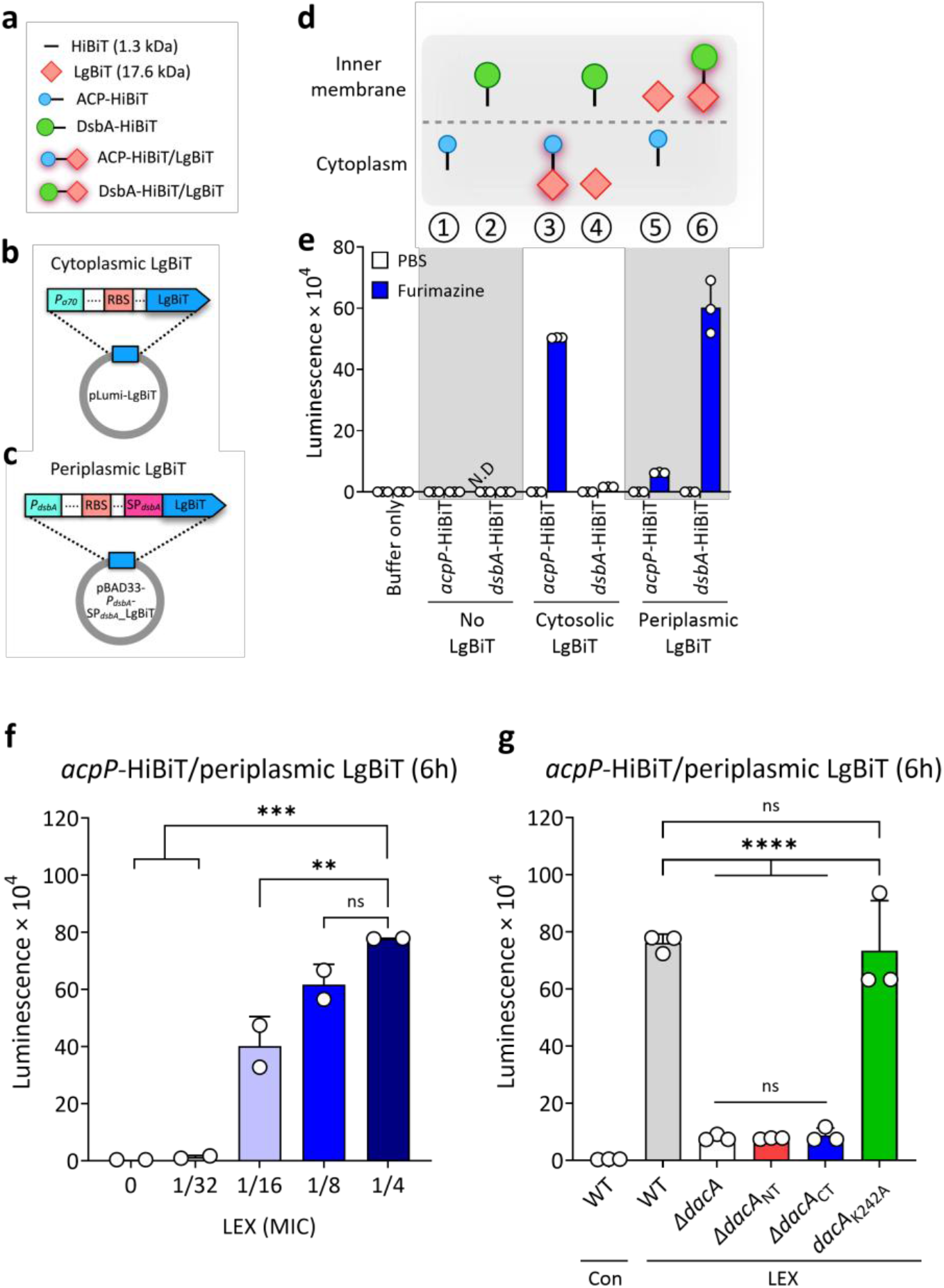
Establishment of the HiBiT–LgBiT system for monitoring ACP localization. **a** Schematic overview of the HiBiT–LgBiT system. HiBiT is a small peptide tag of approximately 1.3 kDa, whereas LgBiT is a 17.6-kDa NanoLuc. ACP-HiBiT was used as a cytoplasmic HiBiT-fused protein, and DsbA-HiBiT was used as a periplasmic control. LgBiT was expressed either in the cytoplasm or periplasm to assess compartment-specific HiBiT–LgBiT. **b** Plasmid construct for cytoplasmic expression of LgBiT. **c** Plasmid construct for periplasmic expression of LgBiT using the DsbA signal peptide. **d** Luminescence signals generated by HiBiT–LgBiT after addition of Nano-Glo substrate with furimazine. **e** Luminescence signals generated by HiBiT–LgBiT after addition of Nano-Glo substrate with furimazine. Luminescence was measured using a luminometer. *N.D, not detected. **f** LEX dose-dependent detection of periplasmic ACP using the *acpP*-HiBiT/periplasmic LgBiT system. **g** Periplasmic ACP levels in WT, *AdacA*, and *dacA* variant strains under control or LEX stress conditions, measured using the *acpP*-HiBiT/periplasmic LgBiT system. Statistical comparisons were performed using one-way ANOVA with Tukey’s multiple comparison test (**P < 0.01, ***P < 0.001, and ****P < 0.0001; ns, not significant).

**Figure S20.**
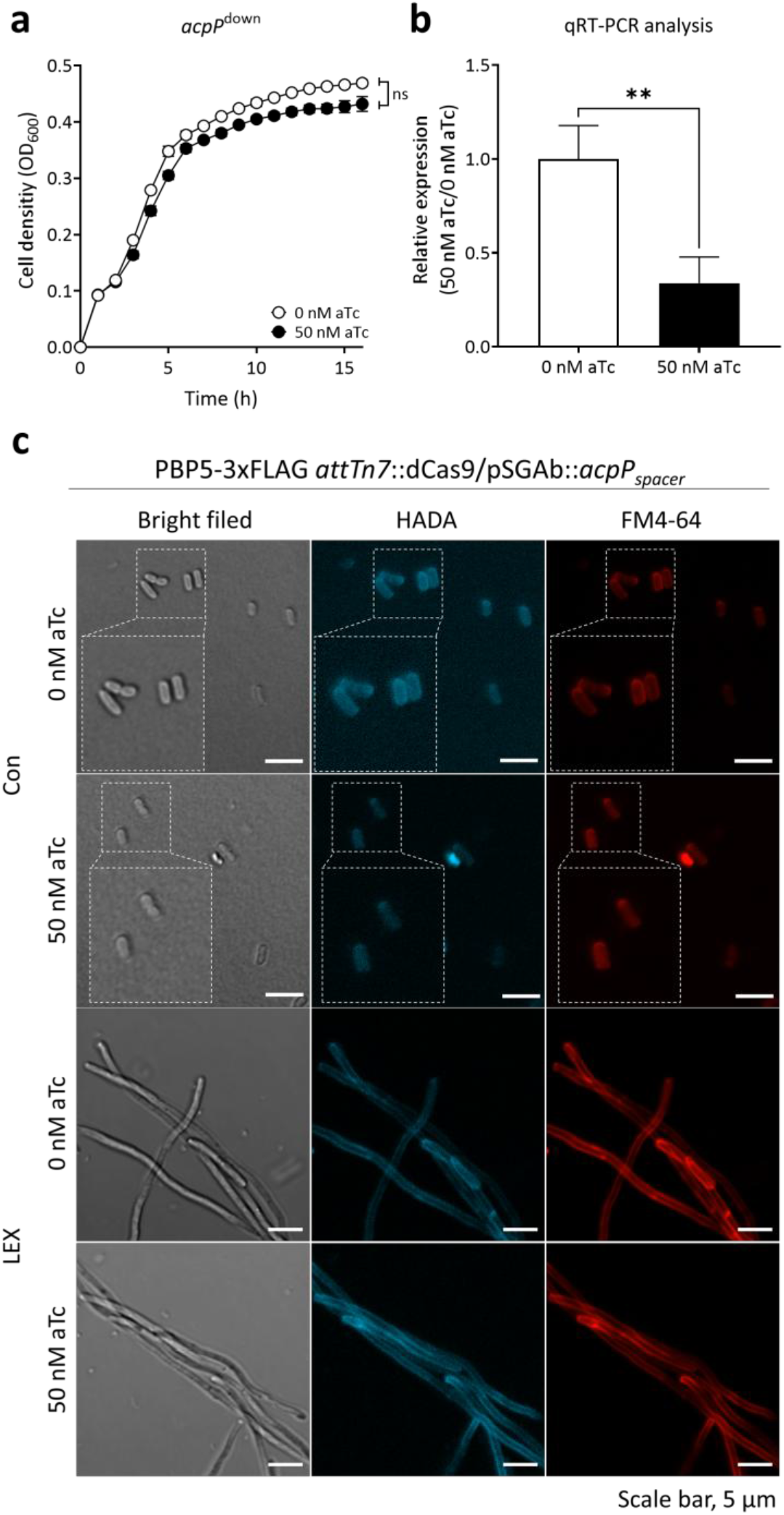
Validation and microscopic analyses of *acpP*^down^ strain under control and LEX stress conditions. The *acpP*^down^ strain, carrying chromosomally integrated *attTn7*::dCas9 and the pSGAb::*acpP*_spacer_, was cultured in LB broth under static conditions at 37°C and treated with 50 nM aTc to induce dCas9-dependent repression of *acpP* gene. **a** Growth curve of the *acpP*^down^ strain cultured in LB with or without 50 nM aTc. **b** Relative expression of *acpP* gene measured by qRT-PCR. **c** Microscopic observation of *acpP*^down^ strain cells under control and LEX stress conditions at 6 h. Cells were stained with HADA to visualize DD-transpeptidase activity and with FM4-64 to visualize the OM. Statistical comparisons were performed using one-way ANOVA with Tukey’s multiple comparison test **P < 0.01; ns, not significant).

**Figure S21.**
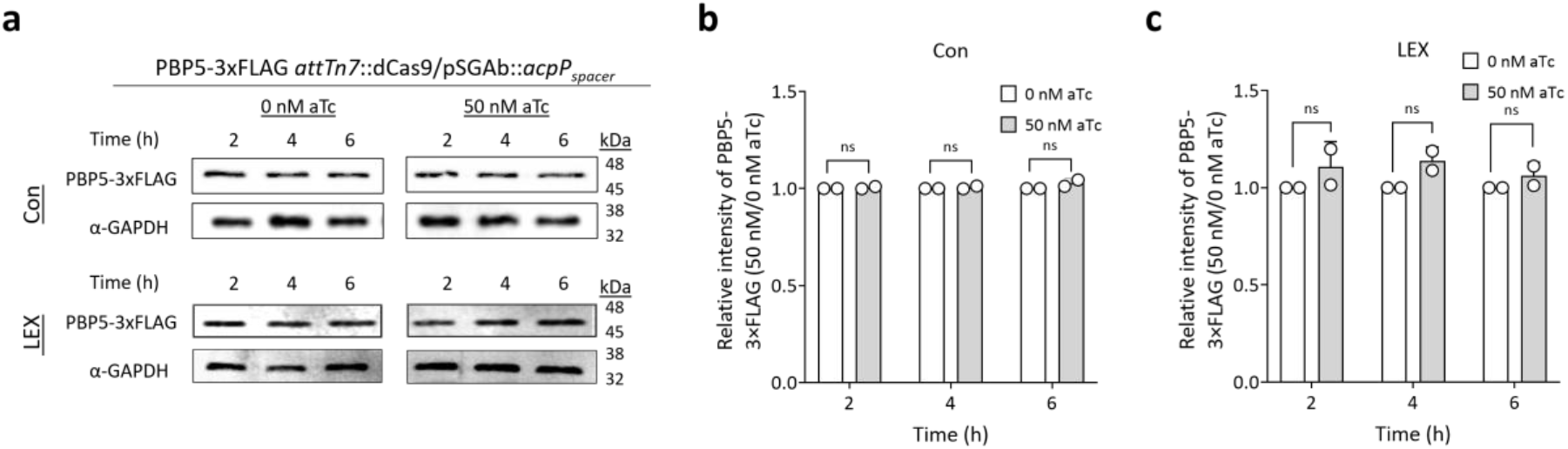
*acpP* knockdown on PBP5 protein levels under control and LEX stress conditions. **a** Representative Western blots of PBP5–3×FLAG in the *acpP* knockdown strain carrying chromosomally 3×FLAG-tagged PBP5, grown at 37°C with or without LEX in the presence of 0 or 50 nM aTc for 2, 4, and 6 h. GAPDH was used as a loading control. Crude extracts were normalized to 100 µg/mL total protein**. b, c** Quantification of relative PBP5–3×FLAG band intensities under control and LEX stress conditions, normalized to the corresponding 0 nM aTc condition at each time point. ns, not significant.

**Figure S22.**
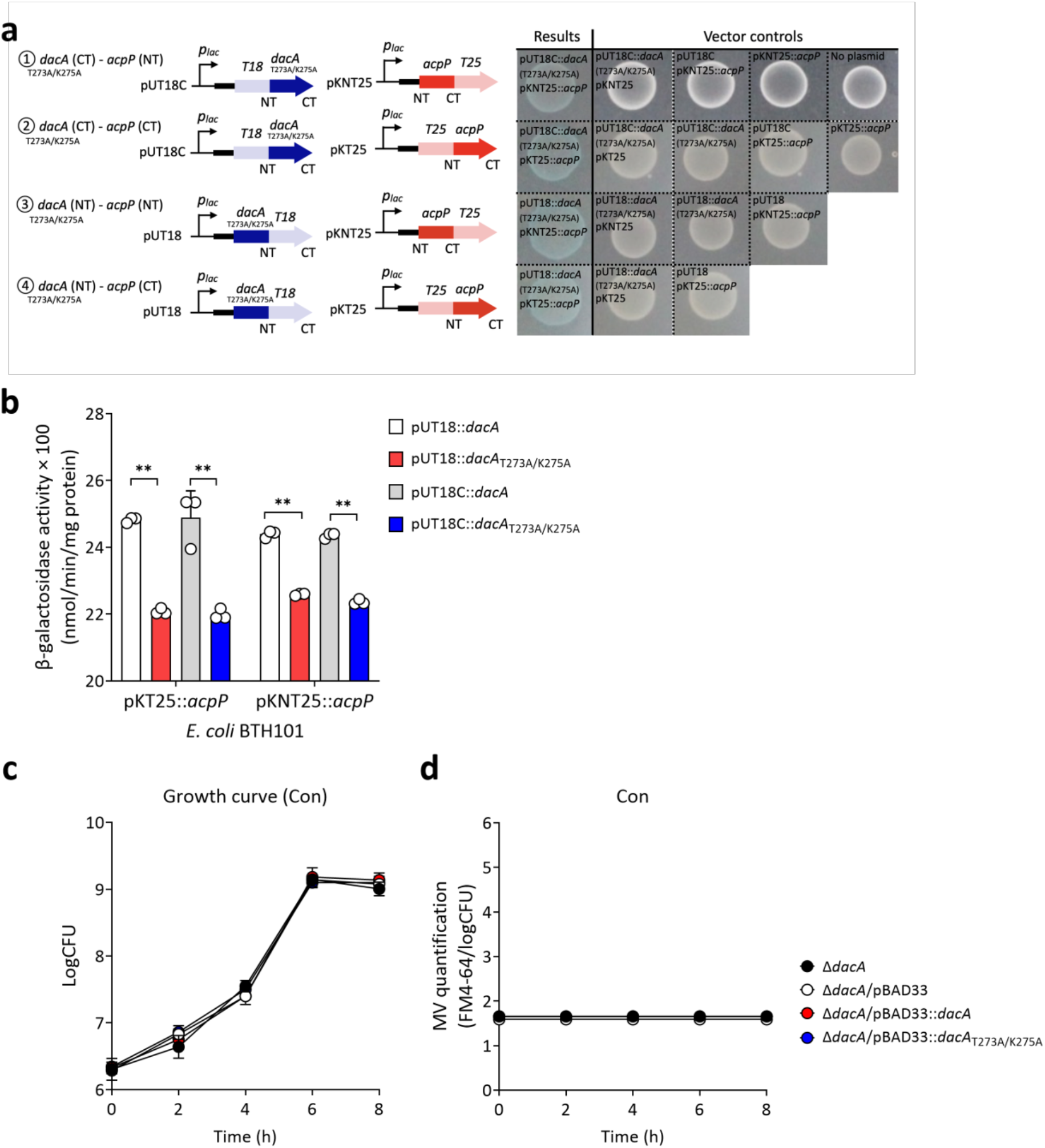
Reduced PBP5–ACP interaction in the PBP5 interface mutant and growth and MV production under control conditions. **a** Bacterial adenylate cyclase two-hybrid (BACTH) assays examining the interaction between the PBP5* interface mutant (T273A/K275A) and ACP in *E. coli* BTH101. Four combinations of N- and C-terminal T18- and T25-fusion constructs were tested, with the corresponding vector controls. **b** Quantitative β-galactosidase activity measurements comparing the interactions of WT PBP5* and the PBP5* interface mutant (T273A/K275A) with ACP using the indicated BACTH fusion constructs. **c**, **d** Growth curves and MV production of the *AdacA* strain carrying an empty pBAD33 vector, WT *dacA*, or the *dacA*_T273A/K275A_ interface mutant expressed from pBAD33 under the native dacA promoter, together with the *AdacA* strain, under control conditions (LB broth without LEX) at 37°C. MV production was quantified using FM4-64 fluorescence and normalized to viable cell counts (log CFU). Data are presented as mean ± SD from three independent biological replicates. Statistical significance was determined using a two-tailed Student’s t-test (**P < 0.01).

**Figure S23.**
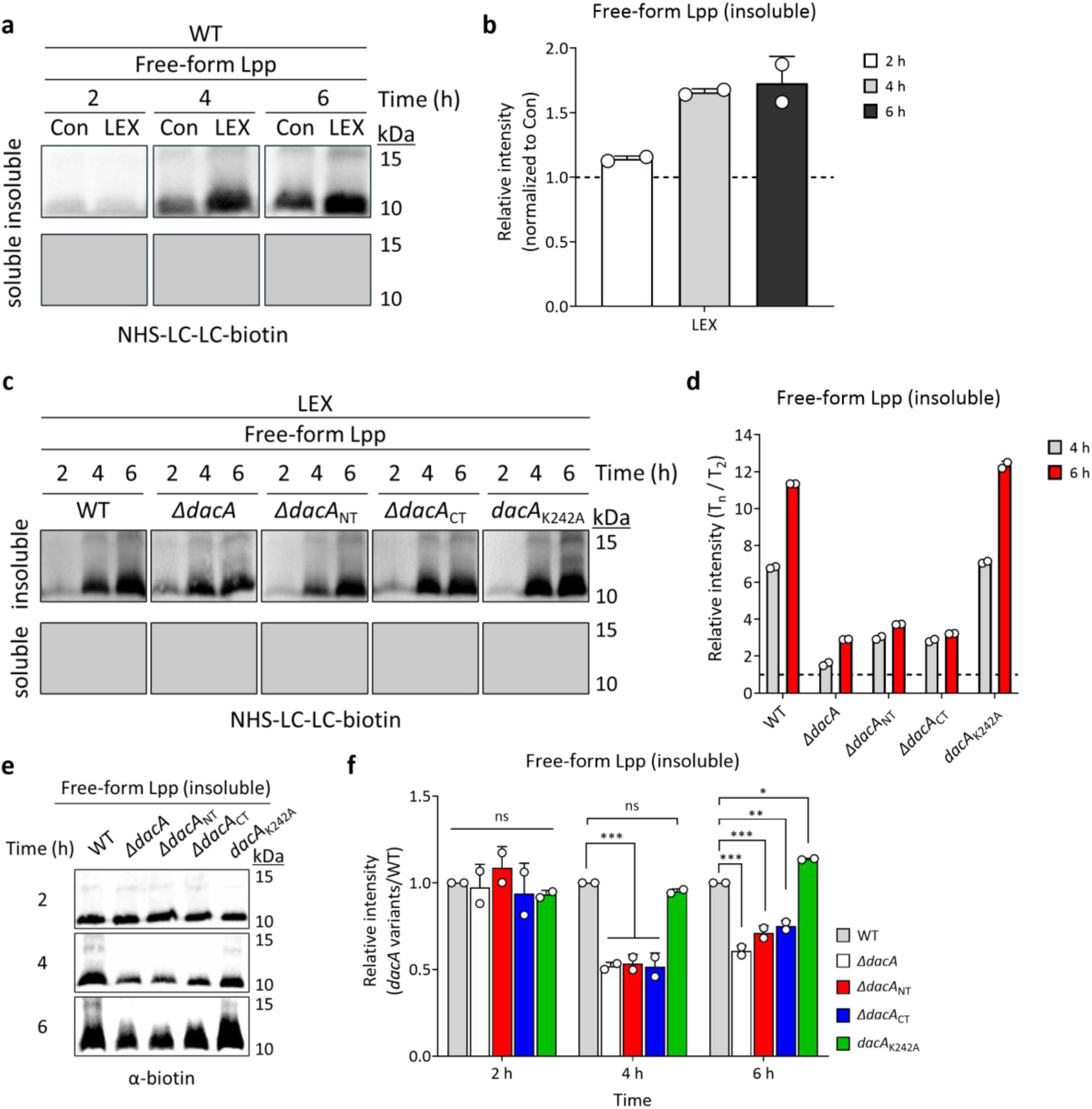
Measurement of free-Lpp accumulation in WT and *dacA* variants under control and LEX stress conditions. Free-Lpp levels were analyzed by NHS-LC-LC-biotin labeling. WT and *dacA* variant cells were cultured in LB broth with or without LEX under static conditions at 37°C and harvested at the indicated time points. Cells were labeled with NHS-LC-LC-biotin, and biotin-labeled proteins in the soluble and insoluble fractions were detected by Western blotting using HRP-conjugated streptavidin**. a, b** Time-dependent accumulation of free-Lpp in WT cells under control and LEX stress conditions at 2, 4, and 6 h. **c, d** Time-dependent changes in free-Lpp levels in WT and *dacA* variants under control and LEX stress conditions. **e, f** Comparison of free-Lpp levels among WT and *dacA* variants at the indicated time points. Statistical comparisons were performed using one-way ANOVA with Tukey’s multiple comparison test (*P < 0.05, **P < 0.01, and ***P < 0.001; ns, not significant).

**Figure S24.**
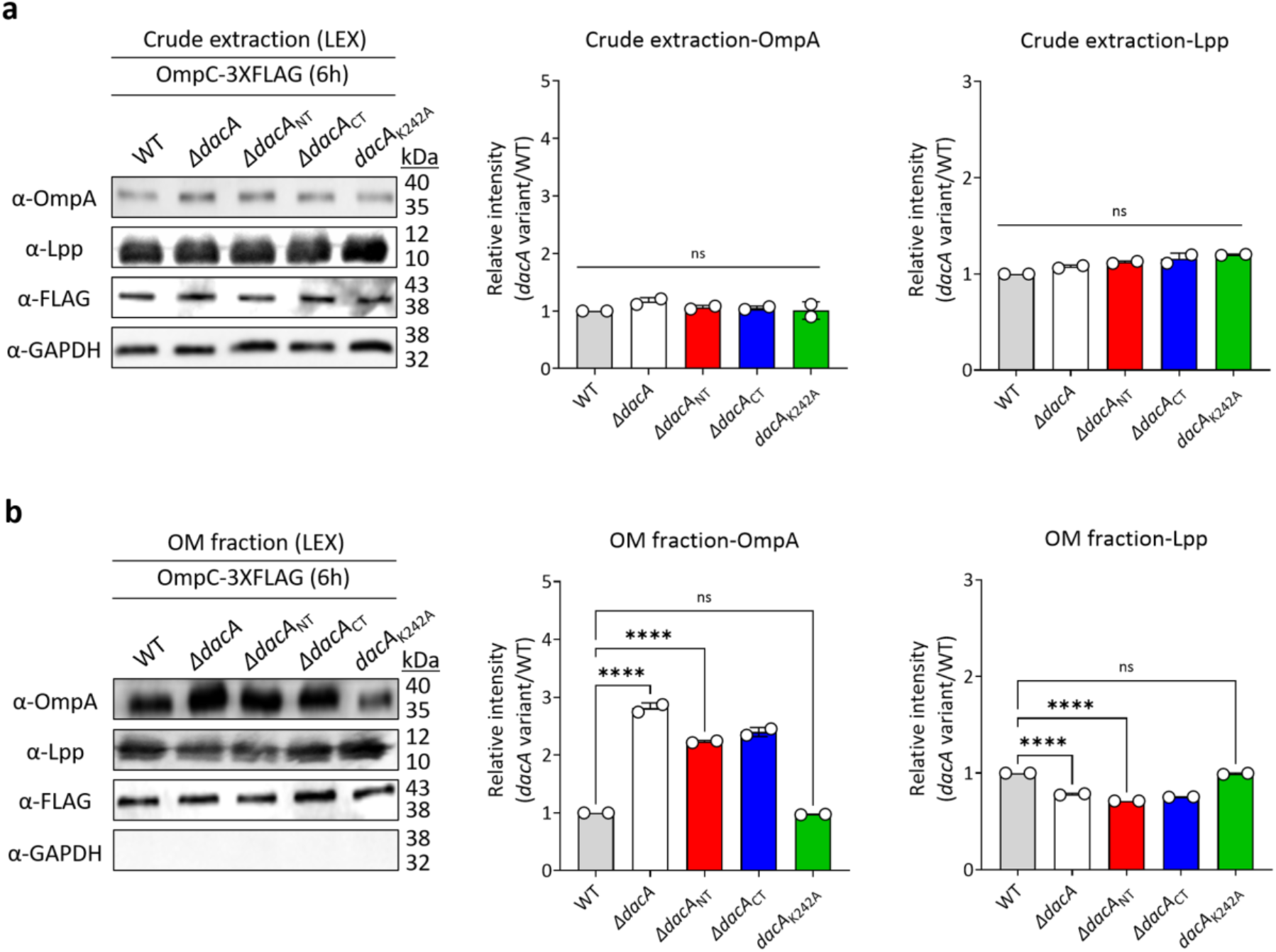
PBP5-dependent periplasmic crowding–induced OM–PG tethering defects. **g–i** Western blot analyses of OmpA and Lpp levels in crude extracts and OM fractions of WT and *dacA* variant strains grown under LEX stress for 6 h. Total OmpA and Lpp levels in crude extracts were comparable among strains, whereas their OM-associated pools was altered depending on the *dacA* variant. OmpC-3×FLAG was used as an OM-associated marker, and GAPDH was used as a cytosolic control. Individual points indicate biological replicates, and bars indicate mean values. Statistical comparisons were performed using one-way ANOVA with Tukey’s multiple comparison test (****P < 0.0001; ns, not significant).

**Figure S25.**
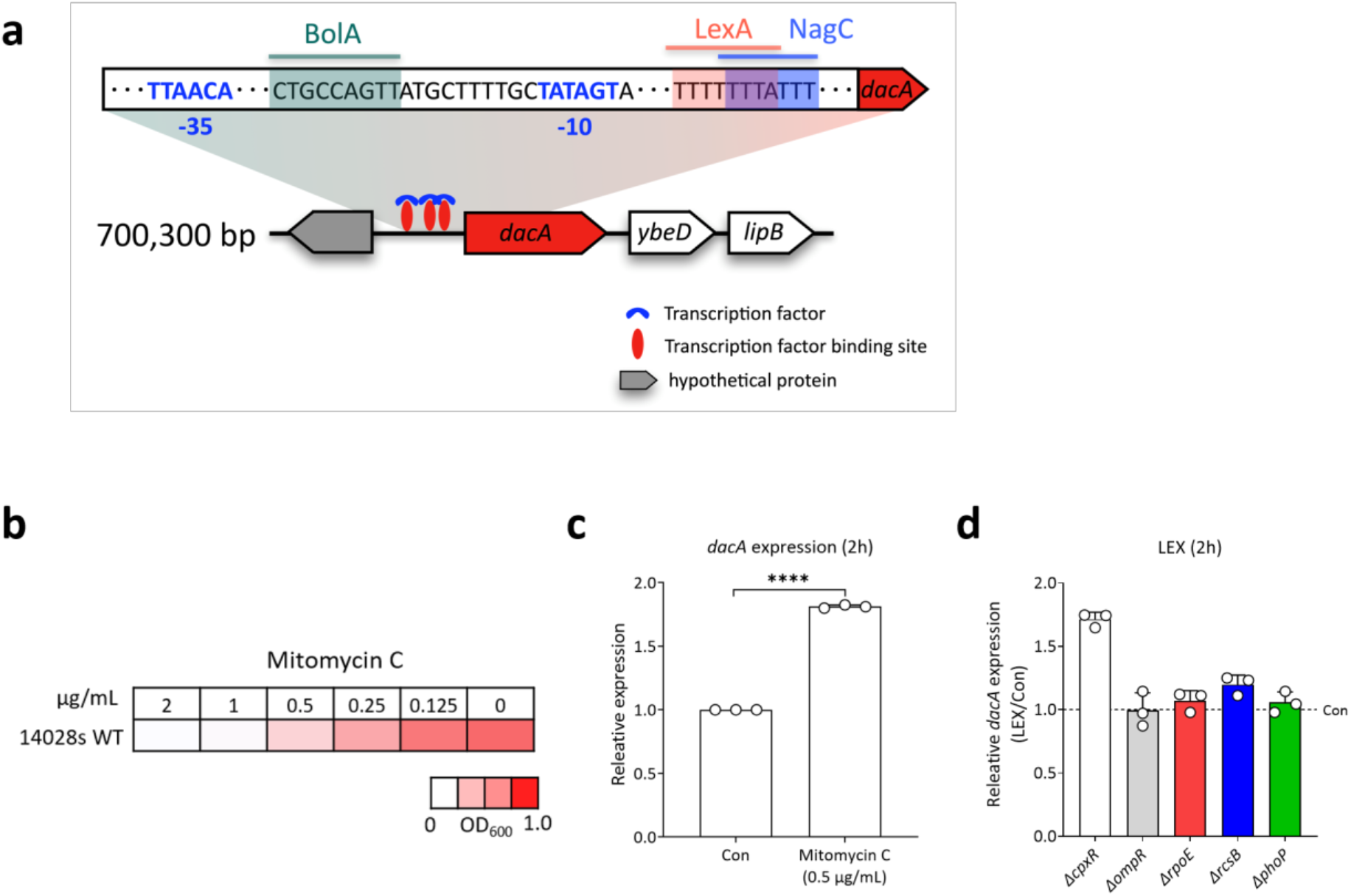
Regulatory analyses of LEX-induced *dacA* expression in *S.* Typhimurium. **a** Schematic representation of the genomic context and predicted regulatory region upstream of *dacA*. Putative transcription-factor-binding sites, including BolA, LexA, and NagC, are indicated together with the predicted −35 and −10 promoter elements. **b** MIC test of WT strain under mitomycin C stress. **c** Relative expression levels of *dacA* gene in WT cells after 2 h of mitomycin C treatment (0.5 µg/mL), normalized to the untreated control. **d** Relative *dacA* expression in *AcpxR, AompR, ArpoE, ArcsB,* and *AphoP* after 2 h of LEX treatment, normalized to the corresponding untreated control. Statistical comparisons were performed using one-way ANOVA with Tukey’s multiple comparison test (****P < 0.0001).

**Figure S26.**
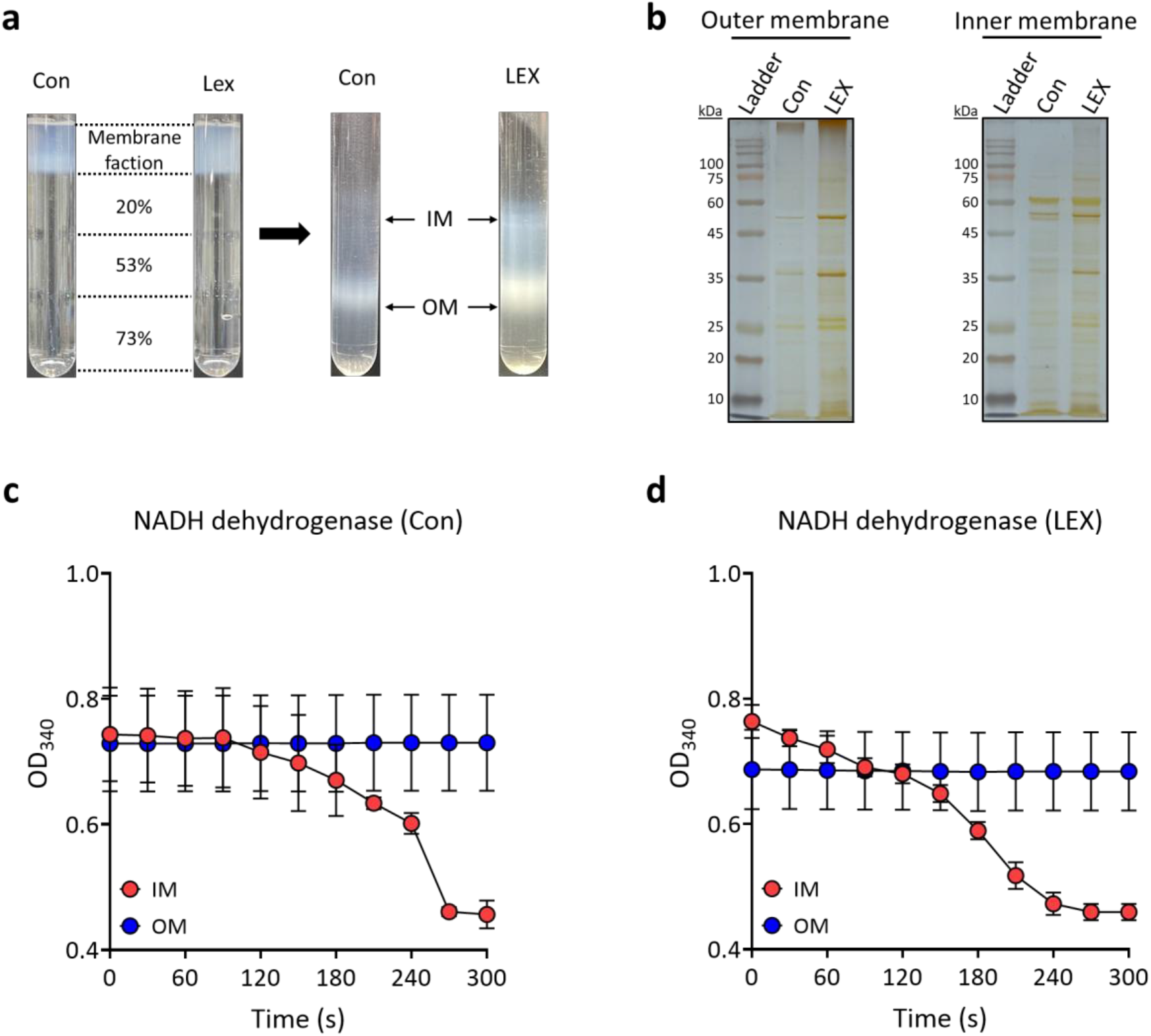
Separation of OM and IM fractions from *S.* Typhimurium cells. OM and IM fractions were separated by sucrose density gradient centrifugation using 20%, 53%, and 73% sucrose layers. **a** Representative images of sucrose density gradients after membrane fractionation. **b** Protein profiles of the recovered OM and IM fractions were visualized by SDS-PAGE (12%) followed by silver staining. **c, d** NADH dehydrogenase activity assays of the recovered OM and IM fractions to validate membrane fraction separation.

**Figure S27.**
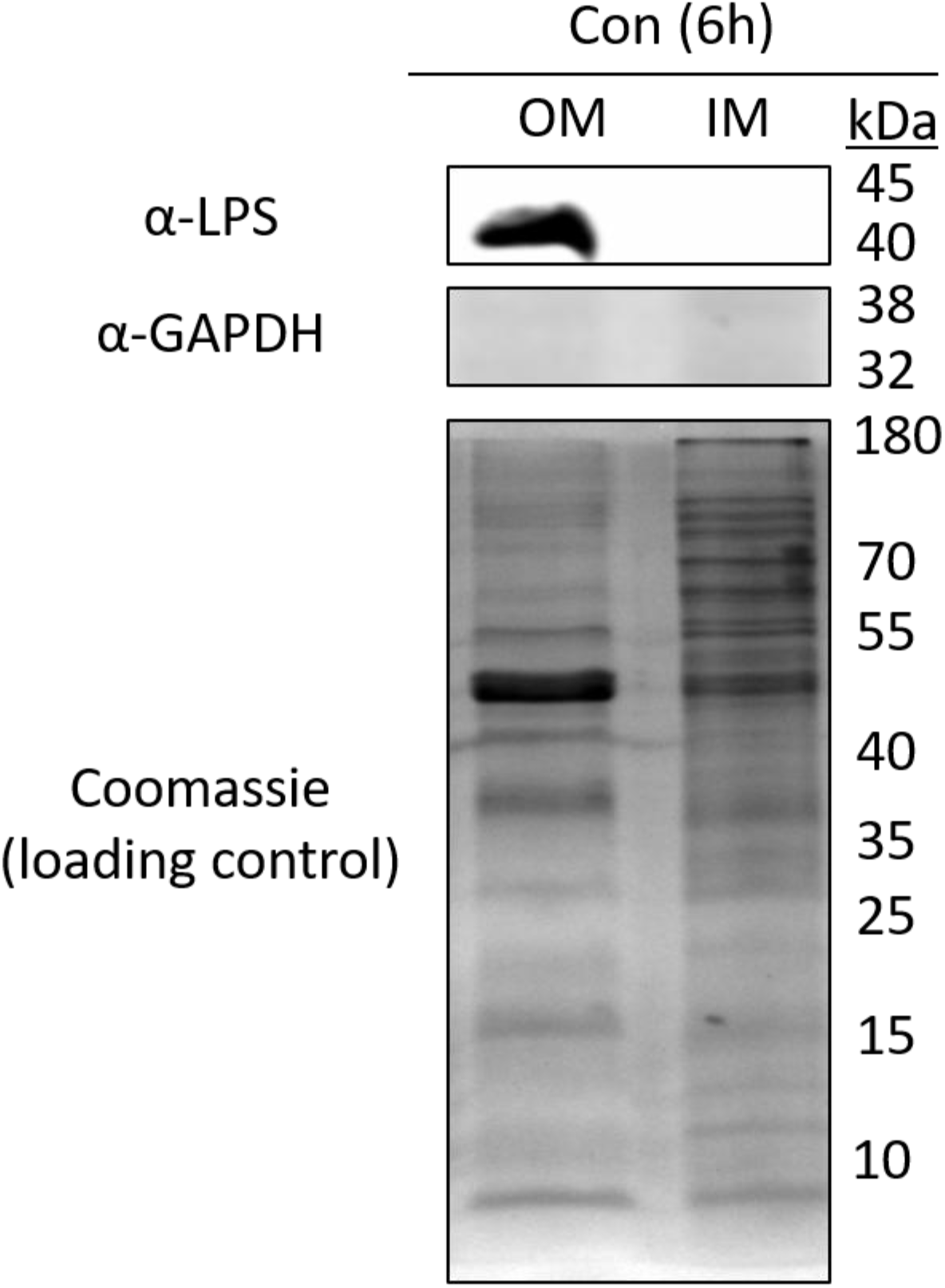
Validation of OM- and IM-enriched membrane fractions from *S.* Typhimurium cells. Fraction enrichment was assessed using LPS as an OM marker and GAPDH as a cytoplasmic marker, and equal protein loading was verified by Coomassie Brilliant Blue staining.

**Video S1. Cryo-electron tomographic visualization of the bacterial envelope under control condition.** Representative tomographic reconstruction of an untreated control cell, visualized using Amira. Th OM and IM are shown in cyan and yellow, respectively. Scale bar, 400 nm.

**Video S2-3. Cryo-electron tomographic visualization of the bacterial envelope under LEX stress.** Representative tomographic reconstruction of a LEX-treated cell, visualized using Amira. The OM, IM, and OMVs are shown in cyan, yellow, and purple, respectively. Scale bar, 200 nm.

